# Inferring Ligand-Receptor Interactions between neuronal subtypes during mouse cortical development

**DOI:** 10.1101/2024.09.02.610245

**Authors:** Rémi Mathieu, Léa Corbières, Tangra Draia-Nicolau, Annousha Govindan, Vianney Bensa, Emilie Pallesi-Pocachard, Lucas Silvagnoli, Alfonso Represa, Carlos Cardoso, Ludovic Telley, Antoine de Chevigny

## Abstract

The cerebral cortex hosts a diverse array of excitatory glutamatergic and inhibitory GABAergic neuron types, each characterized by distinct positional and synaptic connectivity patterns. However, the molecular mechanisms orchestrating this precise organization remain largely unknown. To identify ligand-receptor (LR) pairs regulating interactions and connectivity among cortical neurons during embryonic and postnatal development, we analyzed the transcriptional dynamics of all genes across major cortical neuron subtypes at 17 developmental time points using single-cell transcriptomics. From these data, we constructed a comprehensive bioinformatic atlas that inferred significant LR-mediated interactions between glutamatergic and GABAergic neurons throughout cortical maturation. This atlas not only corroborated known interactions but also enabled the discovery of novel regulators, identifying two cadherin superfamily members as key mediators of perisomatic inhibition in deep and superficial layer excitatory neurons by parvalbumin-expressing basket cells. These findings underscore the power of large-scale transcriptional profiling to unravel fundamental molecular mechanisms driving cortical circuit assembly.

## Main Text

The mammalian cerebral cortex, a six-layered structure located at the brain surface, is responsible for executing the most complex cognitive functions. To achieve these functions, it must undergo highly orchestrated and conserved developmental processes ultimately leading to the finely organized communication between two primary neuron classes: excitatory glutamatergic neurons (GlutNs) and inhibitory GABAergic neurons (GABANs). Both GlutNs and GABANs are further subdivided into numerous subtypes, a diversity primarily uncovered through single-cell RNA sequencing (scRNA-seq). Tens of cortical GlutN and GABAN subtypes have been identified ^1–5^, which (1) exhibit specific morphological, electrophysiological and transcriptomic properties, (2) preferentially localize to certain cortical layers, (3) form synaptic connections with specific pre and post-synaptic partners, preferentially targeting specific subcellular compartments and (4) play specialized roles in cortical computation. The resulting cortical arrangement and associated synaptic connectivity, involving hundreds to thousands of potential neuron pairs, is responsible for cortical function. Even a disruption in the communication between just one pair of neuron subtypes at a particular developmental stage might lead to cortical dysfunction and subsequent neurodevelopmental disorders ^6^. Therefore, understanding how relative physical positions and synaptic connectivity rules between the hundreds of potential neuron pairs are established and regulated is crucial for advancing developmental and clinical neuroscience.

Over the past decade, substantial efforts have been directed toward elucidating the spatial relationships and synaptic connectivity rules among cortical neuronal subtypes. In particular, a wealth of groundbreaking studies has provided high-resolution insights into the laminar organization and connectivity patterns of cortical GlutNs and GABANs in the adult mouse brain. These studies have demonstrated the remarkable specificity, stereotypy, and evolutionary conservation of this intricate cellular and subcellular architecture across mammals ^7–10^. Despite this progress, the molecular mechanisms underlying corticogenesis that culminate in such a finely orchestrated final organization remain largely elusive. Only a handful of pioneering studies have begun to uncover specific molecules driving the cellular interactions and processes shaping cortical structure during development ^11–13^.

It is likely that neuron-neuron communications via ligand-receptor (LR) interactions play a key role in general ^11–13^, as these interactions are critical for the development of many tissues. This hypothesis is supported by several studies suggesting that GlutNs may non-cell autonomously influence the recruitment of specific GABAN subtypes. Indeed, GlutN subtypes settle into their final positions earlier than the synchronically generated GABANs ^14^, and altering specific GlutN subtype identities during development modifies the allocation and synaptic connectivity of corresponding GABAN subtypes ^15–17^.

In this study, we leverage both newly generated and previously published scRNA-seq datasets from mice to elucidate the molecular mechanisms underlying subtype-specific neuron communication during cortical development. First, we tracked gene expression dynamics across all GlutN and GABAN subtypes throughout corticogenesis. Then, we constructed a comprehensive atlas to infer LR interactions between these neuronal subtypes. Finally, we validated the utility of our atlas by (1) confirming known LR interactions, (2) bridging gaps in existing knowledge by identifying receptors involved in well-characterized neuron-neuron interactions, and (3) uncovering novel LRs within the cadherin family that are crucial for perisomatic inhibition in deep versus superficial cortical layers. The data can be explored through an interactive <u>Shiny App</u>, currently accessible via the instructions provided in the README file, which will be available as a web-based interface upon publication of the paper.

### Comprehensive single neuron transcriptomics atlas covering mouse somatosensory cortex development

To infer LR-mediated cell-cell interactions during mouse neurodevelopment, our strategy was to cross-reference single-cell RNA sequencing (scRNA-seq) transcriptomic data from all cortical neurons across all multiple stages with publicly-known LR information. The purpose of scRNA-seq was to both identify neuronal subtypes and predict LR expression over time.

First, we generated a comprehensive scRNA-seq transcriptomic dataset of all cortical neurons throughout somatosensory corticogenesis. While prenatal and adult stages are well-documented in the mouse, the P0-P30 period remains underexplored, despite being crucial for synaptogenesis. We conducted scRNA-seq and snRNA-seq at six key stages within this timeframe: P0-P2 (radial migration and laminar allocation of GABANs ^18,19^), P5-P8 (programmed cell death of GlutNs and GABANs ^20–22)^, and P16-P30 (circuit refinement and synaptogenesis completion). We integrated to this dataset previously published scRNA-seq data covering earlier embryonic (E11.5-E18.5) and adult stages (P53-P102) ^3,23–28^, including ganglionic eminences (GEs) to capture early transcriptional development of future cortical GABANs ^25–28^ (see Methods). In total, our analysis spans 17 time points from E11.5 to adulthood (Fig. 1A).

**Figure 1.**
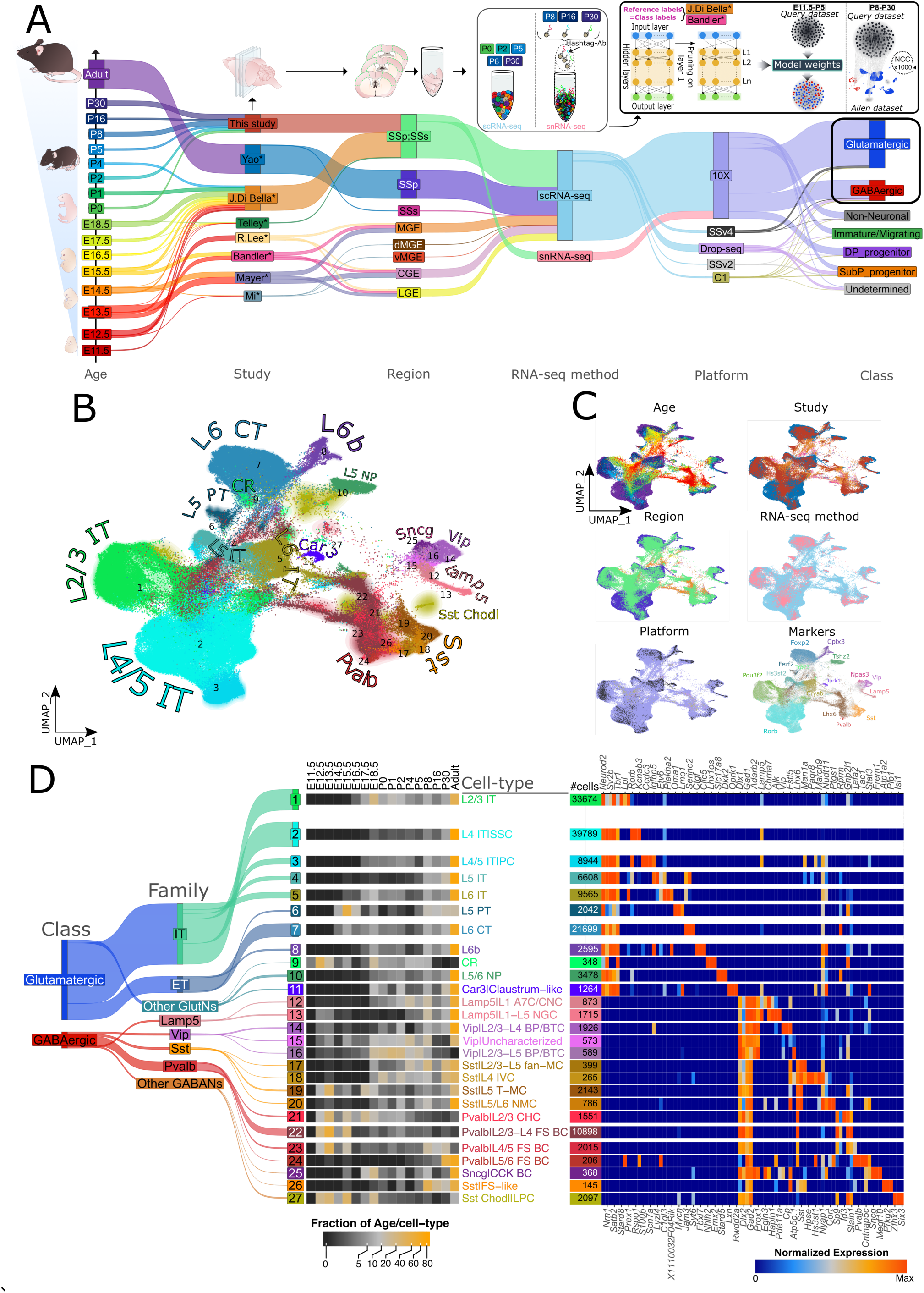
Comprehensive neuronal transcriptomic atlas of mouse somatosensory cortex development. (A) Sankey diagram depicting the experimental paradigm for data collections and integration of published datasets. Yao*: Yao et al., 2021; J. Di Bella*: J. Di Bella et al., 2021, Telley*: Telley et al., 2019; R.Lee*: R.Lee et al., 2022; Bandler: Bandler et al., 2021; Mayer*: Mayer et al., 2018; Mi*: Mi et al., 2018; SSp: primary somatosensory cortex; SSs: supplemental somatosensory cortex; MGE: medial ganglionic eminence; dMGE: dorsal MGE; vMGE: ventral MGE; CGE: caudal ganglionic eminence; LGE: lateral ganglionic eminence; scRNA-seq: single-cell RNA sequencing; snRNA-seq; single-nuclei RNA sequencing; 10X: 10X Genomics; SSv4: Smart-seq version 4; SSv2: Smart-seq version 2; C1: Fluidigm C1. (B) UMAP visualization of post-mitotic neurons from 17 time points after integration. Cells are colored by *cell-type* assignment. (C) UMAP visualization is colored by age, study, region, RNA-seq method, sequencing platform, and gene markers. (D) Left: Sankey plot representing the different hierarchical levels of *cell-type* assignment; middle: heatmap representing the fraction of each cell type per age; right: heatmap representing some cell type-specific genes. “Normalized expression” represents the 25% trimmed mean log2(CPM + 1) expression normalized by row. See also Figs. S1-S4.

After stringent quality control and filtering (Fig. S1A-B and Methods), we first aimed to identify all postmitotic neuronal cell types and track their transcriptional dynamics over cortical development, from E11.5 to adulthood. For final cell type nomenclature, we chose as reference the most comprehensive resource of transcriptomic cell-types in the somatosensory cortex that existed before December 2023*, i.e.*, Yao et al. (2021) ^3^ (from now on referred to as AllenRef21), corresponding to the adult stage (P53-P102) (Fig. S1A). Assigning cellular identities in early age datasets proved difficult with simple methods given that transcriptional signatures of specific cell-types evolve drastically during development ^23,26^. To overcome this difficulty, we developed a hierarchical pipeline assigning identities at increasing resolution levels, similar to the method used by the Allen Institute ^3^ (Fig. S1A).

First, we assigned cell identities at a broad, low resolution level referred to as the *class* level (Glutamatergic, GABAergic, Non-Neuronal, Immature/Migrating, dorsal pallium progenitor and subpallium progenitor) ^3^. For *classes* of embryonic and neonatal datasets (E11.5 to P5), we took as references the datasets of Di Bella et al. ^24^ (largest published perinatal dataset for somatosensory cortex) and Bandler et al. ^27^ (corresponding to early GE GABANs destined to the cortex). For P8 to P30, because cells were transcriptionally more similar to the adult reference, we took directly as the reference the adult dataset AllenRef21 ^3^ (Fig. S1A). To test our hypothesis that a ligand-receptor (LR) code between GlutNs and GABANs plays a key role in the establishment of cortical microcircuits, we focused our downstream analyses on post-mitotic GlutN and GABAN cells (Fig. 1A, B). In total, 182,084 post-mitotic neurons were further analyzed. To analyze biological variation in gene expression and get rid of technical variation due to different studies, RNA-seq methods (scRNA-seq and snRNA-seq) and sequencing platforms (Fig. 1C), we integrated all data with the Seurat SCT workflow (Fig. S1C). The resulting integrated dataset was split into *classes*, i.e., GlutNs and GABANs. For each *class*, cell identities were further delineated at increasing resolutions; they were determined first at the *subclass* level, and then at the *supertype* level in each subclass, as defined in AllenRef21 ^3^. A limitation of the *supertype* level is that it does not correspond to known morpho-functionally identified cortical neuronal subtypes. To identify biologically meaningful cell types in our analysis we leveraged the fact that Yao et al. provided the correspondence between *supertypes* from scRNA-seq studies and morpho-electro-connectomic types (*mec-types*) from patch-seq studies ^1–3,29^ (Fig. S2, S3). Pooling *supertypes* belonging to the same *mec-type* allowed us to reach a biologically meaningful *cell-type* annotation (Fig. S4). Ultimately, our analysis led to the identification of 159 290 cells, representing 27 *cell-types* (Fig. 1D) including 11 GlutN *cell-types* grouped in 3 *families* – intratelencephalic (IT), extratelencephalic (ET), Other GlutNs - and 16 GABAN *cell-types* grouped in 5 *families* - Lamp5, Sst, Pvalb, Vip and Other GABANs - (Fig. 1B-D, Fig. S4).

In the UMAP embedding of all neurons from 17 ages, each GlutN and GABAN *cell-type* exhibited a largely continuous temporal gradient of transcriptional variation across corticogenesis, with gene expression profiles of individual cells correlating closely with their differentiation states / mouse developmental age (Fig. 1B, C). the integration and annotation efforts presented here establish a valuable new reference for the developmental neurobiology community.

### Temporal transcriptional dynamics of cortical neuron subtypes

To track the temporal maturation of each neuronal *cell-type* over migration, programmed cell death, dendritic/axonal development and synaptogenesis in a continuous rather than discrete way, we used an approach in which single cells were ordered on a linear path on the basis of their transcriptional profile ^30^ (see Methods) (Fig. S5A, Fig. S6A-B). This “pseudo-maturation” axis correlated well with the actual age of the cells (Fig. S5A, Fig. S6A-B), indicating gradual variation in gene expression over neuronal differentiation.

For each *cell-type*, we found genes showing significant variation along the “pseudo-maturation” axis and used them to display temporal gene dynamics in 6 consecutive waves (Fig. S5A, Fig. S6A-B). During early development (E11.5-E18.5), most *cell-types* showed transcriptional signatures related with cell-intrinsic properties (Fig. S5B, Fig. S6C), while later on (neonatal to P30), genetic programs related with cell-cell and cell-environment interactions predominated (Fig. S5B, Fig. S6C) ^31,32^. These findings reveal the molecular transitions through which each neocortical cell type shifts from intrinsic programs to extrinsic, interaction-driven programs essential for their integration into the developing cortical network.

### Spatial transcriptional gradients at given developmental ages

In adults, GlutN ITs exhibit a spatial gradient of transcriptional variation along the cortical sheet ^3,33^. We investigated the existence of such spatial transcriptomic gradients for each cell *family* at each age analyzed between E18.5 to P30. The correspondence between our datasets and published Patch-seq ^1,2^ and MERFISH ^33^ datasets allowed us to determine the median adult laminar distribution for each *cell-type* (Fig. S2 & S3). Using a similar approach as for pseudo-maturation, for each *family* that displayed spatial organization along the cortical sheet and for each age, we fitted a curve along UMAP continua, and called this axis the “pseudo-layer” score.

The spatial transcriptional gradient described for the IT *family* in adults ^33^ was also found at all earlier ages examined, from E18.5 to P30 (Fig. S7A). Strikingly, we also identified clear spatial transcriptional continua for ET (Fig. S7C), Sst (Fig. S8A), Pvalb (Fig. S8C), Vip (Fig. S9A) and Lamp5 (Fig. S9C) *families* at all ages examined. For each *family* and at every age, we identified genes with significant variation along pseudo-layers and used them to discretize gene dynamics in 6 spatial transcriptional waves. In GlutNs of neonatal mice (E18.5 to P5), the first waves, which corresponded to upper layer GlutNs (UL), displayed genetic signatures of immaturity such as intrinsic properties (Fig. S7, B and D); this was not the case for late waves, which correspond to deep layers (DL). This is consistent with UL GlutNs being born later than DL GlutNs - due to cortical inside-out patterning - and therefore maturing later ^23^. In contrast, for GABANs, environment-sensing programs were already active by E18.5 (Fig. S8, B and D), supporting the hypothesis that these neurons mature later than the earlier-settled GlutNs and rely on signals from them to reach their final positions and establish proper connections ^15–17,34^.

### Spatiotemporal transcriptional landscapes

On the basis of the pseudo-maturation and pseudo-layer scores, cells of each *family* were embedded within 2D graphs, so as to generate spatiotemporal transcriptional landscapes of gene expression (Fig. 2A-D and S10). For any gene queried in a given *family*, the 2D transcriptional map shows temporal dynamics in the x axis and spatial dynamics in the y axis (Fig. 2B-D). This allowed us to confirm the spatiotemporal expression of genes known to be involved in cortical layer patterning (Cux1, Fezf2, Reln, Fig. 2B-C) or in neuropeptidergic system maturation in specific neuronal *families* (Fig. 2D).

**Figure 2.**
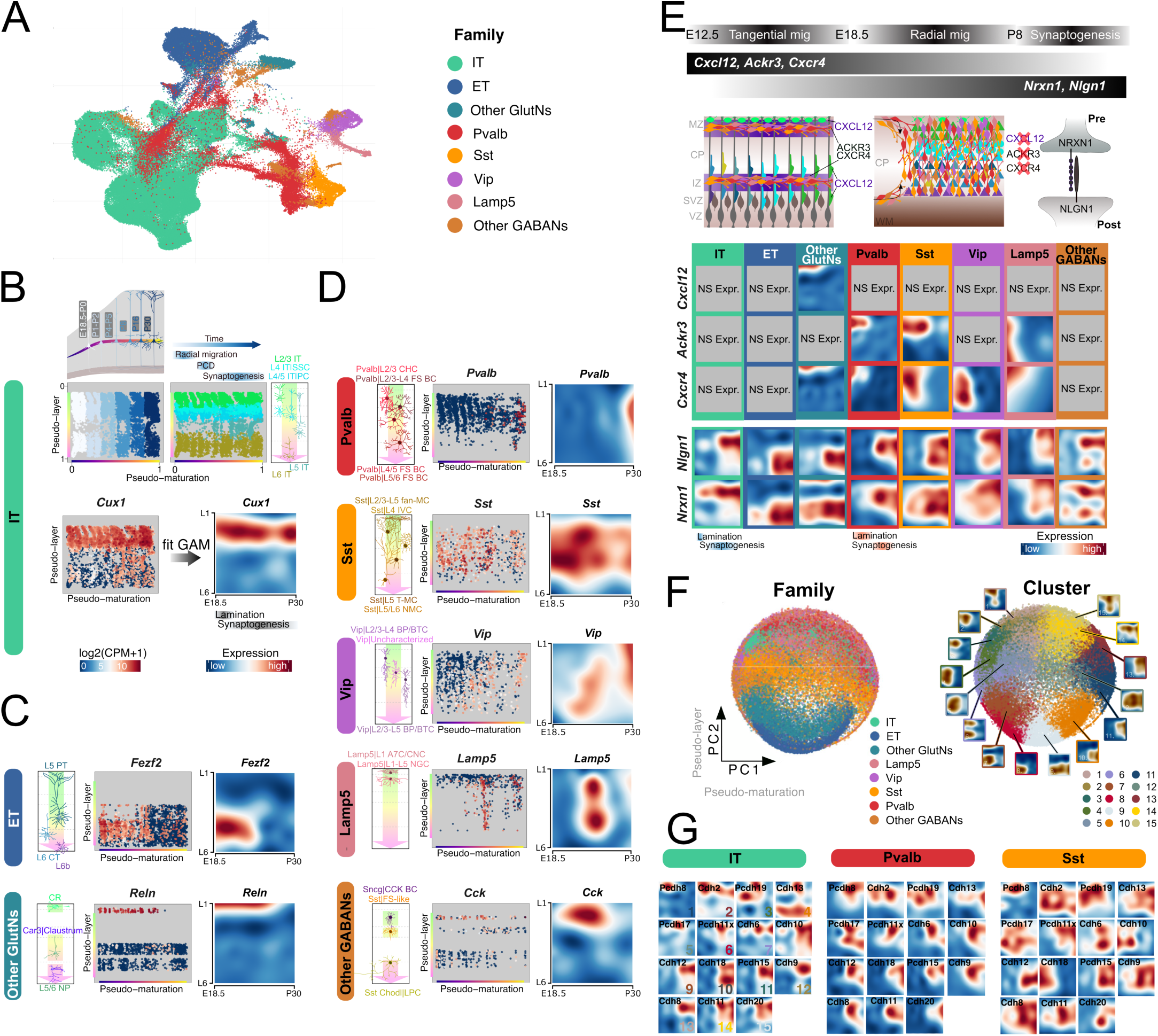
Spatiotemporal transcriptional dynamics of all neuronal cell-types during somatosensory cortex development. (A) UMAP visualization of all neurons color coded with cell family labels. (B) Top: 2D map in which cells are embedded according to their pseudo-maturation (x-axis) and pseudo-layer (y-axis) scores for the IT *family*. Bottom: A generalized additive model (GAM) was applied to generate 2D maps for the spatiotemporal expression of genes (”transcriptional landscape”) throughout somatosensory cortex development; Cux1 expression is represented as an example. (C) Transcriptional landscapes for *Fezf2* in ET (top) and for Reln in other GlutNs (bottom). (D) Transcriptional landscapes for representative genes in each GABAN *family*. (E) Example of transcriptional landscape profiles for genes implicated in GABAN migration (*Cxcl12*, *Ackr3* and *Cxcr4*), and in synaptogenesis (*Nlgn1* and *Nrxn1*). (F) Gene map obtained by performing a PCA on all the significantly expressed genes colored by cell family (left) and by spatiotemporal cluster (right). Average cluster’s transcriptional landscapes are displayed around the gene map. (G) Varying spatiotemporal clusters for the Cadherin gene family among 3 neuronal subtypes. CP, cortical plate; IZ, intermediate zone; MZ, marginal zone; SVZ, subventricular zone; VZ, ventricular zone; WM, white matter. See also Figs S5-S16.

For validation, we examined whether the transcriptional landscapes of specific LRs aligned with their known functions in migration and/or synaptogenesis. We focused first on CXCL12 and its receptors CXCR4 and ACKR3, critical for the tangential migration of GABANs ^35,36^. The literature clearly indicates that CXCL12 is expressed during the perinatal period by the meninges ^37^ and by immature subventricular zone/intermediate zone GlutNs ^38^. Additionally, CXCL12 might also be expressed in CR cells ^39^, although this requires further confirmation. Meanwhile, CXCR4 and ACKR3 are consistently reported to be co-expressed on migrating GABANs ^40,41^, with CXCR4 also expressed in CR cells ^40^. The postnatal extinction of CXCL12 expression has been proposed to drive the transition of GABANs from tangential to radial migration and cortical plate invasion (Fig. 2E) ^35,36,42^. Our transcriptional landscapes corroborated these documented spatio-temporal expression patterns fully (Fig. 2E). Specifically, the enrichment of Cxcl12 in CR cells was confirmed, and the expression of Cxcl12, Cxcr4, and Ackr3 was restricted to early developmental stages (E18.5-P0). This timing aligns with their established role in migration, while the absence of expression at later stages (P0 onward) supports their lack of involvement in synaptogenesis (Fig. 2E).

We then investigated a LR pair associated with synaptogenesis rather than migration. It is well established that the formation of synapses requires specific adhesion molecules, including the NRXN1-NLGN1 LR pair (Fig. 2E) ^43,44^. Our landscapes showed Nrxn1 and Nlgn1 expression in all cell types during time windows consistent with synaptogenesis (Fig. 2E), supporting the role of these proteins in this process.

Analysis of transcriptional landscapes using PCA and k-means clustering allowed us to identify 15 archetypal gene expression profiles. These profiles span early to late developmental stages and map to neurons in deep versus superficial cortical layers (Fig. 2F). This analysis uncovered diverse spatiotemporal patterns: some genes were associated with early developmental processes like lamination, while others were linked to later processes such as synaptogenesis. Additionally, certain genes were specific to neuron types in superficial layers, others to deep layers, while some exhibited a combination of temporal and spatial specificity. Intriguingly, members of the Cadherin gene family were distributed across all 15 clusters (Fig. 2G), highlighting their extensive spatiotemporal diversity and roles throughout cortical development.

We have developed a <u>Shiny App</u> called *scLRSomatoDev* (weblink to be released) to facilitate the exploration of developmental gene expression functions. This app allows researchers to visualize the spatiotemporal expression patterns of any gene(s) across neuronal cell types during cortical development. Users can choose from various visual representations to explore the data, providing an intuitive and comprehensive view of gene expression dynamics (Fig. S11).

### An atlas for inference of LR mediated neuronal interactions during cortical development

The underlying hypothesis driving our study posits that interactions mediated by ligand-receptor (LR) pairs between neuronal subtypes, and in particular between GlutNs and GABANs, play a pivotal role in orchestrating the stereotypical construction of the cortex. We thus anticipated that these interactions would result in highly correlated spatiotemporal expression patterns between GlutN ligands and their corresponding GABAN receptors, and vice versa, compared to random gene pairs. To investigate this, we first assembled LR_DB_2025, the largest curated ligand-receptor database to date. This database was created by aggregating existing LR datasets and manual add-ons from published studies (Fig. 3A & S12, Table S1, see Methods). LR_DB_2025 contains 8,789 curated LR pairs, encompassing 1,421 ligands and 1,233 receptors. Each ligand and receptor were further categorized based on their known molecular functions or their association with specific neurodevelopmental stages (see Methods).

**Figure 3:**
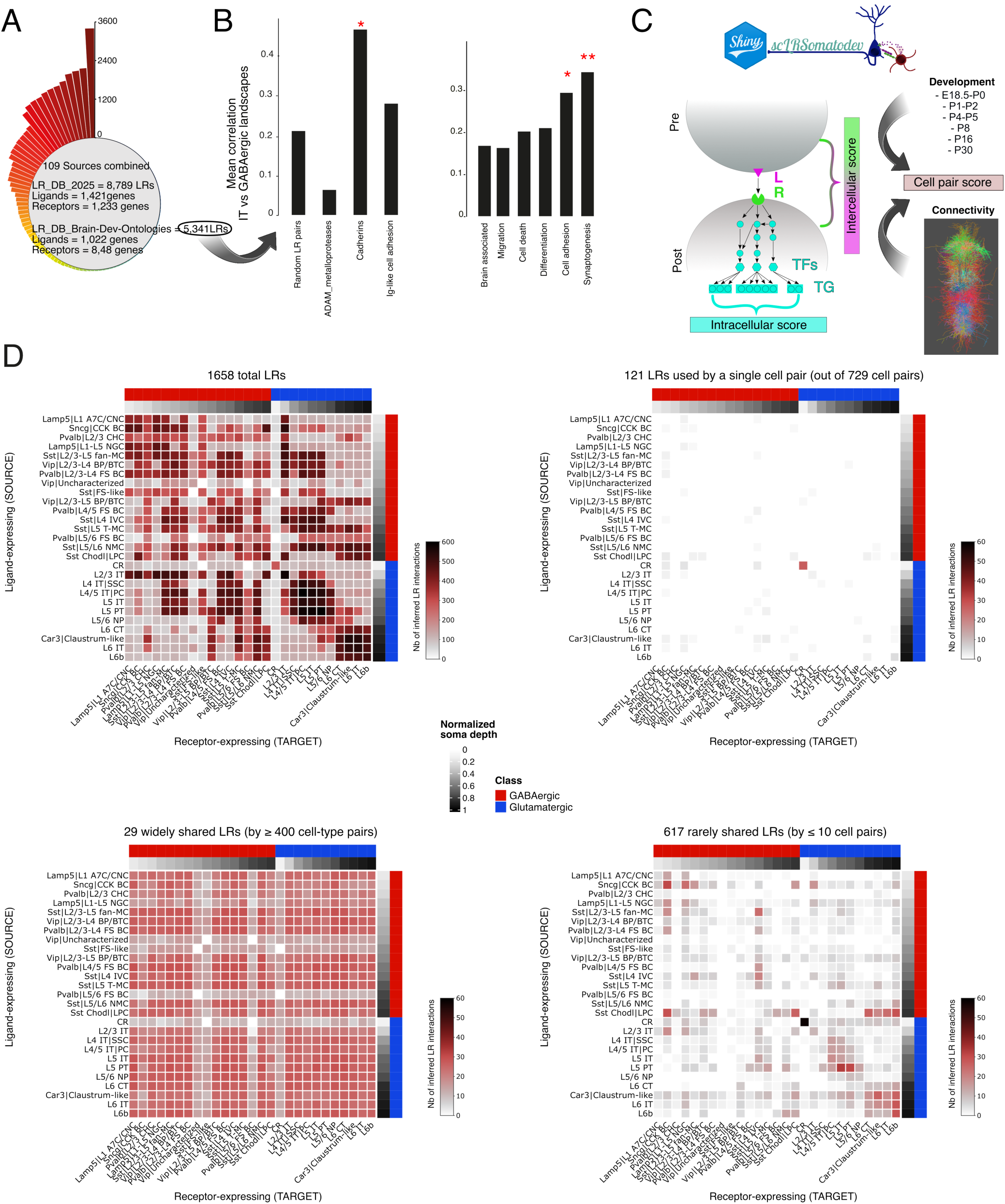
*scLRSomatoDev*, a LR interactome prediction atlas between GlutNs and GABANs over cortex development. (A) Curated database of 8,789 LR pairs. DB, database; L, ligand; R, receptor. LR_DB_2025 contains all LRs, LR_DB_2025_Brain contains brain-expressed LRs with significant expression landscapes. See Tables S1 and S2. (B) High correlation of neurodevelopmental process-associated LR transcriptional landscapes suggest a LR molecular code for neuronal adhesion and synaptogenesis, as well as for the Cadherin family of adhesion molecules. (C) Simplified computational method for the inference of LR-mediated interactions. TF, transcription factor; TG, target gene; S_inter, score for intercellular communication; S_intra, score for intracellular signaling. See Fig. S17-18. (D) Heatmaps illustrating the number of inferred LR interactions that persist across at least two consecutive ages for all 729 *cell-type* pairs. From left to right and top to bottom: all possible interactions; interactions unique to a single *cell-type* pair, widely shared interactions (by >400 pairs) and very rare interactions (shared by <10 pairs). See also Figs S12-23.

For LRs associated with developmental processes, such as cadherins, the transcriptional landscapes of ligands in IT neurons and their receptors in GABANs showed high correlations (Fig. 3B, Fig. S13-S16). This highlights the critical role of cadherin-mediated interactions in GlutN-GABAN communication during development, supporting hypotheses linking cadherins to areal specialization ^34,45^ and synaptic specificity ^46^.

LRs tied to migration, differentiation, cell death, cell adhesion, and synaptogenesis (Table S2, Methods) exhibited varying degrees of spatiotemporal correlation (Fig. 3B). Notably, synaptogenesis displayed the strongest LR correlations, suggesting it heavily depends on cell-cell communication, whereas processes like cell death or differentiation may rely more on intrinsic mechanisms (Fig. 3B).

To identify which LRs drive cell-cell interactions influencing cortical development, we constructed a comprehensive LR atlas that infers likely strong LR interactions between neuronal cell types from E18.5 to adulthood. This approach integrates temporal cell-gene expression matrices with data from our LR database (Fig. 3C and S17, see Methods).

First, we implemented and adapted to our dataset the *scSeqComm* method, which computes intercellular LR scores for each cell pair, based on the relative expression levels of ligands in source cells and receptors in target cells ^47^.

To improve accuracy, we created a cell pair score that prioritized LRs based on known developmental timing and connectivity patterns. The cell pair score is the modified sum of a development score (Fig. 3C & S17, Table S3) accounting for when cell types populate the cortex and a connectivity score that estimates synaptic connections based on physical proximity. The connectivity score was calculated from the morphological reconstruction of 1000 neurons in a cortical column from patch-seq studies (Fig. 3C, S17-18; Table S4). LRs with secreted ligands, we gave a lower weight to the cell pair score and a higher weight to the LR intercellular score to account for their ability to act over greater distances. To minimize false positives further, we only included LR pairs that were significantly present across contiguous developmental stages (Fig. S17).

To enhance robustness and minimize false positives in our LR predictions, we incorporated weights into each selected *cell-type* pair that reflect known patterns of developmental synchronicity and connectivity. Specifically, a *development* score (Fig. 3C & S17, Table S3) accounted for the varying times at which different cell types populate the cortex, affecting the likelihood of certain interactions depending on the mouse’s age (Fig. S18). A *connectivity* score estimated the degree of synaptic connectivity between cell types, based on their physical proximity (Fig. 3C, S17-18; Table S4). These scores were integrated to generate a *cell-type* score, which was applied to each cell pair to incorporate prior knowledge into the threshold calculation for predicting significant LR-mediated intercellular communications (Fig. 3C and S17, see Methods). For LR pairs involving secreted ligands, we adjusted the *cell-type* score by assigning a lower weight, acknowledging that these interactions can occur over greater physical distances compared to non-secreted ligands (Fig. S17). To further reduce false positives and ensure robust identification of LR pairs, we retained only those pairs that were significantly present between two contiguous developmental stages (Fig. S17).

For LRs with available information, our modified *scSeqComm* pipeline quantified intracellular signaling pathways downstream of receptor activation in target cell types, by integrating data from public regulatory gene databases ^47^ (see Methods). This approach generated an intracellular signaling score (Fig. 3C and S17) that allowed us to evaluate the reliability of inferred LR communications. It also facilitated the identification of the preferred receptor(s) engaged when a ligand interacted with multiple receptors, enhancing the understanding of specific LR dynamics within cellular contexts.

Inferences of LR interactions over cortex development are accessible via the <u>Shiny App</u> *scLRSomatoDev* (to-be-released, Fig. S19), allowing researchers to visualize both number and identities of predicted LR interactions between any of the 729 possible neuronal *cell-type* pairs at each of 7 developmental stages covering E18.5-adulthood. Numbers of LR interactions are depicted by heatmaps (Fig. 3D, S19A), while the identities of LR pairs and associated gene ontologies (GO) pathways are represented by dot plots (Fig. S19B).

Which molecular codes allow specific LR pairs to control the position and connectivity of cell types? To explore this, we conducted a comprehensive analysis of our LR predictions. Across 729 cell-type combinations from E18.5 to adulthood, we quantified the number of predicted LR interactions maintained between at least two consecutive stages in the E18-Adult temporal window. We categorized them into unique, “widely shared” (shared by over 400 cell pairs), and “rarely shared” (used by ≤ 10 cell pairs) (Fig. 3D). The total number of predicted LR interactions for specific cell combinations reached up to ∼600, whereas interactions exclusive to unique cell pairs were exceedingly rare. Widely shared LRs, i.e., LRs shared by ≥ 400 cell pairs, were relatively infrequent, accounting for less than 10% of total inferred LRs (Fig. 3D). In contrast, rarely shared LR pairs (by ≤ 10 cell pairs) showed variable frequency, with up to 60 detected in CR-CR interactions. These results suggest that cortical neuron subtypes rely on a diverse repertoire of LR pairs, combining both rare and widely shared interactions in specific patterns to drive their distinct communication and connectivity. Examining the number of inferred LR interactions at different developmental ages further reinforced the conclusion from the global analysis: LRs used exclusively by a single cell pair remained rare regardless of developmental age. This held true even when fixing one of the two cell types in the pair while varying the other (Fig. S20–S22).

By analyzing the number of significant interactions between GlutN and GABAN cell types, we observed that when GlutNs served as the ligand sender, DL GlutNs, as well as UL GlutNs, exhibited stronger interactions with DL GABANs during early developmental stages (Fig. S23). The predominant interactions between UL GlutNs and DL GABANs (compared to UL GABANs) decreased with age (Fig. S23). Consistent with previous findings, UL GlutNs at early developmental stages relied predominantly on intrinsic genetic programs rather than extrinsic inputs (Fig. S7), which aligns with the low number of interactions observed between UL GlutNs and UL GABANs at young ages, followed by an increase in these interactions with maturation. In the reverse direction, when GABANs acted as the ligand source and GlutNs as the target, both UL and DL GABANs interacted significantly more with GlutNs within the same layer from E18.5-P0 to P8 (Fig. S23). By P16 and P30, however, a notable exception emerged, with DL GABANs displaying increased interactions with UL GlutNs (Fig. S23).

### Key roles for LR interactions during synaptogenesis and in neurodevelopmental diseases

We analyzed temporal LR predictions between GlutNs and GABANs. The total number of LRs predicted increased over time (Fig. 4A), which aligns with the gradual establishment of synaptic connectivity. Among GABAN subtypes, Sst and Pvalb neurons were the first to receive a high number of LR interactions from GlutNs (Fig. 4A), while Lamp5, Vip, and other GABAN populations exhibited a later increase. Notably, Pvalb cells initially had fewer LRs than Sst neurons but gradually caught up, consistent with studies showing that Sst neurons precede Pvalb neurons in the formation of cortical networks ^48^. Furthermore, the fact that Sst and Pvalb cells received the most LRs (Fig. 4A) suggests that these subtypes may rely more heavily on non-cell-autonomous signals for their integration into cortical networks than other GABAN subtypes.

**Figure 4:**
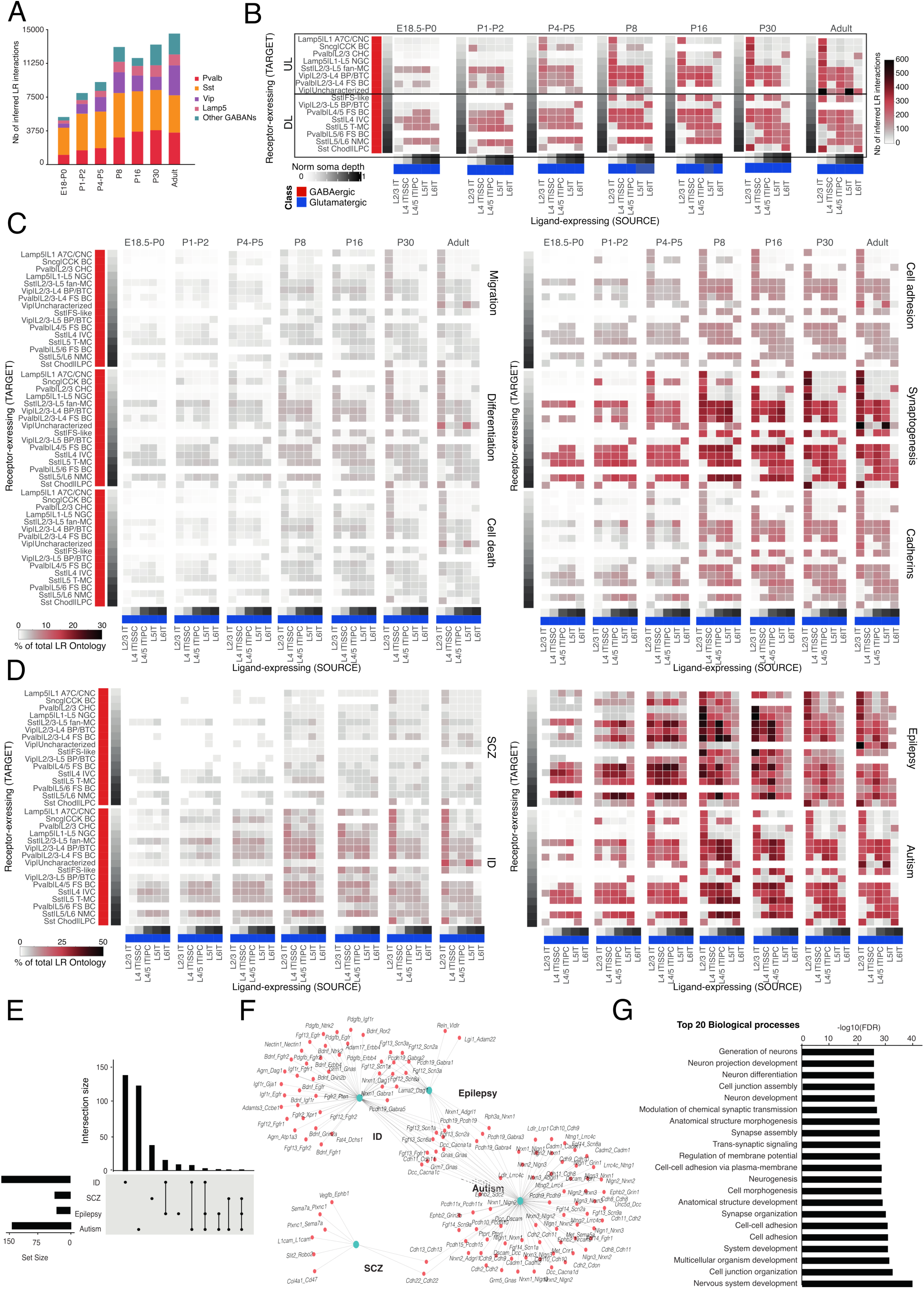
Temporal analysis of LR pairs associated with neurodevelopmental processes and diseases. (A) Number of LR pairs predicted to be utilized per GABAN family as target cells, from E18 to adulthood. (B) Number of predicted LR interactions between IT and GABAN *cell-type*s as source and target cells, respectively, from E18 to adulthood. (C) Percentage of LR pairs of 5 main neurodevelopmental ontologies and of the cadherin LR family that are predicted as utilized between IT and GABAN *cell-type*s as source and target cells, respectively, from E18 to adulthood. (D) Percentage of LR pairs with both L and R associated with neurodevelopmental diseases that are predicted as utilized between IT and GABAN *cell-type*s as source and target cells, respectively, from E18 to adulthood. (E) Upset plot showing intersections between LR pairs of the 4 diseases studied. (F) Interaction network plot showing the LR pairs associated with each disease. (G) Gene ontologies associated with inferred, disease-associated LR pairs.

We then examined the temporal progression of LR predictions for each *cell-type*, using IT and GABAN subtypes as examples of source and target cells, respectively. Overall, the number of predicted LRs increased over time (Fig. 4B), aligning with the trends observed in the global analysis (Fig. 4A). However, different patterns emerged for UL and DL interactions, i.e., interactions involving UL vs DL GABAN targets (Fig. 4B). In UL interactions, LRs were minimal at early stages but increased progressively, reaching high levels in adulthood. Conversely, DL interactions exhibited relatively high LR numbers early on, with only modest increases as the organism aged. This suggests that the earlier developmental onset of DL neurons enables them to establish LR connections at a younger age compared to UL neurons.

We then calculated the proportion of LRs associated with five key neurodevelopmental processes—migration, differentiation, cell death, cell adhesion, and synaptogenesis—that are actively utilized during corticogenesis.

LRs associated with migration, differentiation, and cell death showed minimal inferred utilization, suggesting that these neurodevelopmental processes rely less on LR-mediated cell-cell communication and more on cell-intrinsic mechanisms. In contrast, synaptogenesis-associated LRs, and to a lesser extent cell adhesion-associated LRs, were predicted to be strongly utilized from E18.5 to adulthood (Fig. 4C). Notably, Cadherins exhibited strong utilization from P4 onwards, highlighting their potential role in regulating cell-cell interactions during adhesion and synaptogenesis in the cortex.

We next investigated the timing and involvement of LRs associated with neurodevelopmental diseases during development. Using documented gene-disease associations, we categorized the LRs from the LR_DB_2025 into four major neurodevelopmental diseases: autism, epilepsy, schizophrenia (SCZ), and intellectual disability (ID) (Table S5). Notably, the percentage of utilized LRs linked to SCZ and ID remained consistently low across developmental stages (Fig. 4D). In contrast, LRs associated with autism and epilepsy were utilized at high rates, occasionally reaching up to 50%, and their prevalence increased progressively over time (Fig. 4D). This suggests that the etiology of autism and epilepsy may be more closely related to LR-mediated cell-cell communications in cortical areas compared to SCZ and ID. Additionally, some LRs were associated with multiple diseases (Fig. 4E), and notably, the only two significantly utilized LRs common to both autism and SCZ were the Cadherins, *Cdh13* and *Cdh22* (Fig. 4F). Consistent with expectations, gene ontology analysis revealed that predicted disease-associated LRs were enriched in neurodevelopmental biological processes (Fig. 4G).

### Validation of the LR prediction atlas: Nrg3-Erbb4

We queried our LR atlas to validate known interactions between GlutNs and GABANs, focusing on the Nrg3-Erbb4 pair, which is recognized for mediating the development of excitatory synapses onto GABANs and is linked to neurodevelopmental disorders ^49^ (Fig. 5A). First, consistent with the literature, transcriptional landscapes confirmed Nrg3 expression in GlutNs and the restriction of Erbb4 expression to GABANs (Fig. 5B). Nrg3 and Erbb4 expression levels peaked in the second half of the developmental timeline, aligning with synaptogenesis (Fig. 5B). Second, our atlas predicts significant Nrg3-Erbb4 interactions from E18.5-P0 to P30 between GlutNs (Nrg3 source) and most GABANs (Erbb4-expressing target cells) (Fig. 5C). Regardless of mouse age, the intracellular signaling score for Nrg3-Erbb4 was strongest in pairs for which target cells were Pvalb BCs and Vip|L2/3−L5 BP/BTCs, two GABAN types best known to require Erbb4 signaling for the formation of excitatory synapses (Fig. 5D) ^12,13,50^. Overall, our atlas supports the role of the Nrg3-Erbb4 interaction in excitatory synapse formation onto specific GABAN types during corticogenesis.

**Figure 5.**
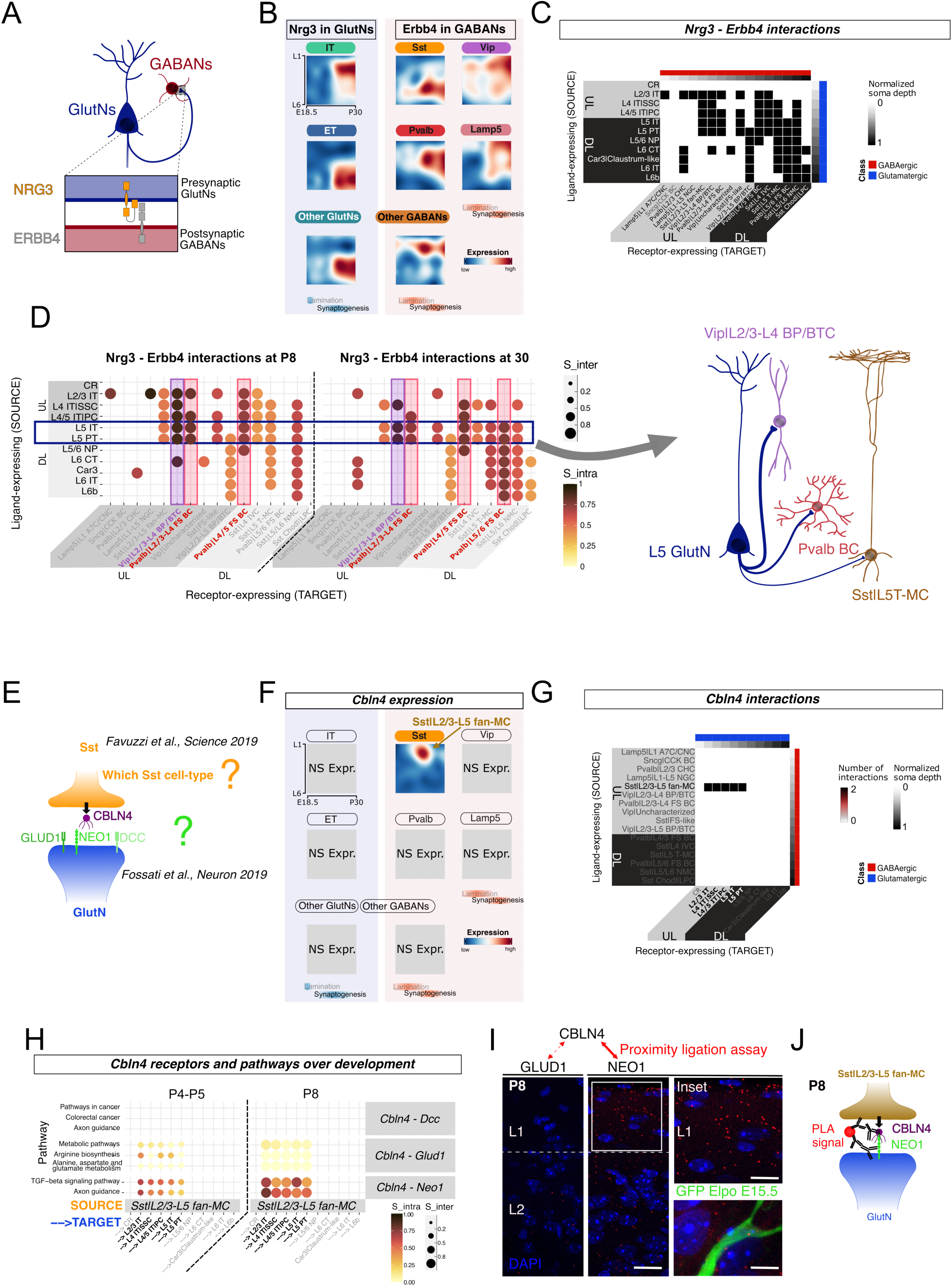
Validation and Application of the Ligand-Receptor Atlas for Knowledge Completion. (A) Diagram illustrating NRG3-ERBB4 interaction between glutamatergic and GABAergic cells. (B) *Nrg3* and *Erbb4* expressions in transcriptional landscapes of glutamateric and GABAergic neuron *families*, respectively. (C) Heatmap depicting predictions of *Nrg3*-*Erbb4* mediated cell-cell interactions between GlutN and GABAN *cell-type*s. Blue represents GlutN types, red represents GABAN types. The white to black gradient indicates the median normalized soma depth, with 0 at the pial surface and 1 at the white matter. (D) Left: Dot plot of predicted *Nrg3*-*Erbb4* mediated cell-cell interactions at P8 and P30. GlutNs (blue) are sources of *Nrg3* and GABANs are targets expressing *Erbb4*. Right: Schematic representation illustrating the predicted interaction strength among three example cell pairs, highlighting preferred connectivity patterns based on computational predictions. (E) Diagram illustrating Cbln4-mediated inhibitory synapse formation from Sst cells to GlutN in the cortex. (F) Transcriptional landscapes showing *Cbln4* expression across different neuronal families. (G) Heatmap of LR pairs involving *Cbln4*. (H) Dot plot of predicted Clbn4 mediated cell-cell interactions at P4-P5 and P8. Pathways are shown on the left. CBLN4-receptor pairs are represented on the right. (I) Proximity ligation assay (PLA) results for interactions between CBLN4 and GLUD1 and CBLN4 and NEO1 in cortical layers L1-2. A magnification of a green process from a GFP-electroporated neuron (E15.5) with red puncta on it is shown on the lower right. (J) Summary of the experimental findings and predictions resulting from our analysis. S_inter, intercellular score; S_intra, intracellular score.

### NEO1 is the main CBLN4 receptor at Martinotti-glutamatergic developing synapses

We then investigated whether our gene expression and LR atlases could shed new light on known but incompletely understood cell-cell communications. We focused on CBLN4, a ligand that facilitates synapse formation between the axonal termini of Sst neurons and the apical dendrites of GlutNs in the mouse cortex ^11,51^ (Fig. 5E).

Despite this knowledge, which precise Sst *cell-type(s)* express(es) Cbln4 and what its receptor(s) is/are in GlutN apical dendrites are two pending questions ^51,52^ (Fig. 5F) that we addressed using our gene expression and LR atlases. Our analysis of transcriptional profiles revealed that among the 4 identified Sst *cell-types* (Sst|L2/3-L5 fan-MC, Sst|L4 IVC, Sst|L5 T−MC, and Sst|L5/L6 NMCs), only the Sst|L2/3-L5 fan-MCs exhibited strong expression of *Cbln4* (Fig. 5G). Then, our LR interactome atlas predicted that CBLN4 was involved in only very few interactions during peak synaptogenesis (P5 to P30), *i.e.*, between Sst|L2/3-L5 fan-MCs as “source” cells and some GlutN types (L2/3 IT, L4 IT|SSC, L4/5 IT|PC, L5 IT and L5 PT) as “target” cells (Fig. 5G) - more unexpectedly, most GABAN *cell-type*s were also predicted as “target” cells. Finally, intercellular signaling scores revealed that among the 3 known CBLN4 receptors (DCC, GLUD1 and NEO1), only GLUD1 and NEO1 were predicted to be involved (Fig. 5H). These intercellular scores exhibited an increase from P4 to P8, whichaligns with earlier findings underscoring the significance of CBLN4 during this developmental period ^11^ and supports its role in synaptic formation.

We then took advantage of the intracellular score to determine whether GLUD1 or NEO1 might predominate as a CBLN4 receptor in the cortex. Interestingly, while earlier studies showed that GLUD1 is a CBLN4 receptor in cortical GlutNs around P21-P30 ^11,51^, our intracellular score predicts that NEO1 downstream signaling is more active than GLUD1 downstream signaling in CBLN4 “receiving*”* neurons at peak synaptogenesis around P4-P8 (Fig. 5H). NEO1-associated genes in “target” GlutNs were notably linked with axon guidance and TGF-beta signaling pathways (Fig. 5H). Based on these predictions, we investigated the presence of a direct interaction between CBLN4 and NEO1 or GLUD1 at P8 in situ. Using proximity ligation assay (PLA), we detected strong interactions between CBLN4 and NEO1 in cortical layers L1-2, while interactions between CBLN4 and GLUD1 were minimal or undetectable, as indicated by the sparse or absent PLA signals (Fig. 5I). Overall, these findings underscore the utility of our gene expression and LR atlases in pinpointing the precise cellular sources of ligands and their primary receptors in target cells.

### Uncovering novel interactions: CDH13 and PCDH8 as mediators of perisomatic inhibition in deep and superficial layers, respectively

We asked whether our LR atlas can identify novel LR pairs regulating the establishment of specific connectivity patterns. We focused on the cadherin *family* of adhesion molecules that are thought to play such a role in the cortex ^45,46,53^. Analysis of transcriptional landscapes revealed notable diversity in the spatio-temporal expression profiles of cadherin family members across different cell types (Fig. 2G).

The atypical *Cdh13*, associated with neurodevelopmental disorders ^54,55^, was particularly interesting as it showed highest expression in DL GlutNs and MGE-derived GABANs (Pvalb & Sst) (Fig. 6A). *Cdh13* expression was notably high in DL GlutNs neurons through cortical development, and in Pvalb and Sst neurons around the P4-P30 synaptogenesis period ^11^ (Fig. 5A). This suggests a role for *Cdh13*-*Cdh13* mediated intercellular communication in the establishment of connectivity between MGE-derived GABANs and DL GlutNs. Accordingly, our LR atlas indeed inferred that *Cdh13*-*Cdh13* might mediate interactions that are mainly between Sst / Pvalb GABANs and DL GlutNs. It was a strict rule at specific time points like P4-P5 and P30 (Fig. 6C), and there were only exceptions when considering all time points collectively (33/41 involved Pvalb/Sst->DL GlutN pairs, Fig. 6B). These findings identify Cdh13 as a potential candidate molecule for establishing connectivity between Pvalb/Sst GABANs and DL GlutNs, consistent with recent evidence suggesting that Cdh13 mediates the perisomatic inhibition of Pvalb BCs onto layer 5 extratelencephalic (L5 ET) neurons ^56^. We used the fact that synaptic connectivity between Pvalb and GlutNs is measurable with SYT2 staining ^56,57^ to investigate the implication of CDH13 in the development of perisomatic inhibition mediated by Pvalb BCs on DL GlutN somata. After *Cdh13* knock-down in L5 GlutNs by in utero electroporation, we assessed SYT2 perisomatic inhibition at P28, after migration and establishment of connectivity have occurred. As expected, the surface area of DL GlutN soma covered by SYT2^+^ boutons was significantly reduced by *Cdh13* knock-down (Fig. 6C). This result shows that post-synaptic CDH13 is necessary for proper Pvalb BC-perisomatic inhibition of DL GlutN somata. The specificity of this regulation was further supported by the observation that Pvalb chandelier cell (CHC) innervation of the axon initial segment was unaffected, as predicted by our LR atlas (Fig. 6D).

**Figure 6.**
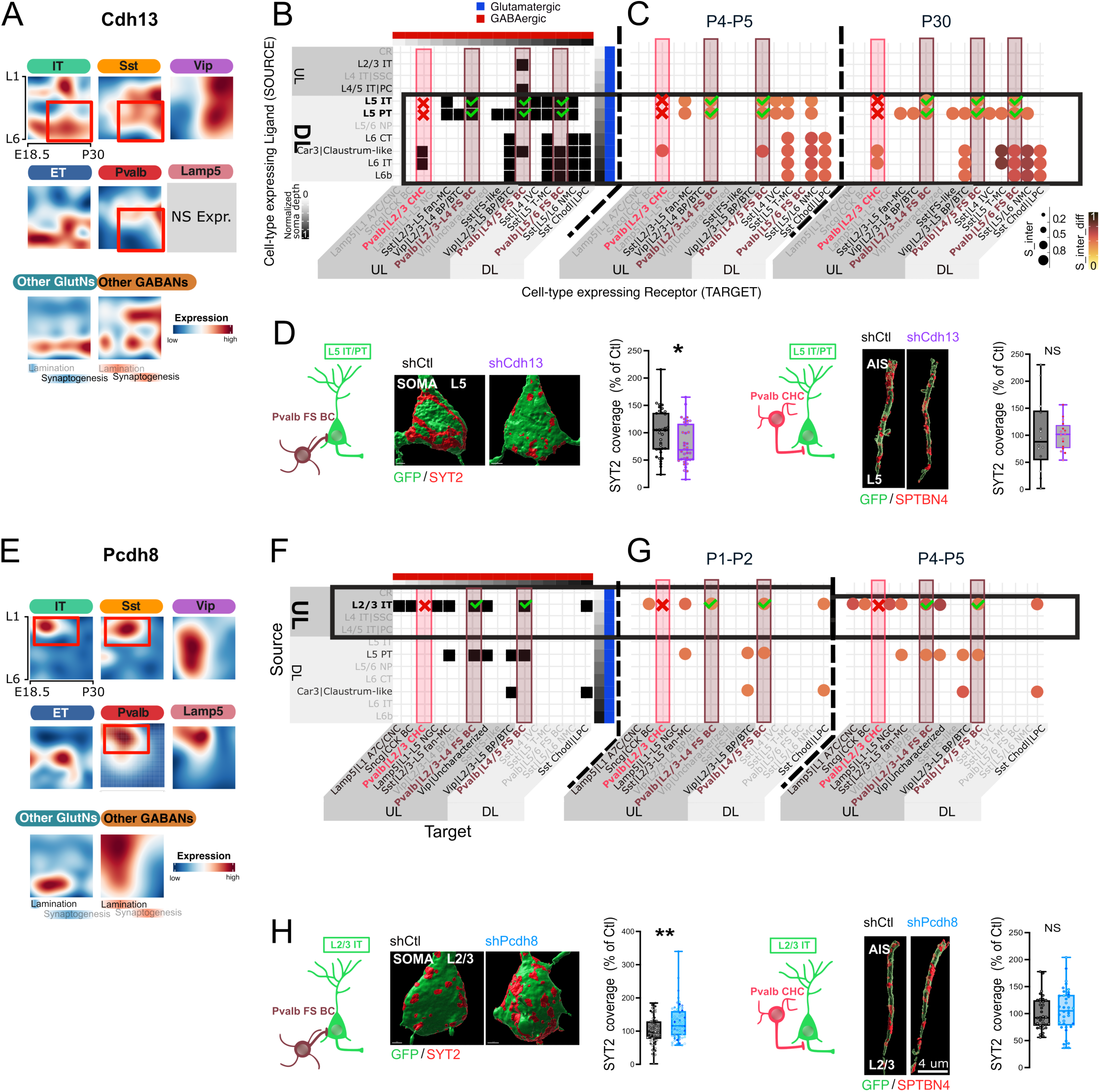
Cadherins Cdh13 and Pcdh8 regulate perisomatic inhibition in deep and superficial cortical layers. (A) Transcriptional landscapes of Cdh13 expression across different cell families. (B) Heatmap of predicted Cdh13-Cdh13 interactions between GlutN and GABAN cell types shared by at least two consecutive ages. (C) Utilization of Cdh13-Cdh13 interactions across GlutNs and GABANs at P4-P5 and P30, with interaction strength quantified by the inter_diff score. (D) Perisomatic (left) and axon initial segment (AIS, right) inhibitory innervation of E13.5-electroporated L5 GlutNs by SYT2-expressing Pvalb BC and CHC boutons, respectively, in control or Cdh13-knockdown (shCdh13) GlutNs (Mann-Whitney test, *p-value < 0.05, n=3-4 mice/condition; ≥ 36 and 14 cells/condition for soma and AIS, respectively). (E) Transcriptional landscapes of Pcdh8 expression across different cell families. (F) Heatmap of predicted Pcdh8-Pcdh8 interactions between GlutN and GABAN cell types shared by at least two consecutive ages. (G) Predicted Pcdh8-Pcdh8 interactions at P1-P2 and P4-P5, with stronger expression in L2/3 IT-involving cell pairs. (H) Perisomatic (left) and axon initial segment (AIS, right) inhibitory innervation of E15.5-electroporated L2/3 GlutNs by SYT2-expressing Pvalb BC and CHC boutons, respectively, in control or Pcdh8-knockdown (shPcdh8) GlutNs (Mann-Whitney test, **p-value < 0.01, n=3-4 mice/condition; > 61 and 35 cells/condition).

We then turned our focus to PCDH8 because its laminar expression is relatively complementary to the one of DL-enriched CDH13, i.e., *Pcdh8* is enriched in both GlutNs and Pvalb/Sst subtypes of UL (Fig. 6E). Out of the only 8.5% of all possible GlutN-GABAN interactions predicted to involve *Pcdh8* (15/176), L2/3 ITs displayed the highest fraction, i.e., 50% (8/16 possible interactions). The fraction of their possible interaction with Pvalb BC types reached 66.7% (2/3 possible interactions) (Fig. 6F). Also, inferred *Pcdh8*-mediated interactions were more prominent in cell pairs involving L2/3 IT neurons compared to those involving L5 PT or Car3|Claustrum-like neurons (Fig. 6G).

Therefore, we used in utero electroporation of a *Pcdh8* shRNA at E15.5 to investigate the role of PCDH8-PCDH8 interactions in Pvalb-mediated perisomatic inhibition of L2/3 IT neurons. PCDH8 knock-down significantly increased SYT2 coverage of GFP-electroporated cells compared to a control situation, which indicates that PCDH18 regulates perisomatic inhibition of L2/3 IT neurons negatively (Fig. 6H). Thus, our LR atlas allowed us to conclude that different cadherins modulate perisomatic inhibition in different ways depending on the layer examined.

## Discussion

Seminal studies in the last decades have indicated that the stereotyped laminar positioning and connectivity of cortical GABAN subtypes are both in large part extrinsically regulated ^15–17^. These data suggest that a molecular code based on LR interactions between firstly settled GlutNs and secondarily arriving GABANs might orchestrate cortical circuit assembly. In this work, we utilized high-throughput scRNA-seq to address this question, by attempting to identify LR pairs that are essential for GlutN-GABAN interaction and cortical circuit assembly. All results from the LR interaction inference, along with gene expression visualizations, are available in a <u>Shiny App</u> *scLRSomatoDev* (to-be-released).

We first characterized the transcriptome of all the main known neuron types over the course of cortical development. We provide this dataset, which results from an extensive integration and annotation effort, as a new reference to capture the dynamic transcriptional landscape of corticogenesis (<u>Shiny App</u>). We then focused, for LR interaction assessment, on the E18-P30 critical period during which GABANs reach their preferred laminar position and establish synaptic connections with GlutNs and other GABAN partners. We developed computational tools to: (1) assess spatiotemporal expression of all genes in all cortical neuron types and (2) infer the number and identities of significant LR interactions between all neuron types that might control stereotypical cortical assembly. We provide evidence that these tools can be utilized to: (1) validate true ground observations (e.g., Nrg3 - Erbb4), (2) complete knowledge about known LR interactions (e.g., Cbln4 – Neo1 for Sst MC-> GlutN interaction) and (3) identify novel LR pairs with a role in the formation of specific connections (*e.g.*, the *Cdh13* and *Pcdh8* cases).

Ligand-receptor mediated cell-cell interactions constitute a fundamental process that participates in shaping most biological tissues. Before the development of scRNA-seq, studying cell-cell interactions was low-throughput, limited to a short selection of genes/proteins and one to few *cell-type* pairs. Since the advent of scRNA-seq, a growing number of computational tools has been developed to infer cell-cell communication. Most tools are based solely on the gene expression level of the LR pair between 2 *cell-types* ^58–60^, but few tools provide a more accurate reflection of ground truth by considering the response triggered by LR binding in the cell expressing the receptor ^47,61–63^. Here, we benchmarked most of these methods and ended up choosing one of the most recent ones that has the advantage of inferring both intercellular and intracellular signaling for a given LR pair between *cell-type* pairs: scSeqComm ^47^. The fact that scSeqComm returns these scores between 0 and 1 makes it easier to increment a threshold to identify relevant LR pairs. For the intercellular score, we determined a threshold using prior knowledge taking into account the final connectivity between *cell-types* (connectivity score) as well as the age (development score) at which the analysis was conducted. This prior knowledge allowed us to increase stringency and robustness of our inferences. Furthermore, by computing an intracellular score, scSeqComm quantifies the evidence of ongoing intracellular signaling triggered by the LR binding in the target cell. This intracellular score allowed us to increase the reliability of inferred communication. Custom thresholds can be set for both intercellular and intracellular scores according to the desired reliability of the inferred interactions. To further increase the reliability of the inferred interactions, we created a meticulously curated LR database by amalgamating existing published databases. Additionally, we incorporated LR pairs from the literature to enrich its content.

Our analysis of LR predictions reveals that neuronal connections are often determined by specific combinations of LRs with distinct specificities. Notably, a core group of approximately 40 broadly expressed LRs mediate interactions in over 50% of cell-type pairs, complemented by a larger subset of LRs with varying degrees of specificity. Importantly, LRs unique to individual neuronal connections are exceedingly rare, suggesting that neuronal connectivity is predominantly shaped by shared and context-dependent LRs rather than by unique or isolated interactions.

Recent studies have shown that members of the cadherin family of adhesion molecules are differentially expressed in subtypes of cortical excitatory and inhibitory neurons ^34,45^. This suggested that a cadherin code may orchestrate the stereotypical organization of cortical circuits. Our findings, which demonstrate that cadherin superfamily members exhibit *cell-type*-pair specific spatiotemporal expression profiles (Fig. S13-S16), and that CDH13 and PCDH8 regulate critical aspects of deep versus superficial circuit formation (Fig. 5), support this hypothesis. Future functional studies based on our LR atlas will be crucial in confirming or refuting this hypothesis.

Based on hypotheses generated by our LR atlas, we experimentally identified two cadherin superfamily members, CDH13 and PCDH8, as regulators of Pvalb BC mediated perisomatic inhibition in DL and SL, respectively. Our observation that *Cdh13*-*Cdh13* interactions are necessary for perisomatic inhibition by Pvalb, but not Sncg, BCs in DL, aligns with a recent study^56^.

In contrast to the promotive role of CDH13 in DLs, we find that PCDH8 acts as a suppressor of perisomatic inhibition in ULs. These findings highlight the predictive power of our LR atlas in identifying both facilitatory and inhibitory interactions, a crucial feature given the role of repulsive interactions in forebrain neurodevelopment. Furthermore, the data suggest that different cadherins or CAM family members may modulate synaptogenesis in region-specific and divergent ways across the cortex.

The suppressive effect of PCDH8 on inhibitory synapse assembly in ULs parallels a similar role observed for another protocadherin, PCDH18, in regulating Sst neuron-to-GlutN synapse assembly ^11^. Collectively, given the established associations of CDH13 and PCDH8 with neurodevelopmental disorders ^54,55,64^, we propose that disrupted Pvalb basket cell-mediated inhibition in specific cortical layers could play a key role in the pathophysiology of these conditions.

In addition to neurons, non-neuronal cells also play a role in the early stages of cortical circuit assembly. For example, blood vessels and ventral oligodendrocyte precursor cells regulate the tangential migration of GABANs from the subpallium to the cortical plate through specific *Cxcl12*-SEMA6A/B-PLXNA3 unipolar contact repulsion ^65^. Future studies should include single-cell transcriptomics-based analyses of neuron-non-neuronal cell communication to further explore these interactions.

Limitations of the approach – Considerations for Future Improvements in Ligand-Receptor Atlases Although our approach is robust and comprehensive in terms of the genes investigated for LR interactions, it does not provide information on the subcellular expression of ligands and receptors across different cell types, which is likely important for explaining specific connectivity patterns^49^. Furthermore, our approach focuses on mRNA expression as a proxy for protein expression, although the two may differ in reality. Recent advances in *in vitro* single cell high-throughput proteomics suggest it will soon be possible to include the required subcellular protein expression information in future atlases ^66^.

Currently, our knowledge of the spatial positions of cortical cell types relative to each other is limited to adult stages ^67,68^, which impedes precise inference of timely LR mediated cell-cell interactions at specific developmental stages, whether transient or stable. The advent of spatial transcriptomic approaches with exceptional sensitivity, throughput, and resolution holds promise to bridge this gap in the near future.

## Acknowledgments

We thank Julien Prados (UNIGE) for providing his Torch model for artificial neural network (ANN)-based cell type identification, and members of the Cardoso laboratory for their input, and the Molecular and Cellular Biology Facility (PBMC), the Animal Core Facility and the Imaging Facility (inMagic) INMED platforms.

## Author contributions

A.d.C. and R.M. initiated the study. A.d.C. and L.T. conceptualized and supervised the study. A.d.C., L.T. and R.M. designed and conceptualized the experiments. A.d.C. and R.M. performed most experiments. R.M. analyzed most experiments, supervised by L.T. and A.d.C.. T.D.N. performed some LR analyses for figures 3 and 4. R.M. generated the Shiny App and L.S. added some improvements. C.C., A.G., L.C. and V.B. performed and analyzed shRNA design/production and in utero electroporations. E.P. validated shRNAs and performed proximity ligation assays. A.d.C. and R.M. wrote the manuscript with inputs from all authors.

## Data and materials availability

All annotated data will be available online once published. All other data are available in the main text or the supplementary materials. License information: Copyright © 2022 the authors, some rights reserved; exclusive licensee American Association for the Advancement of Science. No claim to original US government works. https://www.science.org/about/science-licenses-journal-article-reuse

