## Supplementary Material for "Inferring Ligand-Receptor Interactions between neuronal subtypes during mouse cortical development"

**The PDF file includes:**

Materials and Methods

Figs. S1 to S24

References

**Other Supplementary Materials for this manuscript include the following:**

Tables S1 to S5

#### Materials and Methods

**Animals.** Mice (mus musculus) were group housed (2–5 mice/cage) with same-sex littermates on a 12-hour light-dark cycle with access to food and water *ad libitum*. They were bred and maintained on a mixed SVeV-129/C57BL/6N background. Animal experiments were carried out in accordance with European Communities Council Directives and approved by French ethical committees (Comité d'Ethique pour l'expérimentation animale no. 14; permission number: 62-12112012, Apafis #21683-2019073011285386v4).

**Single-cell isolation.** Mice brains were dissected submerged in a ice-cold bubbled artificial cerebrospinal fluid (ACSF) with carbogen (95% $O_2$ and 5% $CO_2$). Our ACSF consisted of NaCl (7.32 g/L), KCl (0.26 g/L), $NaH_2PO_4$,$H_2O$ (0.165 g/L), $Cacl_2$,$2H_2O$ (0.438 g/L), $MgCl_2$,$6H_2O$ (0.264 g/L), D(+)-Glucose (1.98 g/L), $NaHCO_3$ (2.1 g/L), and acid kynurenic (0.567 g/L). Brains were then sliced into a 300 µm (P0 and P2 mice) or 500 µm (P5, P8 and P30 mice) coronal sections with a vibratome (Leica). Somatosensory cortex area was dissected under a binocular loop. Enzymatic digestion was then processed by using pronase (*Septomyces Argeus* at 1 mg/mL) during 25 min at room temperature (RT) for P0 to P8 datasets. Cells were dissociated and triturated into single cell suspension in a solution consisted of ACSF, 1% FCS and DNAse (1µl/10mL). Trituration was carried out by using 3 glass pasteur pipets prepared at 3 different diameters. For P30 datasets, we used the Worthington Papain Dissociation System to carry out the enzymatic digestion and the cell dissociation following the manufacturer instructions.

**Single-nuclei isolation.** Dissection of the somatosensory cortex was achieved by following the same procedure as described for single-cell isolation. The dissected somatosensory cortices were transferred immediately into 500 µl of Hibernate™-E Medium (#A12476-01), then frozen for 3 minutes in isopentane pre-cooled to -80°C. The samples were subsequently stored at -80°C for long-term preservation. To process the tissue after conservation, the medium was first removed from the Eppendorf tube. Chilled 0.1X NP40 Lysis Buffer was then added in a volume of 500 µl, and the tissue was immediately homogenized using a Pellet Pestle with 15 strokes. The homogenized samples were incubated on ice for 5 minutes. The suspension was then pipette-mixed 10 times using a wide-bore pipette tip and incubate for 10min on ice to ensure proper lysis. Following lysis, 500 µl of chilled wash buffer was added to the suspension, and the mixture was pipette-mixed 5 times using a regular-bore pipette tip. The suspension was passed through a 30 µm cell strainer into a 50 ml tube to remove debris. The filtered suspension was subsequently transferred to a 1.5 ml tube for centrifugation. Samples were centrifuged at 950 × g for 10 minutes at 4°C. After centrifugation, the supernatant was carefully removed to avoid disrupting the nuclei pellet, which was retained for further analysis.

**cDNA Amplification and library construction.** 10xv3 libraries were sequenced on Illumina HiSeq 4000.

For single-nuclueus RNAseq, nuclei suspensions were adjusted following 10X recommendation. For GEM generation and barcoding, we utilized the Chromium Next GEM Single Cell 3' Reagent Kits, following the manufacturer's protocol. In summary, the prepared single-cell suspensions were loaded onto a Chromium Next GEM Chip G, along with the appropriate reagents, and processed using the chromium controller to encapsulate individual cells into GEMs. After GEM-RT incubation, cDNA was recovered and amplified through a series of cleanup and amplification steps, including SPRIselect bead-based purification. The amplified cDNA was then subjected to fragmentation, end repair, A-tailing, adaptor ligation, and sample index PCR to construct the final 3' gene expression libraries. The constructed libraries were sequenced 10xv3 libraries were sequenced on Illumina HiSeq 4000.

**Sequencing data processing.** Sequencing reads were aligned to the mouse pre-mRNA reference transcriptome (mm10) using the 10x Genomics CellRanger pipeline (version 3.1.0 or 6.1.1) with default parameters.

**External datasets.** All external datasets [1–7] used in this article were loaded as it has been made available by the authors. In the Di Bella et al. (2021) [4] dataset, cells that were annotated as low-quality cells, doublets and Red blood cells were discarded. As we have only dissected, in our experiment, the somatosensory cortex (SS CTX), we subsetted from the Allen Institute dataset only the cells coming from the primary somatosensory cortex (SSp) for the cells that they obtained with the 10x platform and from the SSp and the supplemental somatosensory cortex (SSs) for the cells obtained with the Smart-seq version 4 platform. To be sure to have clusters that were encompassed in the SS CTX, we discarded clusters that have less of 5 cells in a given subclass defined by Yao et al. (2021b) and the number of clusters in this subclass should be at least of two. Furthermore, we completed the supertype that encompassed less than 100 cells by 100 cells (when possible) passing the QC criteria (see section Below) randomly taken from the most correlated areas to the SS CTX (primary motor (MOp), secondary motor (MOs), frontal pole (FRP), auditory (AUD), gustatory (GU), visceral (VISC), visual (VIS), posterior parietal association (PTLp)) except for CR Trp73, Meis2, Astro Gfap Apoe and PVM Mrc1 supertype for which all cortical areas were used due to the little total amount of these cells. The resulting dataset from Yao et al. (2021b) [3] constitutes our reference to guide our cell assignment (called AllenRef21).

**Quality control.** To retain only high-quality cells in all datasets, cells that passed the following criteria were kept (see Supplemental information QC metrics) :

- Cells having a percentage of mitochondrial genes below or equal to 10%.
- Cells having a *$log_{10}$(number of detected genes)* within a three double median absolute deviation (doubleMAD) around the population median. The low threshold had to be greater than 500 to be applied.
- Cells having a *$log_{10}$(number of detected UMIs)* above 3 doubleMADs of the population median.
- We plotted the *$log_{10}$(nFeature RNA)* vs the *$log_{10}$(nCount RNA)* and we fitted linear regression model between these 2 variables. Cells lying below an offset of -0.09 were kept.
- We identified doublets in scRNA-seq datasets by using Scrublet [8]. The expected fraction of transcriptomes that are doublets was set to 0.1. the number of doublets to simulate, relative to the number of observed transcriptomes was set to 2. Number of neighbours used to construct the KNN classifier of observed transcriptomes and simulated doublets were set to 8.

**Assigning cells identity.** Cells that passed QC criteria were used for the following analysis. Key steps to determine the identity of the cells consisted of (Figure S1):

1. Assigning Broad class identity. An in-house artificial neural network (ANN) was used to determine the identity of the cells at the class level for E11.5 to P5 datasets. We chose Di Bella et al. (2021) [4] and Bandler et al. (2021) [2] datasets as training set for the ANN as they covered all the

broad classes that can be encountered in these studies. We pooled cells of Di Bella et al. (2021) in 5 classes: GABAergic (Interneuron), Glutamatergic (CR, UL CPN, Layer 4, DL CPN, SCPN, NP, CThPN, Layer 6b), Immature/Migrating (Immature neurons, migrating neurons), Non-Neuronal (Astrocytes, Oligodendrocytes, Microglia, Cycling glial cells, Ependymocytes, Endothelial cells, VLMC, Pericytes) and Dorsal Pallium Progenitor (Apical progenitors, Intermediate progenitors). We pooled cells of Bandler et al. (2021) in 2 classes GABAergic and SubPallium Progenitor. Weights resulting from this model (called class PAB21 model) were independently applied to datasets from E11.5 to P5 except those use as reference.

For P8 to P30 datasets we used the *map_sampling* function of the R package *scrattch.hicat* to train a centroid classifier with 80% of marker genes selected at random and match the test data to the AllenRef21 reference set at the class level (GlutNs, GABANs and NN classes). This was bootstrapped 1 000 times to get an estimate of the robustness of the classification. Cells that got a prediction probability lower than $0.5+1/(\text{number of class})^2$ were considered as undetermined.

Seurat Louvain graph-based clustering was performed on each dataset separately at the top level first (k.param=round(sqrt(Number of cells)), annoy.metric="cosine" in the FindNeighbors function and resolution=1 in the FindClusters function of the Seurat R package) and then one or more rounds of clustering is used to identify subclusters within candidate major clusters by evaluating the heterogeneity of the cluster by computing a Silhouette Score (using an adapted function of the ReclusterCells function of the R package SCISSORs [9] (Leary et al., 2021)). We assigned the identity of the cluster obtained by comparing with the prediction resulting from applying the class PAB21 model or by using *scrattch.hicat*. The most frequent predicted identity was assigned as being the cluster identity and the difference between the top predicted identity and the second most predicted identity had to be greater than 40%.

Only Glutamatergic and GABAergic predicted cells were kept for further analysis. In GE datasets, we only kept GABAergic cells as other postmitotic classes were considered as contaminant cells.

2. Determine future cortical GABANs in the ganglionic eminences datasets. Since not all cells derived from the ganglionic eminences migrate towards the cortex, we aimed to identify those that do. Using the differentially expressed (DE) branch analysis by Mayer et al. (2018) [6], we identified the common DE genes in each branch for each GE and predicted cell identities by applying the *assign_cell* function of the R package MetaMarkers [10]. We further clustered the cells using the method described above, assigning the most frequent predicted branch identity to each cluster. As branch 1 was identified as giving rise to most future cortical cells [6], we retained only cells with branch 1 identity for downstream analysis.

3. Integration of all studies. To identify homologous cell types across 10X, Drop-seq, SSv4, SSv2, C1 scRNA-seq and snRNA-seq; datasets coming from this study and external studies were integrated using Seurat's SCTransform integration workflow. 3 000 variable genes were selected for the integration analysis; k.anchor= 5 and k.filter= 150 were the parameters set for the FindIntegrationAnchors function used to identify anchors between the datasets. For Yao et al. (2021b), only the cells from the SS CTX were integrated (Figure S1c).

4. Assigning subclass and supertype identities. Datasets were split into glutamatergic and GABAergic classes.

- Glutamatergic neurons: For the E11.5 to P5 datasets, the in-house ANN was initially trained with the GlutNs part of the Di Bella et al. (2021) dataset [4] using their defined cell-type labels (referred to as the ctPA21 model). The resulting model weights were then independently applied to each dataset from E11.5 to P5. The clustering module described above was performed on the entire E11.5 to P5 dataset. Cluster identities were assigned by comparing predictions from the ctPA21 model, and the most frequent predicted identity was designated as the cluster identity.
  Correspondence between cell types annotations of Di Bella et al. (2021) and subclass annotation of Yao et al. (2021b) was achieved as followed: UL CPN, Layer 4, DL CPN: IT, SCPN: L5 PT, NP: L5 NP, CThPN: L6 CT, Layer 6b: L6b, Cajal-Retzius cells: CR.
  Next, we aimed to distinguish IT cells among L2/3 IT, L4/5 IT, L5 IT, L6 IT, or Car3 subclass identities. We used the *map_sampling* function of the scrattch.hicat package as previously described, utilizing the IT cells from the AllenRef21 dataset as the reference set.
  For P8 to P30 datasets, subclass identities were directly assigned using the *map_sampling* function of the scrattch.hicat package, as these time points are less far transcriptionally from the adult dataset. After applying our clustering module to the entire dataset, we assigned final subclass identities to clusters based on the most frequently predicted identity. To ensure accuracy, the difference between the top predicted subclass and the second most predicted subclass within each cluster had to exceed 20%. If this difference was less than 20%, we assigned a mixed subclass identity. For the adult dataset, we retained the original subclass labels.
  Finally, for each unique subclass, we assigned the supertype level of Yao et al. (2021b) [3] by using the same procedure used to assign subclass labels with the *map_sampling* function.
- GABAergic neurons: For datasets sampled from SS CTX, we assigned the subclass labels by using the same procedure than described above with scrattch.hicat. Once we obtained the subclass labels in the SS CTX datasets, we used The *MapQuery* function from the R package *Seurat* v4 (with weigh.reduction="cca" parameter) to assign the subclass labels to the datasets sampled from GEs by taking as reference the determined subclass labels from SS CTX datasets. Subclass labels were assigned by comparing to the cluster obtained with the clustering module. As the GE cells for which the subclass remained undetermined were less mature cells, we computed another round of the *MapQuery* function by taking as reference the determined subclass labels of more mature GE cells. The final subclass labels were assigned by comparing to the clusters obtained with the clustering module. We processed with the exact same procedure to determine the supertype labels for each subclass. As the Pvalb Lpl supertype (which corresponds to the Pvalb FS BC) was distributed all along the cortical thickness (Figure S2e), we deepened our analysis in order to discriminate between upper and lower Pvalb FS BC. We carried out the same procedure by using the *map_sampling* function by taking as reference the cluster level of the supertype Pvalb Lpl encompassed in our AllenRef21.

5. Assigning cell-type names. We assigned cell-type names by adding the morphology and/or the electrophysiological properties and/or the connectivity (mec-type) associated with each final identity obtained. mec-type were assigned by using patch-seq and connectivity studies [3,11–13], where they identified that many transcriptomic neuron types are correlated with known cortical neuron types. Finally, final cell-type names were annotated by pooling original cell-type names that shared

a common mec-type.

6. Classification of cell-types into families. We classified our 27 identified cell-types into 8 families: IT family encompassing L2/3 IT, L4 IT|SSC, L4/5 IT|PC, L5 IT, L6 IT cell-types; ET family encompassing L5 PT, L6 CT, L6b cell-types; Other GlutNs encompassing CR, L5/6 NP and Car3|Claustrum-like cell-types; Lamp5 encompassing Lamp5|L1 A7C/CNC and Lamp5|L1-L5 NGC cell-types; Vip encompassing Vip|L2/3-L4 BP/BTC, Vip|Uncharacterized and Vip|L2/3-L5 BP/BTC cell-types; Sst encompassing Sst|L2/3-L5 fan-MC, Sst|L4 IVC, Sst|L5 T-MC and Sst|L5/L6 NMC cell-types; Pvalb encompassing Pvalb|L2/3 CHC, Pvalb|L2/3-L4 FS BC, Pvalb|L4/5 FS BC and Pvalb|L5/6 FS BC cell-types; Other GABANs encompassing Sncg|CCK BC, Sst|FS-like and Sst Chodl|LPC cell-types.

**Pseudo-maturation score analysis.** Pseudo-maturation analysis was performed from the starting time point where a given cell-type was found until P30. For each cell-type, we first performed an integration aiming to maintain the developmental trajectory of the cells by using the R package *FastMNN* [14,15] setting the *prop.k* parameter to 0.1 for all the cell-types except those accounting a huge number of cells where *prop.k* was set to 0.4 (L2/3 IT, L6 CT). Pseudo-maturation score was calculated by first performing a k-means clustering (k=2) and then by using the R package *slingshot* (Street et al., 2018) that allowed us to fit principal curves to identify lineages within each cell-type directly on the Mutual-nearest neighbor graph resulting from the integration. Maturation directionality along the lineage was attributed by providing a starting cluster. This inferred pseudo- maturation score was further normalized between 0 and 1. For each cell-type and at each age, cells having a normalized pseudo-maturation below or above 3 doubleMADs of the population median were considered as outliers. Once outliers were discarded, the pseudo-maturation score was scaled in 3 bins spanning the ages E11.5-E17.5, E18.5-P5, and P8-P30 in the intervals [0;1/3], [1/3;2/3] and [2/3;1] respectively. Cells spanning the ages E18.5-P5 and P8-P30 were further rescaled on respectively 6 and 3 bins of equal size. Genes differentially expressed along this inferred maturation axis were identified using the *differentialGeneTest* function of the R package *Monocle 2* (v2.18.0) [16] using parameters *"fullModelFormulaStr = ˜sm.ns(pseudomaturation, df=3)+study.RNAseq.method.platform"*, and *"reducedModelFormulaStr=˜study.RNAseq.method.platform"*. We maintained genes with q-values less than 0.05 for downstream analysis. Genes with similar expression dynamics were grouped in 6 clusters using partition around medoids on the smoothed expression profiles of the significantly differentially expressed genes.

**Pseudo-layer score analysis.** Pseudo-layer analysis was performed from E18.5 to P30 by pooling the 2 nearest time points 2 by 2 from E18.5 to P5. Furthermore, this analysis was done independently for each family except for the Other GlutNs and Other GABANs families as they included cell-types that were not related to each other. For IT and ET families, at each time point defined, UMAP was computed by achieving an integration with the *Seurat* SCT workflow beforehand when necessary. For Sst, Pvalb, Vip and Lamp5 families, at each time point defined, all GABANs were used to compute the UMAP still by achieving an integration when necessary; and then families were separated by keeping the initial UMAP coordinates. Pseudo-layer score was calculated by using the same method than the pseudo-maturation described above (k was set to 3 or 2 for k-means clustering except for ET family where instead of k-means the cell-type labels were directly provided by setting L5 PT as the starting cluster and L6b as the ending cluster) with the *slingshot* package [17]. The pseudo-layer score was normalized between 0 and 1. Cells having a normalized pseudo-layer below or above 3 doubleMADs of the population median were considered as outliers. Once outliers were discarded, the pseudo-layer score was renormalized between 0 and 1. To allow both time point to time point comparison and family to family comparison, the pseudo-layer score was further scaled between values corresponding to the 5$^{th}$ percentile and the 95$^{th}$ percentile of each cell-type distribution determined in Figure S2e and Figure S3e and by adjusting the median of each cell-type to the median position determined in Figure S2e and Figure S3e that was rescaled in such a way that the 95$^{th}$ percentile corresponding to the L6b cell-type distribution was equal to 1. Genes differentially expressed along this inferred layer axis were identified using *Monocle 2* using parameters *"fullModelFormulaStr =˜sm.ns(pseudolayer, df=3)+orig.ident"*, and *"reducedModelFormulaStr=˜orig.ident"*. Genes with similar expression dynamics were grouped in 6 clusters.

**Transcriptional landscape analysis.** For each family, cells were independently embedded on a 2D graph based on their pseudo-maturation and pseudo-layer scores from E18.5 to P30. $\text{Log}_2$(CPM +1) of expression values for genes were used. We fitted to all the genes that were expressed in at least 5 cells a generalized additive model using the *gam* function from the *mgcv* R package over both the pseudo-maturation and the pseudo-layer axis. For each family, to avoid an amplified smooth expression profile, 1/3 of the total number of cells were artificially added with a $\log_2$(CPM +1) of expression values for all genes equal to 0. As pseudo-layer score was not computed for Other GlutNs and Other GABANs families, an artificial pseudo-layer score was assigned to each cell included in these families according to their relative position determined in Figure S2e. As described above, the *gam* function was applied to these 2 families. In a given family, genes were considered significantly expressed if and only if in at least one cell-type of the family and at least in one age the number of cells was at least of 5, the gene was expressed in more than 20% of the cells and its mean expression was higher or equal to the median of the median of the log2(CPM + 1) values of all cell-types in a given age. Significant genes were used to perform a PCA on their smooth expression landscape. We discretized the resulting PCA by using a k-mean clustering. K was set to 15 by using the gap statistic method. Average cluster smooth expression landscape was computed to associate a transcriptional landscape to each cluster.

**Gene ontology analysis.** We used the clusterProfiler [18,19] R package (v4.0) to find enriched biological processes in gene sets by using the enrichGO function. Gene ontology analyses were applied for each wave identified along the pseudo-maturation axis, and along the pseudo-layer axis.

**Construction of the ligand-receptor database.** LR_DB_2025 is the result of integrating and curating 109 existing public databases, to which we manually added 203 LR pairs based on literature (Table S1). To the best of our knowledge, it is the largest curated LR database. Public databases were found by using some R and python packages, in particular OmnipathR [20], singleCellSignalR [21], CellPhoneDB [22], NATMI [23], CellCall [17], scMLnet [24] and CytoTalk [25]. We tried to keep only strictly intercellular interaction type of LR. LRs referenced in LR_DB_2025 are curated by citing PMIDs matching the interaction evidence (Table S1). We attributed to all LRs *categories* and *families*. The LR category was already attributed by Omnipath for those referenced in this database; for the add-on LRs, we manually provided the category. In total, there are 20 distinct ligand categories and 10 distinct receptor categories.

The LR family was attributed by using the HUGO Gene Nomenclature Committee (HGNC) resource and to a lesser extent the Uniprot database. There are 344 distinct ligand families and 313 distinct receptor families (Table S1).

**Construction of the Transcription regulatory database.** The transcription factor (TF) database is the result of combining the merged version of mouse TRRUST v2 [26] and RegNetwork [27] "high" confidence TFs provided by the R package scSeqComm [28] and the transcription regulatory database from Omnipath by keeping only those with at least 1 reference.

**Receptor-Transcription factor a priori association.** Lists of Reactome and KEGG signaling pathways converted in directed graph provided in the GitLab associated with the scSeqComm package [28] were used to compute the score of a-priori association between a receptor and a TF using the compute tfactor PPR function of scSeqComm.

**LR landscape correlations in specific LR *categories* or LR neurodevelopmental ontologies (Fig3B).** We analyzed LR pairs where ligands showed significant transcriptional landscapes in IT neurons and receptors exhibited significant expression in GABANs. LRs belonging to neurodevelopmental ontologies were identifed using the MSigDB resource via the msigdbr R package [29,30], focusing on gene sets from the H, C2, C5, and C8 ontologies, while excluding the "cellular component (CC)" ontology. Using specific keywords, we identified LRs belonging to six neurodevelopmental ontologies: Migration (2,008 LRs), Cell Death (1,135 LRs), Differentiation (3,158 LRs), Cell Adhesion (961 LRs), Synaptogenesis (554 LRs), and Brain-associated processes (1,243 LRs), the latter encompassing nervous system functions not directly tied to development. LRs identified in these ontologies formed a new database: LR_DB_2025_Brain-Dev-Ontologies. We calculated the average correlation between ligand and receptor expression landscapes for LRs of specific categories (Fig. 3B, left plot) or neurodevelopmental ontologies (Fig. 3B, right plot), with ITs as source cells and GABANs as target cells. Statistical significance was assessed using Fisher Z-transformation.

**Inferring ligand-receptor interactions.** To infer ligand-receptor interactions between all cell type pairs, we used the R package scSeqComm 28 using log2(CPM + 1) of expression values. The interaction analysis was conducted from E18.5 to P30 by pooling the 2 nearest time points 2 by 2 from E18.5 to P5 and at the cell-type level. Only cell-types with at least 5 cells were kept for analysis. *scSeqComm* takes each LR pairs referenced in LR_DB_2025 and computes an intercellular and an intracellular score (S_inter and S_intra respectively) between all cell-types couples. *scSeqComm* computes a score between 0 and 1 for the ligand and the receptor that aimed at measuring how much the observed ligand/receptor average expression level in a given cell-type is high compared to the average expression levels observable by chance for random genes in the same cell-type. The S_inter score is equal to the minimum value obtained between the ligand and the receptor implicated in the LR pair. For a known biological signaling pathway and a given receptor, S_intra is computed and used to measure the evidence that the receptor in a given cell-type activated intracellular signaling in this specific pathway. We also computed the S_inter-diff score allowed by *scSeqComm* which represents an alternative version of the default *scSeqComm* intercellular signaling score. For each gene g in the input matrix, gene expression levels of g are normalized by the average expression level of gene g across all cell-types before computing the intercellular scores. This score aims at prioritizing ligands (receptors) that behave differently across cell-types, prioritizing LR pairs that are specific of some cell-types'interactions. To reduce false positives and increase robustness of our approach, we generated a cell-type score combining 2 scores taking into account prior knowledge about the likelihood of interaction between two *cell-types*:

- a connectivity score: We constructed a cortical column in 3D of the SS cortex including our identified cell-types. To determine the distribution of our cell types within a cortical column, we took advantage of two patch-seq studies from Gouwens et al. (2020) [11] and Scala et al. (2020) [12] and a multiplexed error-robust fluorescence in situ hybridization (MERFISH) study [31]. For each subclass, we applied the *map_sampling* function of *scrattch.hicat* to independently map cells of the 2 patch-seq studies to our reference AllenRef21 by taking the supertype level of the matching subclass (Figure S2a, Figure S2b). We assigned to each of the clusters defined in these studies the most frequent predicted supertype identity. For the MERFISH dataset, as the dataset encompassed only 254 genes, it was not optimal to find the corresponding supertype of the AllenRef21. Therefore, we took advantage of the published confusion matrix between this MERFISH dataset and 7 scRNAseq and snRNAseq datasets [13,32] (Figure S2d). For each subclass, we applied the *map_sampling* function to the 7 datasets and assigned supertype of AllenRef21 to the cluster defined in Callaway et al. (2021) [13] (Figure S2c). The confusion matrix between MERFISH dataset and the 7 datasets allowed us to find the corresponding supertype in the MERFISH dataset. We excluded clusters that were predicted into a subclass that did not correspond to their original subclass. Finally, supertype labels were merged in cell-types as we defined them in the section above to obtain the distribution of our cell-types along the cortical thickness (Figure S2e). For some cell-types we did not find correspondence in the patch-seq and MERFISH studies. We overcame this issue by assigning laminar distribution of these cell-types as follows:
  - CR: we took 100 random values encompassed in the L1 (<0.07) to reconstruct the distribution.
  - Vip|L2/3−L4 BP/BTC is a mix cell-type of the Vip Mypc1 and Vip Lmo1 supertype of the AllenRef21 thus we have pooled the cells corresponding to these 2 cell-types to reconstruct the distribution of these cell-types.

  We further applied to each "Pvalb FS BC clusters" encompassed in the 2 patch-seq and MERFISH studies, the map_sampling function of the R package scrattch.hicat by taking as reference labels the cluster levels of the AllenRef21 to determine layer specific Pvalb FS BC (Figure S3a, Figure S3b, Figure S3c). We finally pooled the obtained cluster identities of the AllenRef21 in 3 cell-types based on their corresponding distribution extrapolated from the normalized soma depth of patch-seq and MERFISH studies: Pvalb | L2/3-L4 FS BC (114 Pvalb/113 Pvalb and 114 Pvalb clusters), Pvalb | L4/5 FS BC (115 Pvalb, 116 Pvalb/115 Pvalb, and 117 Pvalb/116 Pvalb clusters), and Pvalb | L5/6 FS BC (111 Pvalb, 112 Pvalb, 116 Pvalb, 116 Pvalb/112 Pvalb, 117 Pvalb, and 119 Pvalb).

  To gain access to the morphology of our *cell-types*, morphological reconstructions in SWC format (https://download.brainimagelibrary.org/3a/88/3a88a7687ab66069/) from Scala et al. (2020) [12] were used. Only 3 of our identified cell-type did not have matched cell- type morphology: Vip Uncharacterized, Car3 Claustrum-like and CR. For Vip Uncharacterized, corresponding morphological reconstruction from Vip | L2/3-L4 BP/BTC and Vip | L2/3-L5 BP/BTC cell-types were used. Some studies suggested that Car3 Claustrum-like were IT cell-type mostly located in L6 [13,33], consequently L6 IT *cell-type* morphological reconstructions were used as Car3 Claustrum-like morphologies. For CR cells, mouse neocortex morphological reconstruction was found in the *neuromorpho.org* website (<https://neuromorpho.org/neuron info.jspneuron name=Anstoetz NC CajalRetzius 9>). This CR morphological reconstruction was rescaled in order to match the morphological reconstruction scale of Scala et al. (2020).

The constructed 3D cortical column encompassed 1 000 cells. The proportion of cell-types corresponded, as much as possible, to the proportion of each cell-type in the somatosensory cortex based on the literature [34–42] (80%GlutNs: 28.25% L2/3 IT, 23.25% L4 IT (85% L4 IT|SSC (60% strictly SSC morphology, 25% star pyramidal cell morphology (SPC)), 15% L4/5 IT|PC), 16.25% L5 (80% L5 PT, 20 % L5 IT), 29.25% L6 (85% L6 CT, 10% L6 IT, 5% L6b), 1% CR, 1% L5/6 NP, 1% Car3|Claustrum-like; 20% GABANs: 45% Pvalb (42% Pvalb|FS BC (45% Pvalb|L2/3-L4 FS BC, 29% Pvalb|L4/5 FS BC, 26% Pvalb|L5/6 FS BC), 3% Pvalb|L2/3 CHC), 28% Sst (15% Sst MC (50% Sst|L5 T-MC, 50% Sst|L2/3-L5 fan-MC), 6% Sst|L4 IVC, 5% Sst|L5/L6 NMC, 2% Sst|FS-like), 2% Sst Chodl|LPC, 10% Vip (60% Vip|L2/3-L4 BP/BTC, 20% Vip|L2/3-L5 BP/BTC, 20% Vip|Uncharacterized), 6% Sncg|CCK BC, 9% Lamp5 (44% Lamp5 L1 A7C/CNC, 56% Lamp5 L1-L5 NGC). The centroid corresponding to the position of the soma for each reconstructed cell was calculated by using the *nGauge* python package [43]. All the reconstructed cells were embedded in a 3D cortical cortical column by using the *natverse* R package [44] by positioning the soma according to the normalized cortical depth provided in [12]. If more cells than provided in[12] were needed, a random normalized cell soma depth was selected between the 5th and the 95th percentile of the cell-type distribution along the cortical column. Values of soma position were adjusted in such a way that the cortical column had a cortical depth of 1 500 µm (Y axis) and the X and Z axis had a length of 150 µm. X and Z coordinates were randomly taken between 0 and 150 for each cell. Once the 1 000 cells were embedded in the reconstructed cortical column, the potential synapses between each cell and the 999 other cells were computed by using the *potential_synapses* function of the *natverse* package that implements the method of Stepanyants and Chklovskii [45]. This method created a technical artifact by facilitating potential synapses between cells belonging to the same cell-type as they were similarly distributed within the cortical column. All the potential synapses involving 2 cells belonging to the same cell-type were assigned as NA values. A connectivity score between 2 *cell-types* was obtained as follows:

$$\forall a \in [1:999], \forall b \in [1:999],$$

$$Norm.potS_{cell_a - cell_b} = \frac{potS_{cell_a - cell_b}}{\frac{max(PotS_{cell_a}) + max(PotS_{cell_b})}{2}}$$

$where:$

$- \, a : cell \; a$

$- \, b : cell \; b$

$- \, potS : Potential \; synapses$

$$\forall j \in [1:27], \forall k \in [1:27],$$

$$Norm.potS^{ct_j \longrightarrow ct_{k_{j \neq k}}} = \frac{\sum_{j=1}^{n_{ct_j}} \sum_{k=1}^{n_{ct_k}} Norm.PotS_{cell_a - cell_b}^{ct_j \longrightarrow ct_{k_{j \neq k}}}}{n_{ct_j} + n_{ct_k}}$$

$where:$

$- \, j : cell - type \; j$

$- \, k : cell - type \; k$

$- \, n_{ct_j} : Number \; of \; cells \; belonging \; to \; cell - type \; j$

$- \, n_{ct_k} : Number \; of \; cells \; belonging \; to \; cell - type \; k$

$$\forall j \in [1:27], \forall k \in [1:27],$$

$$conn.score^{ct_j \longrightarrow ct_{k_{j \neq k}}} = \frac{Norm.potS^{ct_j \longrightarrow ct_{k_{j \neq k}}}}{\frac{max(Norm.potS^{ct_j}) + max(Norm.potS^{ct_k})}{2}}$$

For cells pairs belonging to the same *cell-type*, because a technical artifact applied, we assigned a medium connectivity score:

$$\forall j \in [1:27], \forall k \in [1:27],$$

$$conn.score^{ct_j \longrightarrow ct_{k_{j=k}}} = 0.5$$

- a development score: Some cell types colonize the cortex before others. For example, GlutNs colonize the cortex before GABANs and deep-layer GlutNs before superficial layers GlutNs. We attributed a development score between cell-types to reflect the degree of maturity of the different cell-types across cortical development. These scores span values from 0.1 to 1. (Table S9, S10, S11, S12, S13, S14, S15).

The connectivity score and the development score allowed us to define a *cell-type* score:

$$\forall j \in [1:m], \forall k \in [1:m],$$
$$c.score^{ct_j \longrightarrow ct_k} = dev.score^{ct_j \longrightarrow ct_k} \times (conn.score^{ct_j \longrightarrow ct_k} + \frac{1}{2} \times (1 - dev.score^{ct_j \longrightarrow ct_k}))$$

$where:$

$- j : cell - type\ j$

$- k : cell - type\ k$

$- m : Total\ number\ of\ cell - types\ in\ the\ processed\ dataset$

`

A ligand-receptor pair $(LR)_i$ between 2 cell-types pairs was considered significant if and only if:

$$\forall i \in [1:n], \forall j \in [1:m], \forall k \in [1:m], \forall l \in [1:M], \forall r \in [1:M],$$
$$w_{ct_{score}} \times ct_{score}^{ct_j \longrightarrow ct_k} + w_{inter_{score}} \times S\_inter_{(LR)_i}^{ct_j \longrightarrow ct_k} > w_{ct_{score}} \times (1 - ct_{score}^{ct_j \longrightarrow ct_k}) + w_{inter_{score}} \times \frac{\sum_{i=1}^{n} S\_inter_{(LR)_i}^{ct_j \longrightarrow ct_k}}{n}$$
$$\&$$
$$\bigwedge_{l=1}^{M} Pct_{L_l}^{ct_j} > 15\% \ \& \ \bigwedge_{r=1}^{M} Pct_{R_r}^{ct_k} > 15\%$$

$$\begin{aligned}
&where:\\
&-i: LR\ pair\ i\\
&-n: Total\ number\ of\ LR\ pairs\ in\ LRintercellNetworkDB\ detected\ in\ the\ processed\ dataset\\
&-l: subunit\ l\ composing\ the\ ligand\ L\\
&-r: subunit\ r\ composing\ the\ receptor\ R\\
&-M: Total\ number\ of\ subunits\ composing\ the\ ligand\ L\ or\ the\ receptor\ R\\
&-w_{ct_{score}}: weight\ applied\ to\ the\ cell-type\ score.\ If\ the\ ligand\ L\ of\ the\ (LR)_i\ was\ \boldsymbol{not\ a\ secreted}\\
&\boldsymbol{molecule}\ then\ \boldsymbol{w_{ct_{score}}=0.6};\ else\ if\ the\ ligand\ L\ of\ the\ (LR)_i\ was\ \boldsymbol{a\ secreted\ molecule}\ then\ \boldsymbol{w_{ct_{score}}=0.4}\\
&-w_{inter_{score}}: weight\ applied\ to\ the\ ligand-receptor\ interaction\ score.\ If\ the\ ligand\ L\ of\ the\ (LR)_i\ was\ \boldsymbol{not\ a\ secreted}\\
&\boldsymbol{molecule}\ then\ \boldsymbol{w_{inter_{score}}=0.4};\ else\ if\ the\ ligand\ L\ of\ the\ (LR)_i\ was\ \boldsymbol{a\ secreted\ molecule}\ then\ \boldsymbol{w_{inter_{score}}=0.6}\\
&-Pct_L^{ct_j}: Ligand\ L\ percentage\ expression\ in\ cell-type\ j\\
&-Pct_R^{ct_k}: Receptor\ R\ percentage\ expression\ in\ cell-type\ k
\end{aligned}$$

To prioritize cell-type specific interactions, we retained only significant LR pairs that had a S_inter-diff score above the population median minus 1.5 DoubleMAD for each cell-type pair and at each age.

In the case of homophilic interaction, *i.e.*, LR pairs involving the same molecule as ligand and receptor, as the mean of S_inter values could differ according to the direction of the interaction between 2 cell-types, the above threshold had to be verified in the 2 directions to consider the LR pair between 2 cell-types as significant.

To further increase the likelihood of interactions, only LR pairs that are common between 2 contiguous ages were kept.

**General analysis of inferred LR interactions (Fig. 4).** We analyzed the percentage of LR pairs associated with neurodevelopmental processes and disorders predicted by our atlas, focusing on their utilization at specific time points and between defined cell pairs. In Fig. 4C, LRs were categorized into six neurodevelopmental processes using an approach similar to the landscape analysis (Fig. 3B). However, in this case, we included all significant LRs from the LR_DB_2025 database, as determined by intercellular and cell pair scores, regardless of their landscape significance or prior classification within brain ontologies (Fig. 3B). This analysis identified 2,576 LRs associated with neuronal migration, 1,494 with neuronal cell death, 3,921 with differentiation/morphogenesis, 1,391 with cell recognition/adhesion, 613 with synaptogenesis, and 238 specific to the Cadherin family. These groups were further stratified by developmental time points (E18.5-P0 to adulthood) and cell families (Pvalb, Sst, Vip, Lamp5, and Other GABANs). We used curated neurodevelopmental disorder gene lists to identify LRs implicated in specific conditions (Fig. 4D-F). We found 29 LRs associated with epilepsy [46], 104 with intellectual disorders (from the ITHACA database: https://id-genes.orphanet.app/ithaca/), 14 with schizophrenia [47], and 100 with autism [48] (from the SFARI Gene database, accessed December 2024) (Table S5). For Fig. 4F, we visualized the disease-associated LR pairs as an interaction network using the ggraph and igraph R packages.

**ShRNAs.** RNAi experiments were conducted using shRNAs targeting the coding sequence of Mus musculus Cdh13 and Pcdh8 coding sequences (GenBank accession number NM_019707 and NM_021543) based on the following criteria (http://www.promega.com/siRNADesigner/program) :
- the sequence must start with either a Cysteine (C) or Guanine (G)
- It must have more than 50 % G or C bases
- No more than 3 consecutive base repetitions in the sequence

The sequences chosen to design oligonucleotides (Table) for shRNA genesis recognized nucleotides 1521-1541 of Cdh13 coding sequence and nucleotides 2302-2322 of Pcdh8 coding sequence. BLAST searches against Mus musculus databases confirmed the specificity of each target. As negative controls, we used corresponding non-targeting shRNAs with the same nucleotide sequence except in four positions. These shRNAs were subcloned into the mU6pro vector (gift from Dr J. LoTurco) and validated in vitro using classical western blot assays.

| **shRNA** | **Oligonucleotide sequences** |
|---|---|
| shCdh13 | F: 5’ TTTGGCAACATCAAACTATCAGGTATCAAGAGTACCTGATAGTTTGATGTTGCTTTTT3’ |
| | R: 5’ CTAGAAAAAGCAACATCAAACTATCAGGTACTCTTGATACCTGATAGTTTGATGTTGC 3’ |
| shCdh13_Ctl | F: 5’ TTTGGCAATATCAGACTATAATGTATCAAGAGTACATTATAGTCTGATATTGCTTTTT 3’ |
| | R: 5’ CTAGAAAAAGCAATATCAGACTATAATGTACTCTTGATACATTATAGTCTGATATTGC 3’ |
| shPcdh8 | F: 5’ TTTGGACCGGTTTCAGTGTTGTATATCAAGAGTATACAACACTGAAACCGGTCTTTTT 3’ |
| | R: 5’ CTAGAAAAAGACCGGTTTCAGTGTTGTATACTCTTGATATACAACACTGAAACCGGTC 3’ |
| shPcdh8_Ctl | F: 5’ TTTGTACCAGTTTCAGTATTATATATCAAGAGTATATAATACTGAAACTGGTATTTTT 3’ |
| | R:5’CTAGAAAAATACCAGTTTCAGTATTATATACTCTTGATATATAATACTGAAACTGGTA 3’ |

**In utero electroporations.** Timed pregnant C57BL6/J females were anesthetized with isoflurane (75% for induction and 2 to 2.5% for surgery) at E13.5 to trace DL neurons, at E15.5 to trace SL neurons. The uterine horns were exposed. A volume of 1–2 µL of small hairpin RNA-expressing DNA plasmid (shRNA against Cdh13 or against Pcdh8 vs their respective control shRNAs, 1.5 $\mu$g/µl) was mixed with pCAG-GFP plasmids (1 $\mu$g/$\mu$l) and Fast Green (2 mg/ml, Sigma) for further injection into the lateral ventricle of each embryo with a pulled glass capillary and a microinjector (Picospritzer II, General Valve Corporation, Fairfield, NJ, USA). Electroporation was then conducted by discharging a 4000 µF capacitor charged to 27 V using a BTX ECM 830 electroporator (BTX Harvard Apparatus, Holliston, MA, USA). Five electric pulses (5 ms duration) were delivered at 950 ms intervals using electrodes. Embryos were allowed to be born and thrive before being sacrificed at P28, and 50 $\mu$m coronal slices cut using a sliding microtome (Microm).

**Histology and immunostainings.** Mice were perfused transcardially with ice-cold 4% paraformaldehyde (in PBS). Brains were removed and post-fixed overnight at 4 °C with the same fixative. Coronal sections were cut at 50 μm thickness using a sliding microtome (Microm). Briefly, for Immunofluorescence experiments, free-floating sections were blocked and permeabilized for 2 hours in a blocking buffer composed of 10% Normal Bovine Serum, 0.2% Triton X-100 (Sigma) in PBS. Primary antibodies, diluted in blocking solution and added overnight at 4 °C, were as follows: rabbit anti-Parvalbumin (1:1000, Swant), chicken anti-GFP (1:500, Aves), rabbit anti-Synaptotagmin 2 (SYT2) (1:100, DSHB), mouse IgG2a anti-SYT2 (DSHB, 1:100), and rabbit anti-SPTBN4 (Thermofisher, 1:1000). Corresponding fluorescently labeled secondary antibodies (AlexaFluor, Invitrogen) were added for 2 h in blocking solution at room temperature. Hoechst was added in PBS for 10 min, and sections were mounted on microscope slides that were coversliped using Mowiol solution (Sigma).

**Proximity ligation assay.** 60 μm sagittal sections were treated for rabbit anti-CBLN4 (Invitrogen, PA5-36472) and anti-Mouse GLUD1 (Proteintech, 67026-1-Ig) or anti-CBLN4 and goat anti-NEO1 (Biotechne, AF1079) co-immunolabeling before application of Rabbit Plus and Mouse Minus or Rabbit Plus and Goat Minus probes, respectively (Merck, Duolink). Experiments were then done according to the manufacturer's instructions.

**Image acquisition.** Images were obtained from 50 μm thick sections using a Zeiss LSM-800 confocal microscope. Electroporated zones were imaged with a 10X objective (Plan-Apochromat, Numerical aperture 0.3) to provide an overall view of all electroporated cells. For the proximity ligation assay, we imaged layers 1 and 2 with an oil-immersed 40X objective using mosaic tiles, or with an oil-immersed 63X objective (Olympus, Numerical aperture 1.4) for GFP colocalization with puncta. For synaptic analysis, GFP+ GlutNs located in deep or superficial cortical layers were imaged with the 63X objective. A 3X digital zoom was applied to achieve a lateral and z-axis resolution of 85 nm. The Z-stack was adjusted for each slice to ensure complete imaging of the entire neuron for subsequent 3D reconstruction. Laser power and detection filter settings were optimized based on the staining quality of each slice.

**Image analysis.** All cell analyses were performed on GFP-electroporated neurons found in the DL of the S1 cortex. All images were blinded using "blind analysis tool" plugins in ImageJ. 3D-Image reconstructions and analyses were performed with IMARIS 9.9.0 software. First, a zoomed crop was done on the $GFP^+$ soma compartment. Then, to assess Pvalb FS BC cell input onto GFP-expressing electroporated cells, the $Syt2^+$ presynaptic boutons physically contacting the $GFP^+$ soma of pyramidal electroporated cells were analyzed. First, spots (diameter 0.6 μm) corresponding to individual $Syt2^+$ presynaptic boutons were created by using the "create spots" function. The $GFP^+$ soma was then reconstructed using the "create surface" tool. The density of $Syt2^+$ synaptic spots contacting the $GFP^+$ soma's surface was then measured using the object-object statistic tool with the filter "shortest distance from soma." To analyze the mean volume of presynaptic $Syt2^+$ puncta, these puncta were modelized using the "create surface" tool followed by the filter "shortest distance from soma" adjusted to 0 to isolate only the $Syt2^+$ surfaces contacting the $GFP^+$ soma surface.

**Statistics.** All statistical tests are described in the figure legends. Statistical methods to predetermine sample size were not used. Unless otherwise stated, all values represent the averages of independent experiments ± SEM. Shapiro-Wilk or Anderson-Darling test was used to test the normality of the data. Statistical significance for comparisons of one variable was determined by Student's t-test using twotailed distribution for two normally distributed groups, and by Mann-Whitney non-parametric test when distributions were not normal. For proportion comparisons, $\chi^2$ test was applied. Differences were considered significant when p-value $<0.05$. All statistical analyses were performed with R and Rstudio or with Prism 8.0.2 software (GraphPad).

**Figs. S1 to S24**

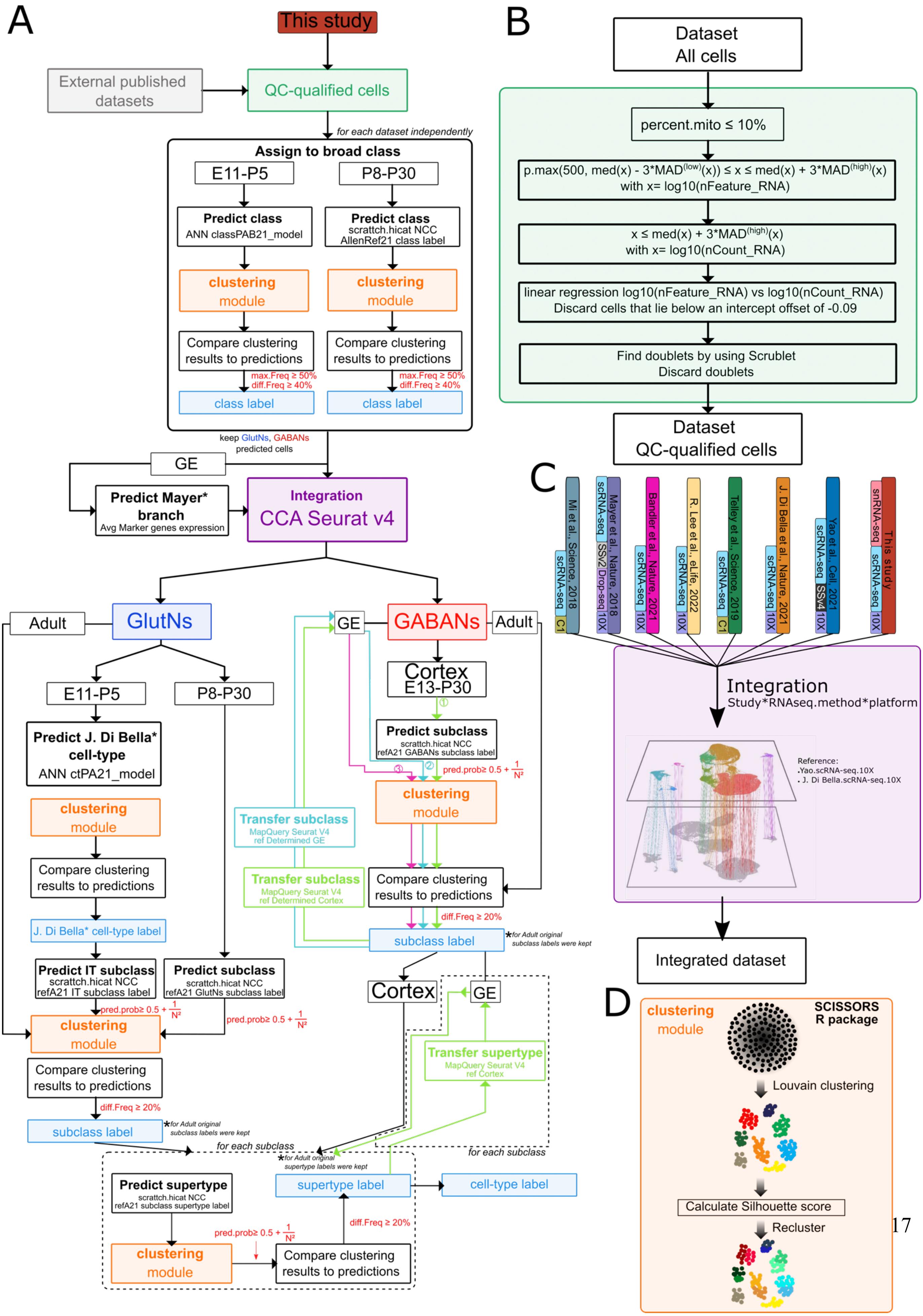
A
This study
External published datasets
QC-qualified cells
for each dataset independently
Assign to broad class
E11-P5
P8-P30
Predict class
ANN classPAB21_model
Predict class
scrattch.hicat NCC
AllenRef21 class label
clustering module
Compare clustering results to predictions
max.Freq ≥ 50%
diff.Freq ≥ 40%
class label
keep GlutNs, GABANs predicted cells
GE
Predict Mayer* branch
Avg Marker genes expression
Integration
CCA Seurat v4
GlutNs
Adult
GABANs
Predict J. Di Bella* cell-type
ANN ctPA21_model
J. Di Bella* cell-type label
Predict IT subclass
scrattch.hicat NCC
refA21 IT subclass label
Predict subclass
scrattch.hicat NCC
refA21 GlutNs subclass label
pred.prob≥ 0.5 + 1/N²
diff.Freq ≥ 20%
subclass label
*for Adult original subclass labels were kept
Cortex
E13-P30
Predict subclass
scrattch.hicat NCC
refA21 GABANs subclass label
Transfer subclass
MapQuery Seurat V4
ref Determined GE
Transfer subclass
MapQuery Seurat V4
ref Determined Cortex
Cortex
Transfer supertype
MapQuery Seurat V4
ref Cortex
for each subclass
*for Adult original supertype labels were kept
Predict supertype
scrattch.hicat NCC
refA21 subclass supertype label
supertype label
cell-type label
B
Dataset
All cells
percent.mito ≤ 10%
p.max(500, med(x) - 3*MAD^(low)(x)) ≤ x ≤ med(x) + 3*MAD^(high)(x)
with x= log10(nFeature_RNA)
x ≤ med(x) + 3*MAD^(high)(x)
with x= log10(nCount_RNA)
linear regression log10(nFeature_RNA) vs log10(nCount_RNA)
Discard cells that lie below an intercept offset of -0.09
Find doublets by using Scrublet
Discard doublets
Dataset
QC-qualified cells
C
Mi et al., Science, 2018
scRNA-seq
C1
Mayer et al., Nature, 2018
scRNA-seq
SSv2
Drop-seq
10X
Bandler et al., Nature, 2021
scRNA-seq
10X
R. Lee et al., eLife, 2022
scRNA-seq
10X
Telley et al., Science, 2019
scRNA-seq
C1
J. Di Bella et al., Nature, 2021
scRNA-seq
10X
Yao et al., Cell, 2021
scRNA-seq
SSv4
10X
This study
snRNA-seq
scRNA-seq
10X
Integration
Study*RNAseq.method*platform
Reference:
.Yao.scRNA-seq.10X
. J. Di Bella.scRNA-seq.10X
Integrated dataset
D
clustering module
SCISSORS R package
Louvain clustering
Calculate Silhouette score
Recluster

**Figure S1. scRNA-seq pipeline and analysis workflow. (a)** Pipeline of the hierarchical procedure adopted to assign the identity of the cells. ANN: Artificial Neural Network, class PAB21_model: Di Bella et al., 2021 and Bandler et al., 2021 class model; NCC: Nearest centroid classifier; AllenRef21: reference dataset from Yao et al., 2021; ctPA21 model: Di Bella et al., 2021 cell-type model; IT: Intratelencephalic; CCA: Canonical correlation analysis. **(b)** Criteria used to keep cells passing quality control from individual experimental animals scRNA-seq data. **(c)** Integration of all the compiled datasets by using the SCT workflow of *Seurat* by taking the Yao et al, 2021 and J. Di Bella et al., 2021 datasets as reference. **(d)** Several rounds of Louvain clustering were performed by using the *SCISSOR* R package.

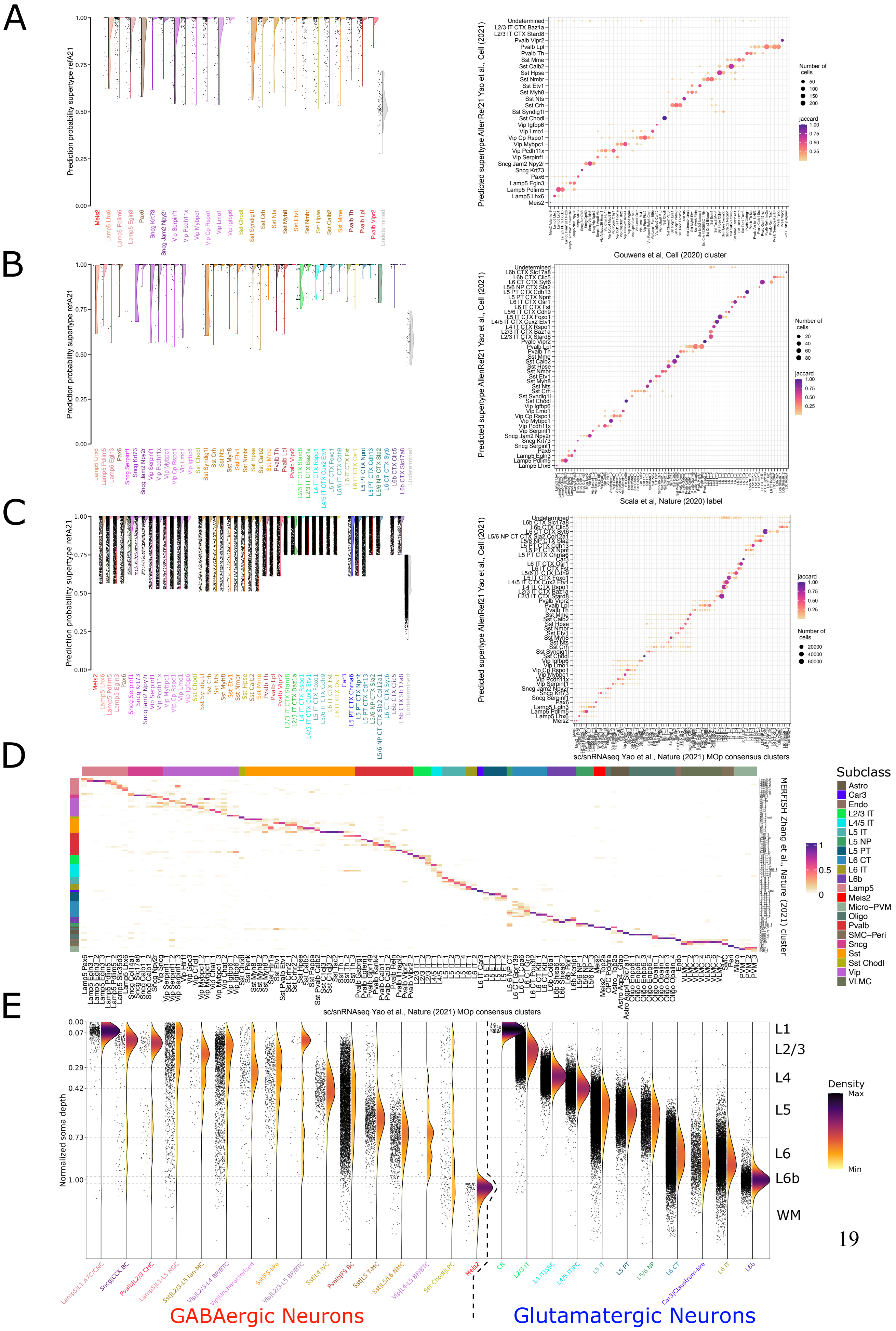
A
Prediction probability supertype refA21
Predicted supertype AllenRef21 Yao et al., Cell (2021)
Gouwens et al, Cell (2020) cluster
Number of cells
jaccard
B
Scala et al, Nature (2020) label
C
sc/snRNAseq Yao et al., Nature (2021) MOp consensus clusters
D
MERFISH Zhang et al., Nature (2021) cluster
Subclass
Astro
Car3
Endo
L2/3 IT
L4/5 IT
L5 IT
L5 NP
L5 PT
L6 CT
L6 IT
L6b
Lamp5
Meis2
Micro–PVM
Oligo
Pvalb
SMC–Peri
Sncg
Sst
Sst Chodl
Vip
VLMC
E
Normalized soma depth
L1
L2/3
L4
L5
L6
L6b
WM
Density
Max
Min
GABAergic Neurons
Glutamatergic Neurons

**Figure S2. Correspondence between supertype of the AllenRef21 reference and cell-types defined in patch-seq and MERFISH studies.** Mapping results of Gouwens et al., 2020 patch-seq study (a), Scala et al., 2020 patch-seq study (b), and Yao et al., 2021 (c) on the AllenRef21 by using the nearest centroid classifier (NCC) of the *scrattch.hicat* R package. Left: Prediction probability resulting from the mapping performed with the NCC. Right: annotation comparisons resulting from the mapping of the different studies on the AllenRef21. The size of the dots indicates the number of overlapping cells, and the color indicates the Jaccard index (number of cells in intersection/number of cells in union). (d) Correlation matrix provided in BICCN study (Extended data figure 5) that gives the correspondence between MOp consensus cluster taxonomy and MERFISH data. (e) Distribution of the different cell-types in the cortex extrapolating from data of normalized soma depth provided in Gouwens et al., 2020, Scala et al., 2020 and Zhang et al., 2021.

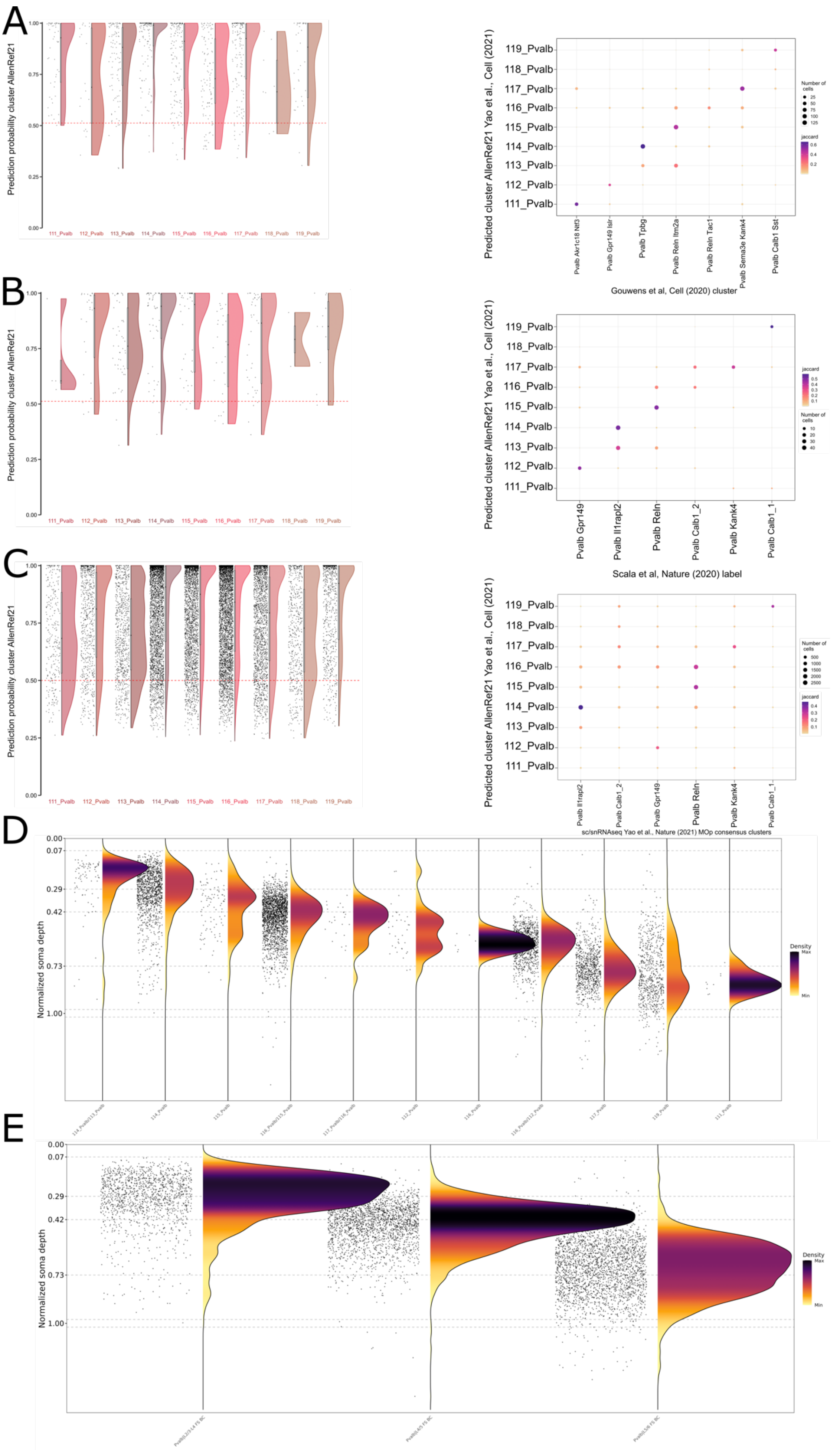

A
B
C
D
E
Prediction probability cluster AllenRef21
Predicted cluster AllenRef21 Yao et al., Cell (2021)
119_Pvalb
118_Pvalb
117_Pvalb
116_Pvalb
115_Pvalb
114_Pvalb
113_Pvalb
112_Pvalb
111_Pvalb
Gouwens et al, Cell (2020) cluster
Pvalb Gpr149
Pvalb Il1rapl2
Pvalb Reln
Pvalb Calb1_2
Pvalb Kank4
Pvalb Calb1_1
Scala et al, Nature (2020) label
sc/snRNAseq Yao et al., Nature (2021) MOp consensus clusters
Number of cells
jaccard
Normalized soma depth
0.00
0.07
0.29
0.42
0.73
1.00
Density
Max
Min

**Figure S3. Correspondence between Pvalb cluster of the AllenRef21 reference and ”Pvalb FS BC” cell- types defined in patch-seq and MERFISH studies.** Mapping results of ”Pvalb FS BC” cell-types from Gouwens et al., 2020 patch-seq study (a), Scala et al., 2020 patch-seq study (b), and Yao et al.,Nature, 2021 (c) on the AllenRef21 by using the nearest centroid classifier (NCC) of the scrattch.hicat R package by taking the cluster labels as reference. Left: Prediction probability resulting from the mapping performed with the NCC. Right: annotations comparison resulting from the mapping of the different studies on the AllenRef21. The size of the dots indicates the number of overlapping cells, and the color indicates the Jaccard index (number of cells in intersection/number of cells in union). (d) Distribution of the different Pvalb clusters of the AllenRef21 in the cortex extrapolating from data of normalized soma depth provided in Gouwens et al., 2020, Scala et al., 2020 and Zhang et al., 2021. (e) Clusters in (d) were pooled in 3 cell-types: Pvalb L2/3-L4 FS BC, Pvalb L4/5 FS BC, Pvalb L5/6 FS BC. Distribution of the different defined Pvalb cell-types are plotted.

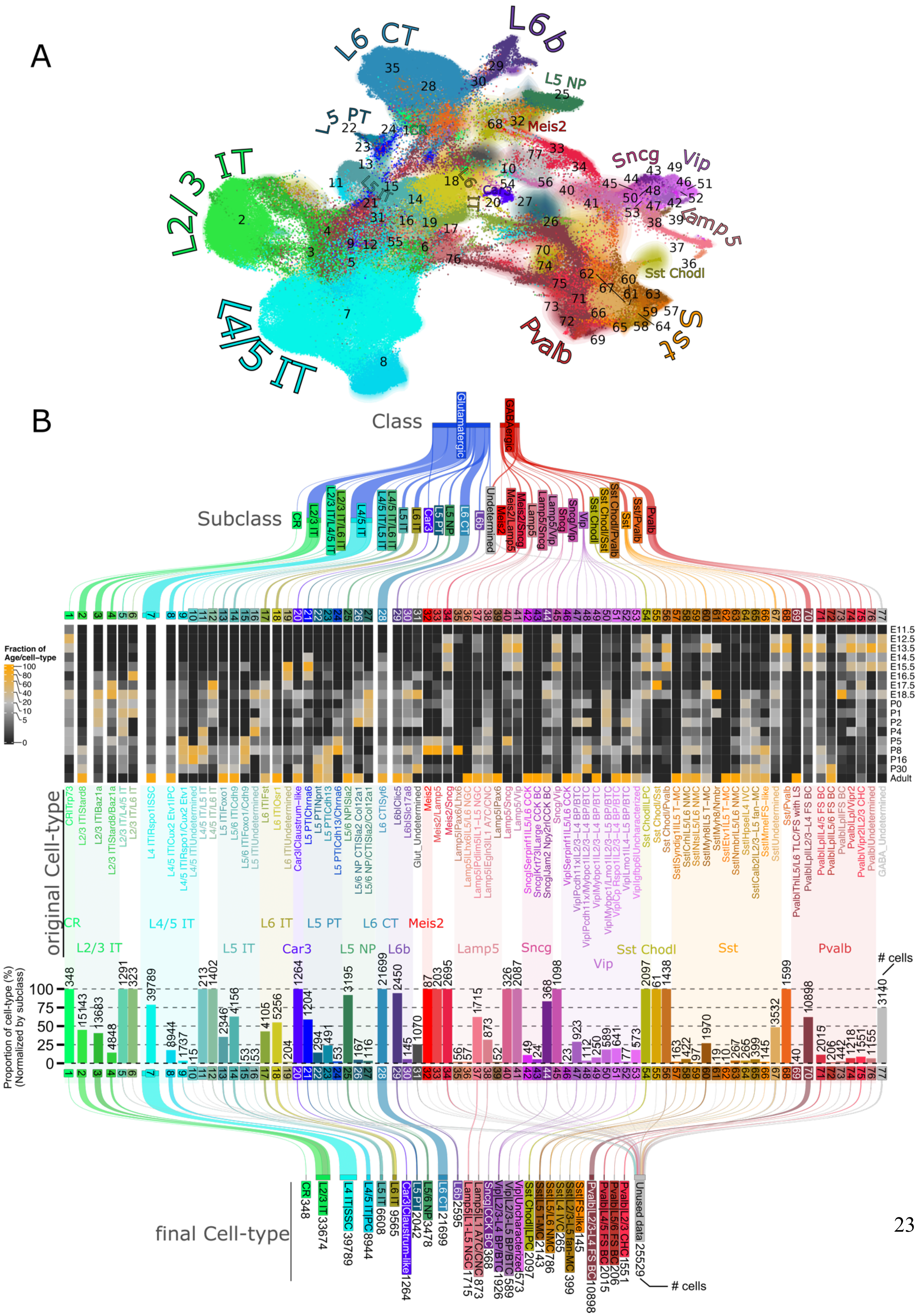
A
L6 CT
L6b
L5 NP
L5 PT
Meis2
L2/3 IT
Sncg
Vip
Lamp5
Sst Chodl
L4/5 IT
Pvalb
Sst
B
Class
Glutamatergic
GABAergic
Subclass
original Cell-type
final Cell-type
Fraction of Age/cell-type
Proportion of cell-type (%) (Normalized by subclass)
### cells

**Figure S4. Assignment of final cell-types from original cell-types assignment. (a)** UMAP visualization of scRNA-seq and snRNA-seq data after integration. Cells are coloured by original cell-types assignment. **(b)** Simplification of the number of original cell-types found by pooling original cell-types sharing the same morphological, electrophysiological and connectivity type. From top to bottom: Sankey diagram depicting the different hierarchical levels of cell-type identities; heatmap representing the fraction of age per cell-type; bar plot representing the proportion of original cell-types in each subclass; Sankey diagram depicting the grouping of original cell-types in final cell-types that were used for downstream analyses.

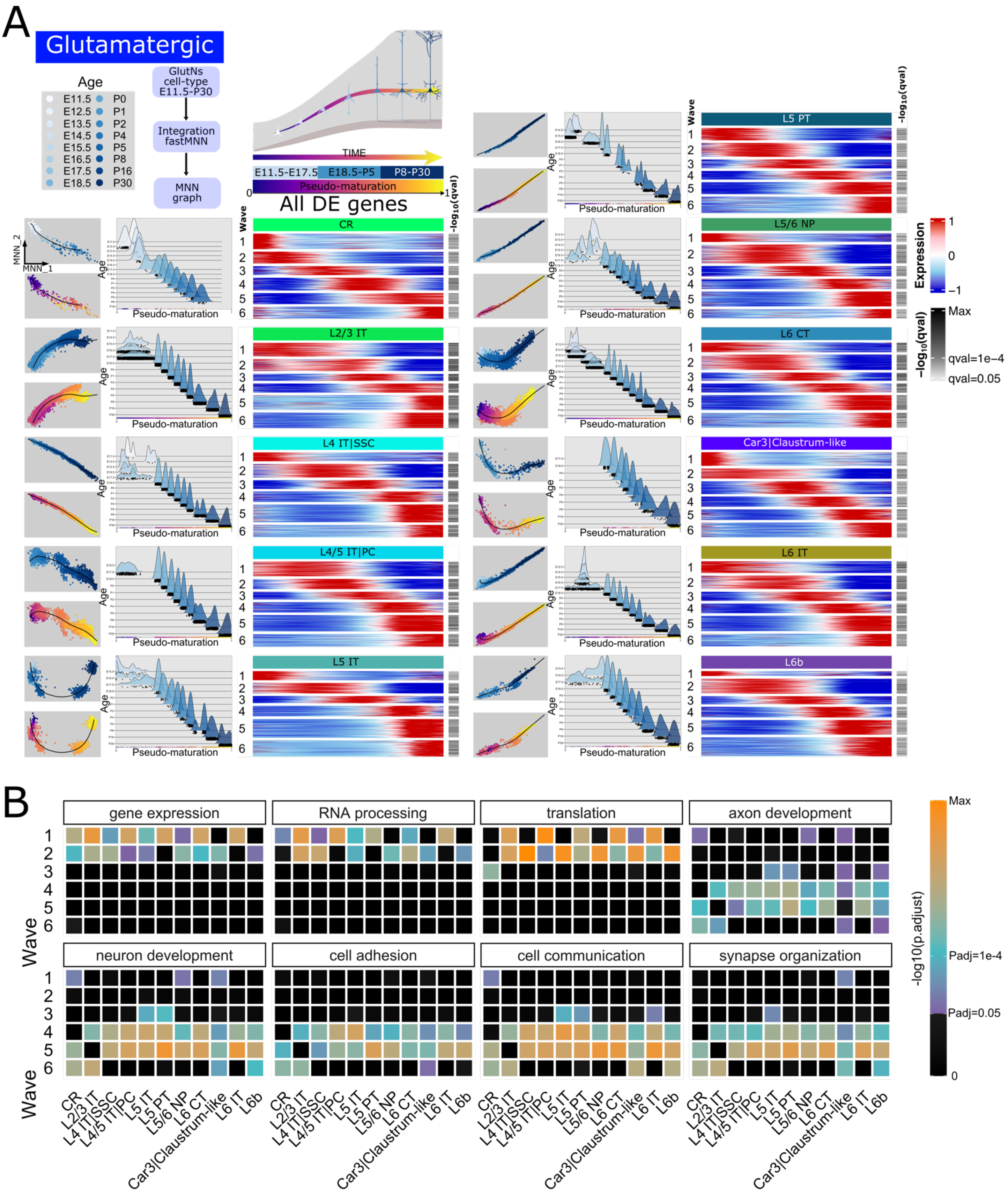
A
Glutamatergic
Age
E11.5 P0
E12.5 P1
E13.5 P2
E14.5 P4
E15.5 P5
E16.5 P8
E17.5 P16
E18.5 P30
GlutNs cell-type E11.5-P30
Integration fastMNN
MNN graph
TIME
E11.5-E17.5
E18.5-P5
P8-P30
Pseudo-maturation
All DE genes
MNN_2
MNN_1
Age
Wave
-log10(qval)
CR
L2/3 IT
L4 IT|SSC
L4/5 IT|PC
L5 IT
L5 PT
L5/6 NP
L6 CT
Car3|Claustrum-like
L6 IT
L6b
Expression
-log10(qval)
Max
qval=1e-4
qval=0.05
B
gene expression
RNA processing
translation
axon development
neuron development
cell adhesion
cell communication
synapse organization
Wave
-log10(p.adjust)
Max
Padj=1e-4
Padj=0.05
0

**Figure S5. Transcriptional dynamics of glutamatergic neuron cell-types during somatosensory cortex development. (a)** Transcriptional dynamics of glutamatergic neuron cell-types along a pseudo-maturation axis. left: Mutual nearest-neighbor plot obtained after fastMNN integration showing chronotopic organization along a pseudo-maturation axis. Black line indicates pseudo-maturation axis onto which cells were aligned. right: Heatmap illustrating the expression of the differentially expressed (DE) genes along the pseudo-maturation axis. Genes with similar patterns were grouped in waves (numbers). **(b)** Examples of gene ontology processes associated with each of the expression waves for each glutamatergic neuron cell-types.

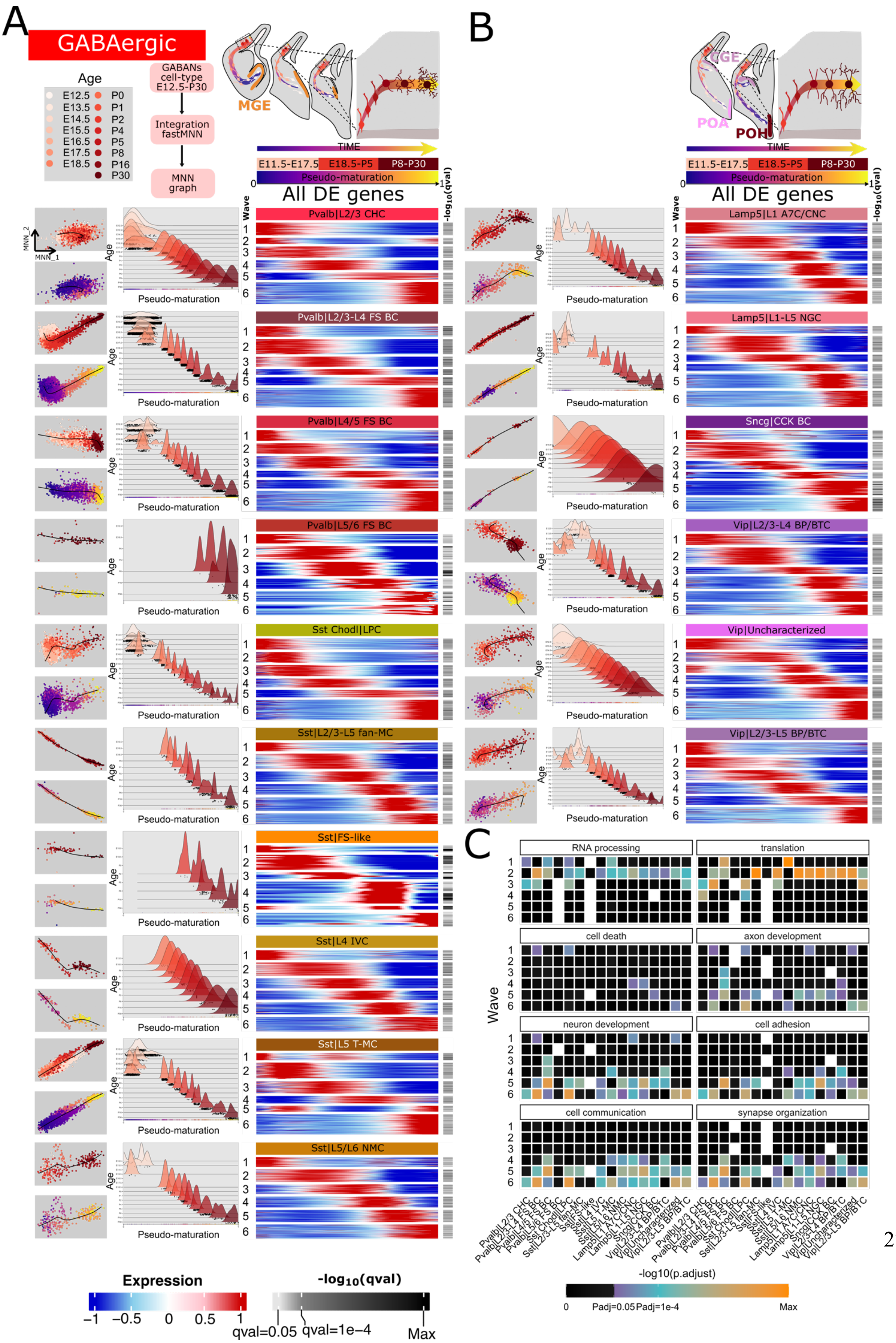
A
GABAergic
Age
E12.5 E13.5 E14.5 E15.5 E16.5 E17.5 E18.5
P0 P1 P2 P4 P5 P8 P16 P30
GABANs cell-type E12.5-P30
Integration fastMNN
MNN graph
MGE
TIME
E11.5-E17.5 E18.5-P5 P8-P30
Pseudo-maturation
All DE genes
Wave
-log10(qval)
MNN_2
MNN_1
Age
Pseudo-maturation
Pvalb|L2/3 CHC
Pvalb|L2/3-L4 FS BC
Pvalb|L4/5 FS BC
Pvalb|L5/6 FS BC
Sst Chodl|LPC
Sst|L2/3-L5 fan-MC
Sst|FS-like
Sst|L4 IVC
Sst|L5 T-MC
Sst|L5/L6 NMC
B
CGE
POA
POH
Lamp5|L1 A7C/CNC
Lamp5|L1-L5 NGC
Sncg|CCK BC
Vip|L2/3-L4 BP/BTC
Vip|Uncharacterized
Vip|L2/3-L5 BP/BTC
C
RNA processing
translation
cell death
axon development
neuron development
cell adhesion
cell communication
synapse organization
Wave
-log10(p.adjust)
0 Padj=0.05 Padj=1e-4 Max
Expression
−1 −0.5 0 0.5 1
-log10(qval)
qval=0.05 qval=1e−4 Max

**Figure S6. Transcriptional dynamics of GABAergic neuron cell-types during somatosensory cortex development. (a)** Transcriptional dynamics of MGE-derived GABAergic neuron cell-types along a pseudo-maturation axis. left: Mutual nearest-neighbor plot obtained after fastMNN integration showing chronotopic organization along a pseudo-maturation axis. Black line indicates pseudo-maturation axis onto which cells were aligned. right: Heatmap illustrating the expression of the differentially expressed (DE) genes along the pseudo-maturation axis. Genes with similar patterns were grouped in waves (numbers). **(b)** Same analysis than in **(a)** but for CGE/POA/POH-derived GABAergic neurons. **(c)** Examples of gene ontology processes associated with each of the expression waves for each GABAergic neuron cell-types.

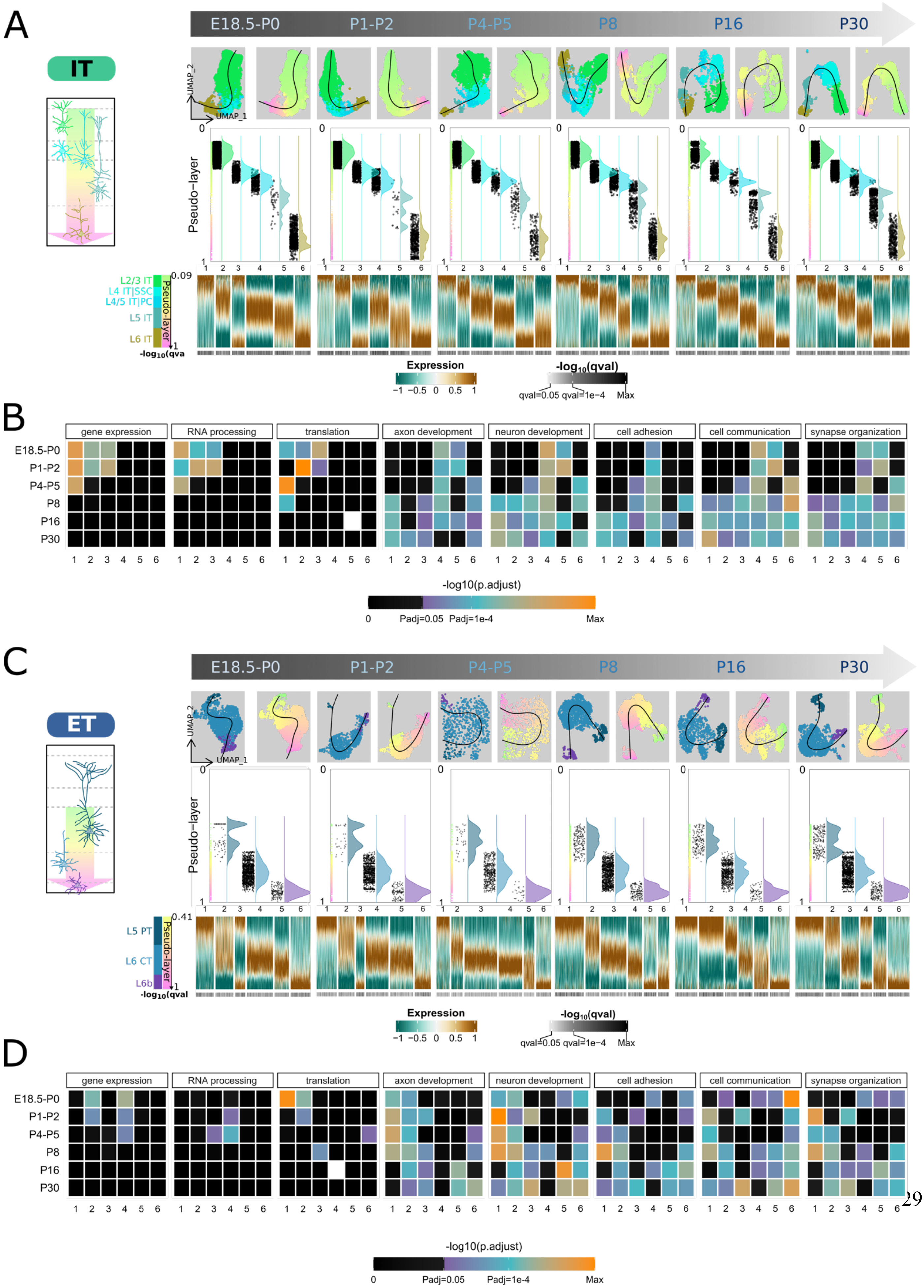
A
E18.5-P0
P1-P2
P4-P5
P8
P16
P30
IT
UMAP_2
UMAP_1
Pseudo−layer
L2/3 IT
L4 IT|SSC
L4/5 IT|PC
L5 IT
L6 IT
0.09
Expression
−1 −0.5 0 0.5 1
-log10(qval)
qval=0.05 qval=1e−4 Max
B
gene expression
RNA processing
translation
axon development
neuron development
cell adhesion
cell communication
synapse organization
-log10(p.adjust)
0 Padj=0.05 Padj=1e-4 Max
C
ET
L5 PT
L6 CT
L6b
0.41
D

**Figure S7. Continuous distribution of IT and ET glutamatergic neurons along the cortical depth from E18.5 to P30. (a)** Continuous distribution of IT glutamatergic neuron cell-types along a pseudo-layer axis. top: UMAP embedding obtained after *Seurat* SCT workflow integration showing spontaneous spatial organization along a pseudo-layer axis (middle). Black line indicates pseudo-layer axis onto which cells were aligned. right: Heatmap illustrating the expression of the differentially expressed (DE) genes along the pseudo-layer axis. Genes with similar patterns were grouped in waves (numbers). **(b)** Examples of gene ontology processes associated with each of the expression waves for each age. **(c)** Continuous distribution of ET glutamatergic neuron cell-types along a pseudo-layer axis. top: UMAP embedding obtained after *Seurat* SCT workflow integration showing spontaneous spatial organization along a pseudo-layer axis (middle). Black line indicates pseudo-layer axis onto which cells were aligned. right: Heatmap illustrating the expression of the differentially expressed (DE) genes along the pseudo-layer axis. Genes with similar patterns were grouped in waves (numbers). **(d)** Examples of gene ontology processes associated with each of the expression waves for each age.

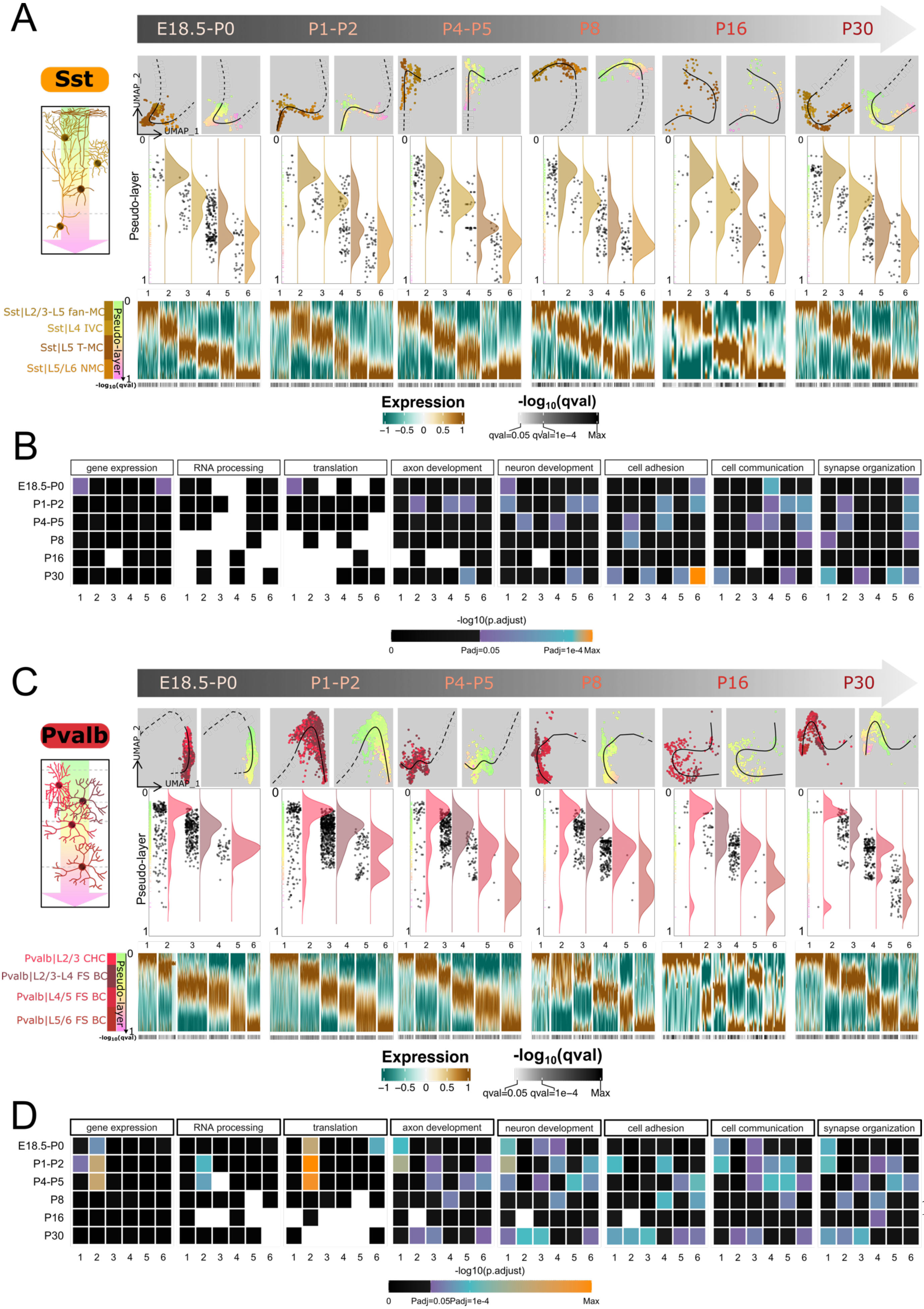
A
E18.5-P0
P1-P2
P4-P5
P8
P16
P30
Sst
UMAP_2
UMAP_1
Pseudo-layer
Sst|L2/3-L5 fan-MC
Sst|L4 IVC
Sst|L5 T-MC
Sst|L5/L6 NMC
-log10(qval)
Expression
-log10(qval)
qval=0.05 qval=1e−4 Max
B
gene expression
RNA processing
translation
axon development
neuron development
cell adhesion
cell communication
synapse organization
E18.5-P0
P1-P2
P4-P5
P8
P16
P30
-log10(p.adjust)
Padj=0.05
Padj=1e-4 Max
C
E18.5-P0
P1-P2
P4-P5
P8
P16
P30
Pvalb
UMAP_2
UMAP_1
Pseudo-layer
Pvalb|L2/3 CHC
Pvalb|L2/3-L4 FS BC
Pvalb|L4/5 FS BC
Pvalb|L5/6 FS BC
-log10(qval)
Expression
-log10(qval)
qval=0.05 qval=1e−4 Max
D
gene expression
RNA processing
translation
axon development
neuron development
cell adhesion
cell communication
synapse organization
E18.5-P0
P1-P2
P4-P5
P8
P16
P30
-log10(p.adjust)
Padj=0.05Padj=1e-4
Max

1

**Figure S8. Continuous distribution of Sst and Pvalb GABAergic neurons along the cortical depth from E18.5 to P30. (a)** Continuous distribution of Sst GABAergic neuron cell-types along a pseudo-layer axis. top: UMAP embedding obtained after *Seurat* SCT workflow integration showing spontaneous spatial organization along a pseudo-layer axis (middle). Black line indicates pseudo-layer axis onto which cells were aligned. right: Heatmap illustrating the expression of the differentially expressed (DE) genes along the pseudo-layer axis. Genes with similar patterns were grouped in waves (numbers). **(b)** Examples of gene ontology processes associated with each of the expression waves for each age. **(c)** Continuous distribution of Pvalb GABAergic neuron cell-types along a pseudo-layer axis. top: UMAP embedding obtained after *Seurat* SCT workflow integration showing spontaneous spatial organization along a pseudo-layer axis (middle). Black line indicates pseudo-layer axis onto which cells were aligned. right: Heatmap illustrating the expression of the differentially expressed (DE) genes along the pseudo-layer axis. Genes with similar patterns were grouped in waves (numbers). **(d)** Examples of gene ontology processes associated with each of the expression waves for each age.

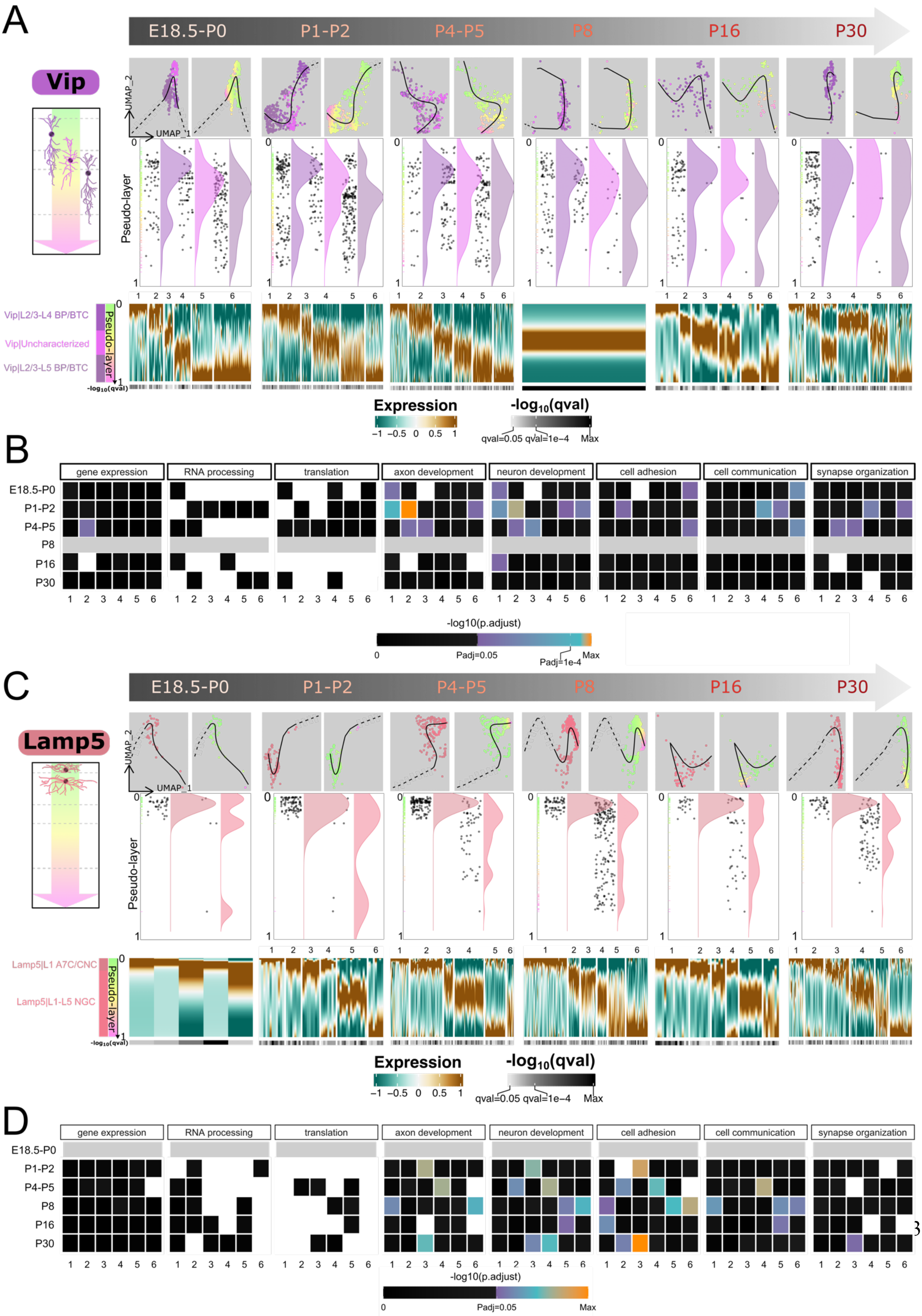
A
E18.5-P0
P1-P2
P4-P5
P8
P16
P30
Vip
UMAP_2
UMAP_1
Pseudo-layer
Vip|L2/3-L4 BP/BTC
Vip|Uncharacterized
Vip|L2/3-L5 BP/BTC
-log10(qval)
Expression
-1 -0.5 0 0.5 1
-log10(qval)
qval=0.05 qval=1e-4 Max
B
gene expression
RNA processing
translation
axon development
neuron development
cell adhesion
cell communication
synapse organization
-log10(p.adjust)
0
Padj=0.05
Padj=1e-4
Max
C
Lamp5
Lamp5|L1 A7C/CNC
Lamp5|L1-L5 NGC
D
Padj=0.05

**Figure S9. Continuous distribution of Vip and Lamp5 GABAergic neurons along the cortical depth from E18.5 to P30. (a)** Continuous distribution of Vip GABAergic neuron cell-types along a pseudo-layer axis. top: UMAP embedding obtained after *Seurat* SCT workflow integration showing spontaneous spatial organization along a pseudo-layer axis (middle). Black line indicates pseudo-layer axis onto which cells were aligned. right: Heatmap illustrating the expression of the differentially expressed (DE) genes along the pseudo-layer axis. Genes with similar patterns were grouped in waves (numbers). **(b)** Examples of gene ontology processes associated with each of the expression waves for each age. **(c)** Continuous distribution of Lamp5 GABAergic neuron cell-types along a pseudo-layer axis. top: UMAP embedding obtained after *Seurat* SCT workflow integration showing spontaneous spatial organization along a pseudo-layer axis (middle). Black line indicates pseudo-layer axis onto which cells were aligned. right: Heatmap illustrating the expression of the differentially expressed (DE) genes along the pseudo-layer axis. Genes with similar patterns were grouped in waves (numbers). **(d)** Examples of gene ontology processes associated with each of the expression waves for each age.

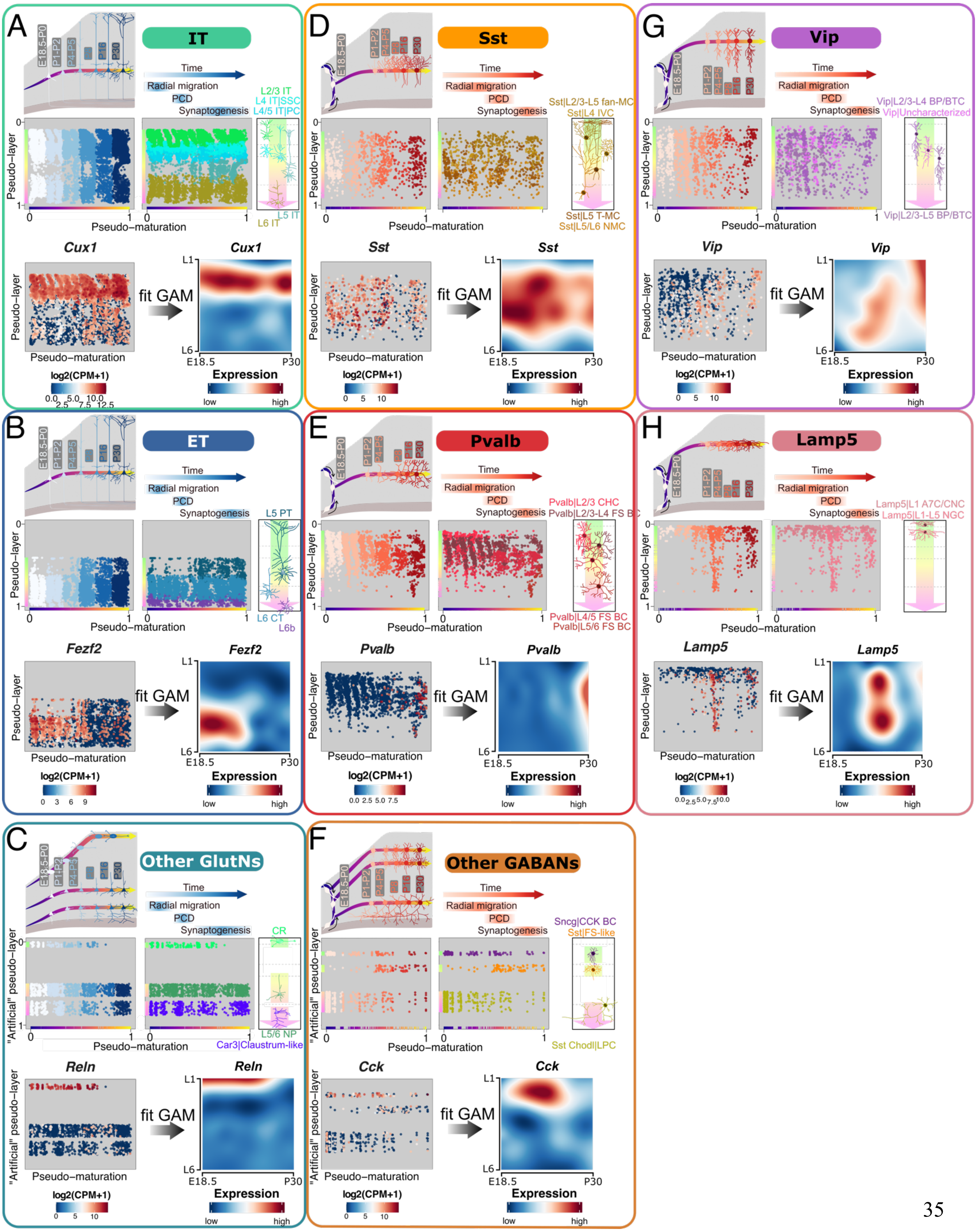
A
IT
Time
Radial migration
PCD
Synaptogenesis
L2/3 IT
L4 IT|SSC
L4/5 IT|PC
L5 IT
L6 IT
Pseudo–layer
Pseudo–maturation
Cux1
fit GAM
log2(CPM+1)
Expression
low
high
E18.5
P30
L1
L6
B
ET
L5 PT
L6 CT
L6b
Fezf2
C
Other GlutNs
CR
L5/6 NP
Car3|Claustrum-like
"Artificial" pseudo–layer
Reln
D
Sst
Sst|L2/3-L5 fan-MC
Sst|L4 IVC
Sst|L5 T-MC
Sst|L5/L6 NMC
E
Pvalb
Pvalb|L2/3 CHC
Pvalb|L2/3-L4 FS BC
Pvalb|L4/5 FS BC
Pvalb|L5/6 FS BC
F
Other GABANs
Sncg|CCK BC
Sst|FS-like
Sst Chodl|LPC
Cck
G
Vip
Vip|L2/3-L4 BP/BTC
Vip|Uncharacterized
Vip|L2/3-L5 BP/BTC
H
Lamp5
Lamp5|L1 A7C/CNC
Lamp5|L1-L5 NGC
E18.5-P0
P1-P2
P4-P5
P16
P30

**Figure S10. Spatiotemporal transcriptional dynamics.** For each family (one panel per family). Top: 2D maps in which cells are embedded according to their pseudo-maturation (x-axis) and pseudo-layer (y-axis) scores. Bottom: A generalized additive model (GAM) was applied to generate 2D maps for the spatiotemporal expression of genes (”transcriptional landscape”) throughout somatosensory cortex development. Exemple of one single gene for each family is represented. GlutN (A) and GABAN (B) families are represented.

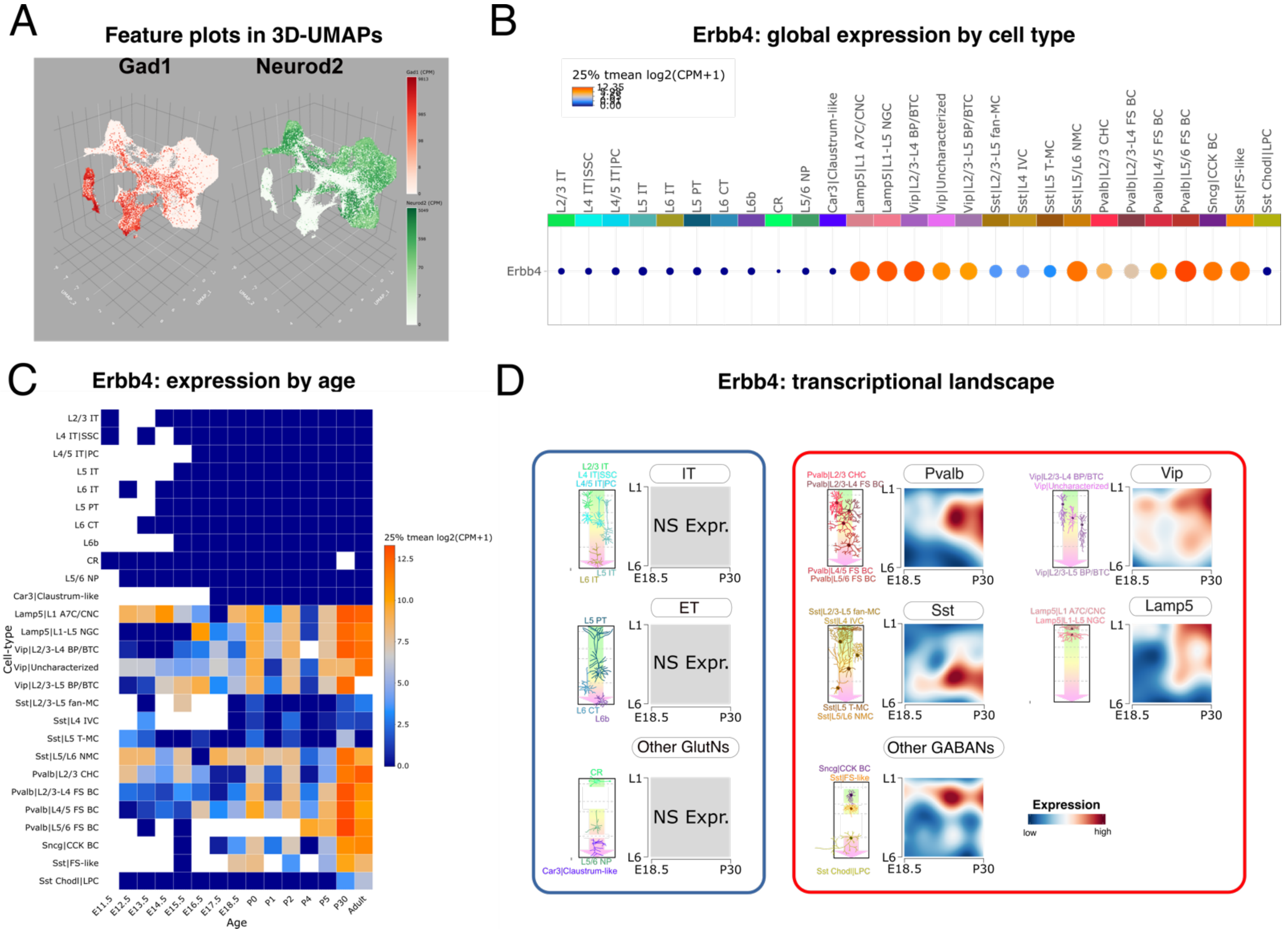


**Figure S11: Gene expression explorations with the shiny app scLRSomatodev.** (A) Feature plot for two example genes (Gad1 and Neurod2) on 3D-UMAP of all cells from 17 ages. (B) Dot plot of absolute gene expression per cell-type (grouping of all ages). Circle size reflects percentage of cells expressing Erbb4 per cell-type. (C) Heat map of Erbb4 absolute expression (25 % trimmed mean of log2(CPM +1) ) over mouse age in all 27 neuronal cell-types. (D) Transcriptional landscapes of Erbb4 expression in each GlutN (left, blue contour) and GABAN (right, red contour) families.

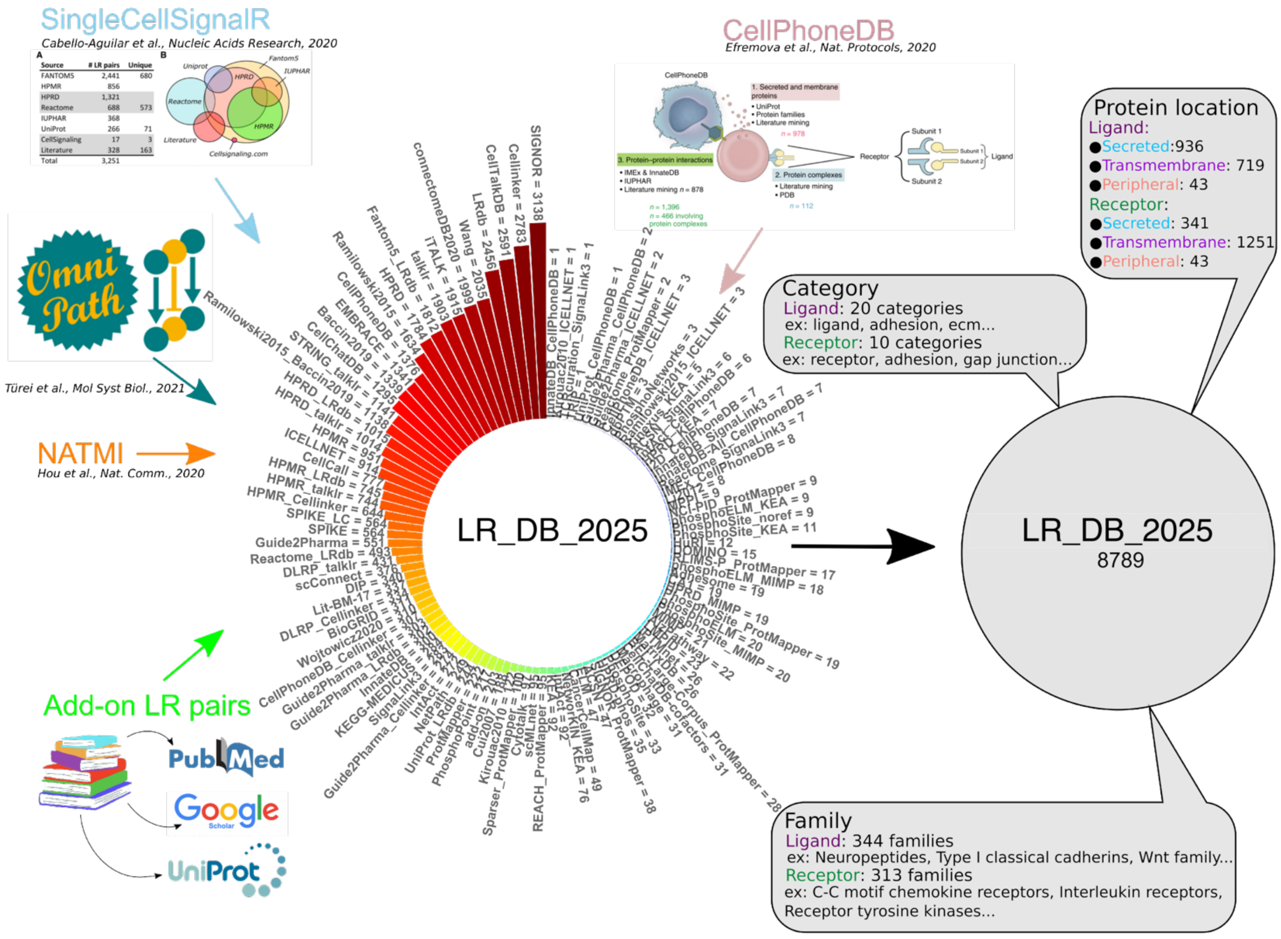


**Fig. S12. The ligand-receptor database.**

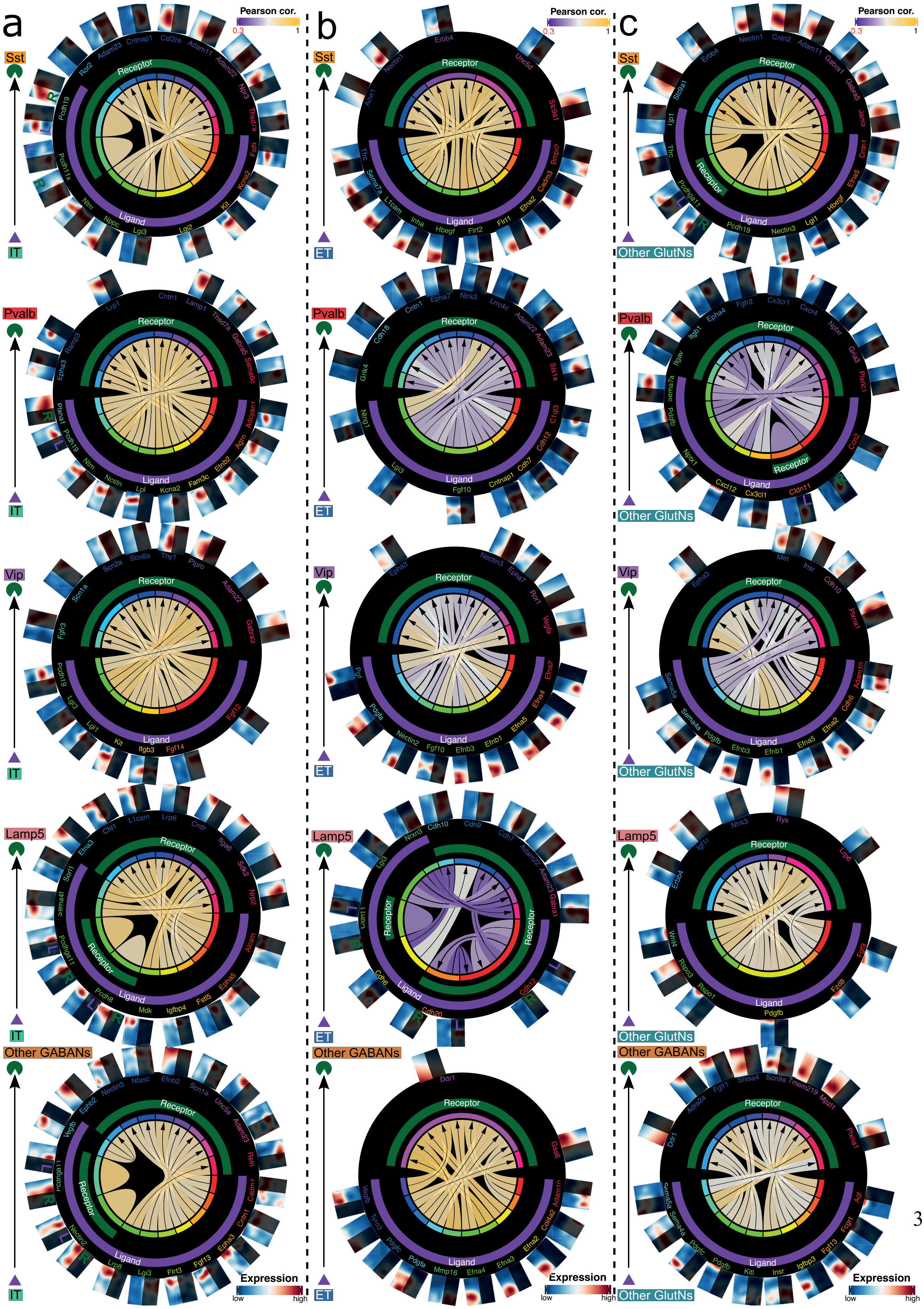
a
b
c
Pearson cor.
0.3
1
Sst
Pvalb
Vip
Lamp5
Other GABANs
IT
ET
Other GlutNs
Receptor
Ligand
Expression
low
high

**Figure S13. ”GABAergic neuron lamination process” transcriptional landscapes correlation of ligand-receptor pairs between glutamatergic expressing ligand and GABAergic expressing receptor families.** (a) Chord diagram representing the correlation between transcriptional landscapes of IT expressing ligand and GABAN families expressing receptor. (b) Chord diagram representing the correlation between transcriptional landscapes of ET expressing ligand and GABAN families expressing receptor. (c) Chord diagram representing the correlation between transcriptional landscapes of Other GlutNs expressing ligand and GABAN families expressing receptor. Only the top 10 most correlated LR pairs are represented between each family couple. Pearson correlation values were calculated based on the non-shaded area of the transcriptional landscapes (Pseudo-maturation values from 0 to 0.5 were used).

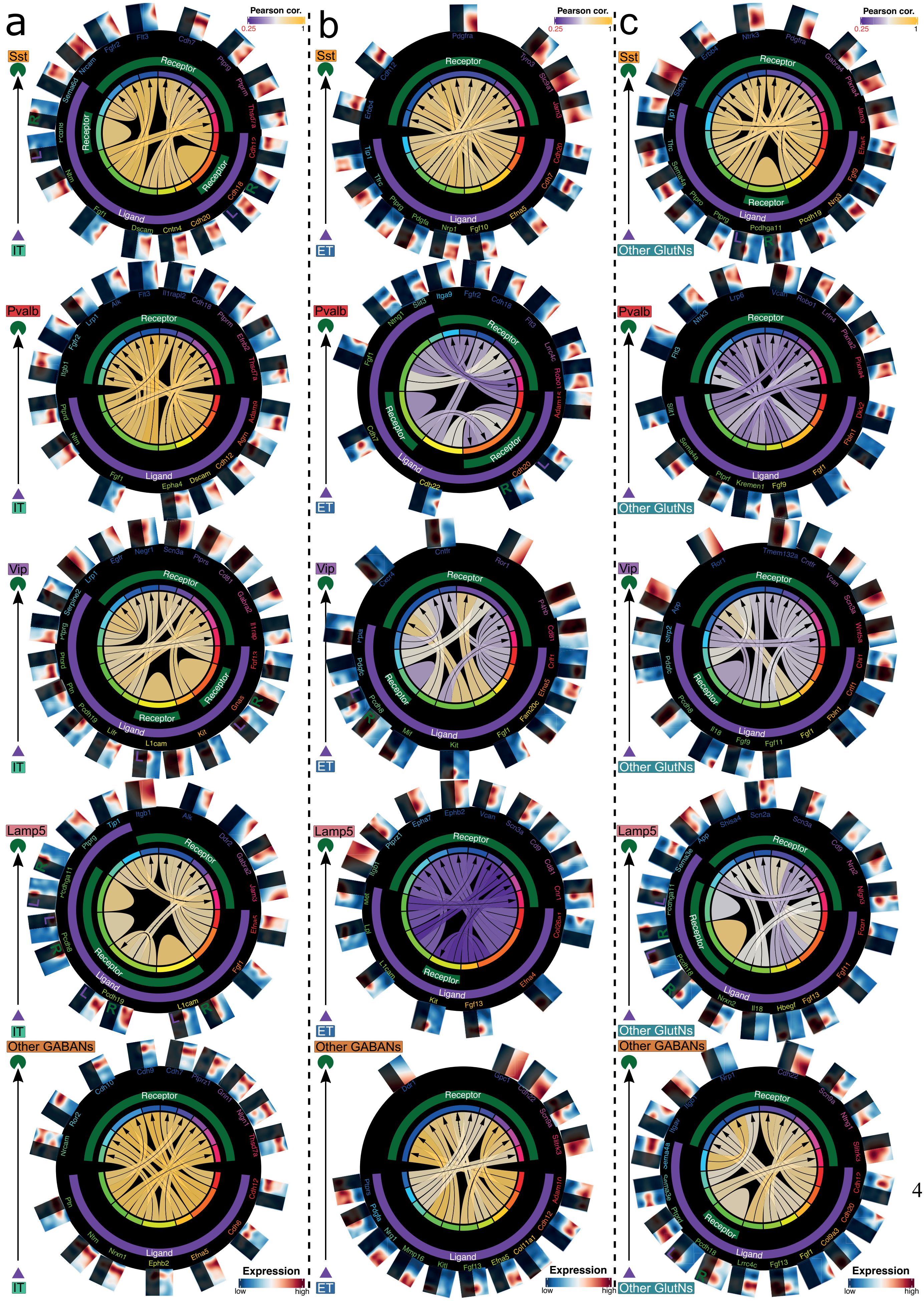

a
b
c
Pearson cor.
0.25
1
Sst
Pvalb
Vip
Lamp5
Other GABANs
IT
ET
Other GlutNs
Receptor
Ligand
Expression
low
high

**Figure S14. ”GABAergic neuron synaptogenesis process” transcriptional landscapes correlation of ligand-receptor pairs between glutamatergic expressing ligand and GABAergic expressing receptor families.** (a) Chord diagram representing the correlation between transcriptional landscapes of IT expressing ligand and GABAN families expressing receptor. (b) Chord diagram representing the correlation between transcriptional landscapes of ET expressing ligand and GABAN families expressing receptor. (c) Chord diagram representing the correlation between transcriptional landscapes of Other GlutNs expressing ligand and GABAN families expressing receptor. Only the top 10 most correlated LR pairs are represented between each family couple. Pearson correlation values were calculated based on the non-shaded area of transcriptional landscapes (Pseudo-maturation values from 0.5 to 1 were used).

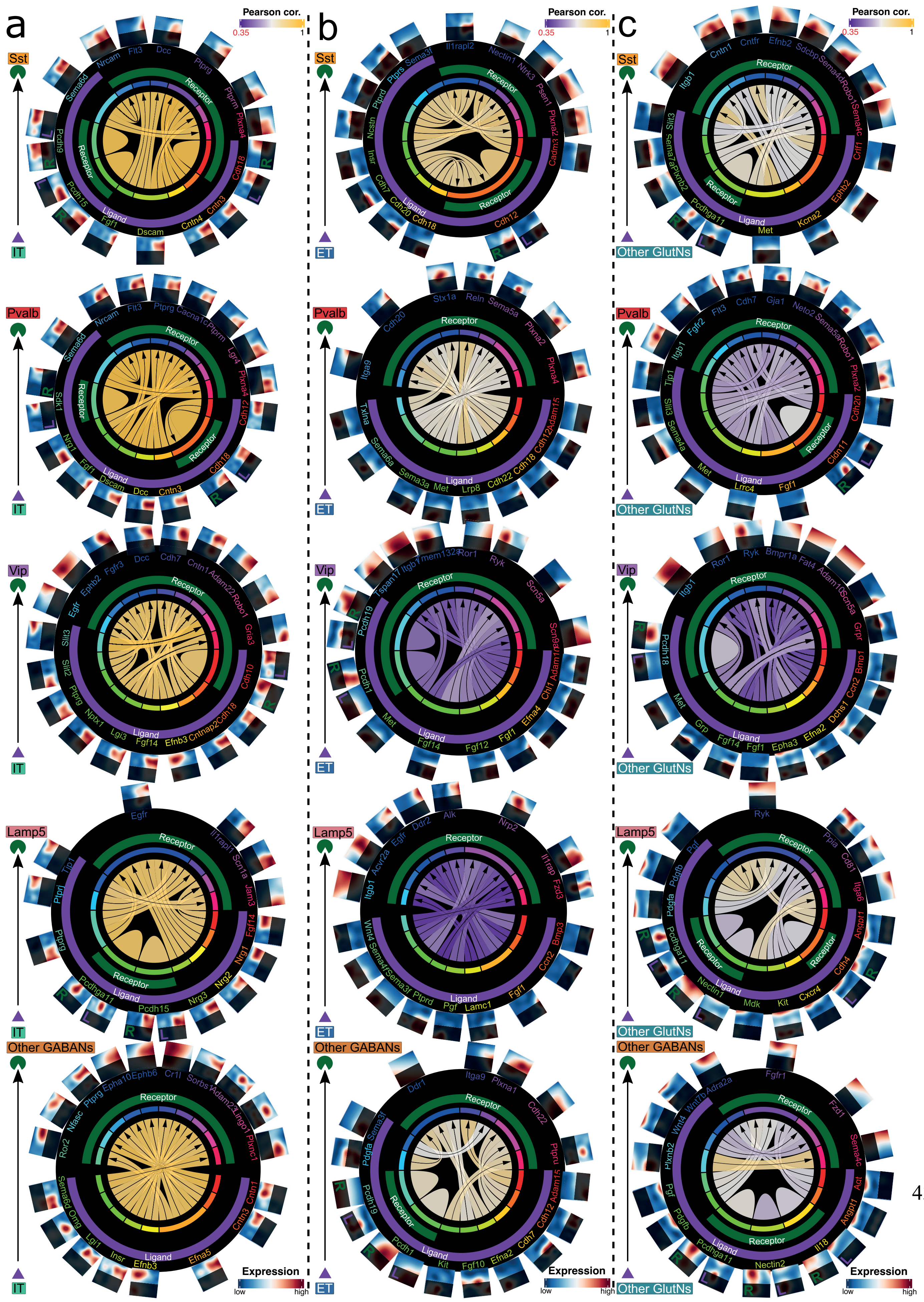

a
b
c
Pearson cor.
0.35
1
Sst
Pvalb
Vip
Lamp5
Other GABANs
IT
ET
Other GlutNs
Receptor
Ligand
R
L
Expression
low
high

**Figure S15. ”Upper layers” transcriptional landscapes correlation of ligand-receptor pairs between glutamatergic expressing ligand and GABAergic expressing receptor families.** (a) Chord diagram representing the correlation between transcriptional landscapes of IT expressing ligand and GABAN families expressing receptor. (b) Chord diagram representing the correlation between transcriptional landscapes of ET expressing ligand and GABAN families expressing receptor. (c) Chord diagram representing the correlation between transcriptional landscapes of Other GlutNs expressing ligand and GABAN families expressing receptor. Only the top 10 most correlated LR pairs are represented between each family couple. Pearson correlation values were calculated based on the non-shaded area of transcriptional landscapes (Pseudo-layer values from 0 to 0.5 were used).

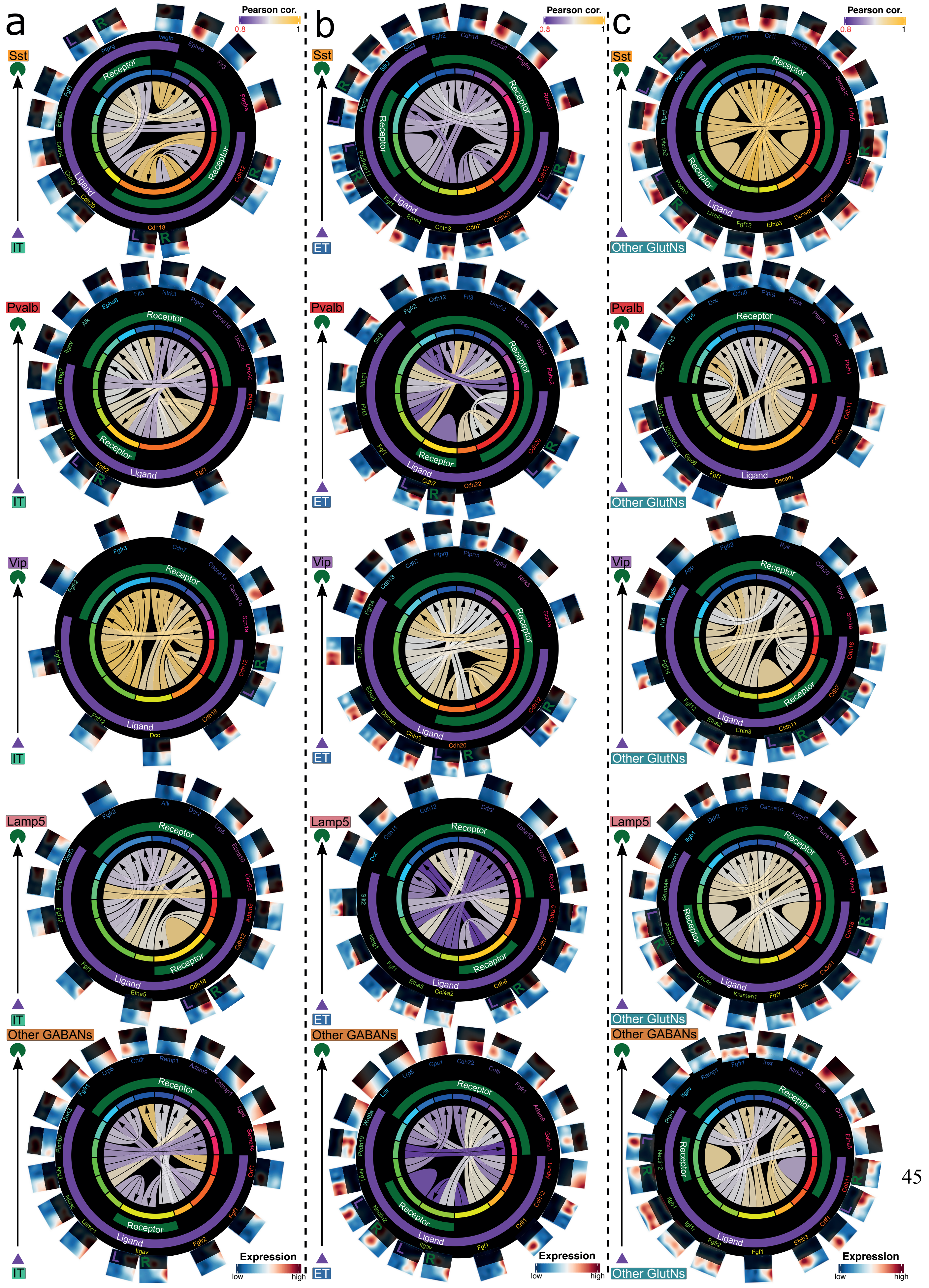
a
b
c
Pearson cor.
0.8
1
Sst
Pvalb
Vip
Lamp5
Other GABANs
IT
ET
Other GlutNs
Receptor
Ligand
Expression
low
high

**Figure S16. ”Deep layers” transcriptional landscapes correlation of ligand-receptor pairs between glutamatergic expressing ligand and GABAergic expressing receptor families.** (a) Chord diagram representing the correlation between transcriptional landscapes of IT expressing ligand and GABAN families expressing receptor. (b) Chord diagram representing the correlation between transcriptional landscapes of ET expressing ligand and GABAN families expressing receptor. (c) Chord diagram representing the correlation between transcriptional landscapes of Other GlutNs expressing ligand and GABAN families expressing receptor. Only the top 10 most correlated LR pairs are represented between each family couple. Pearson correlation values were calculated based on the non-shaded area of transcriptional landscapes (Pseudo-layer values from 0.5 to 1 were used).

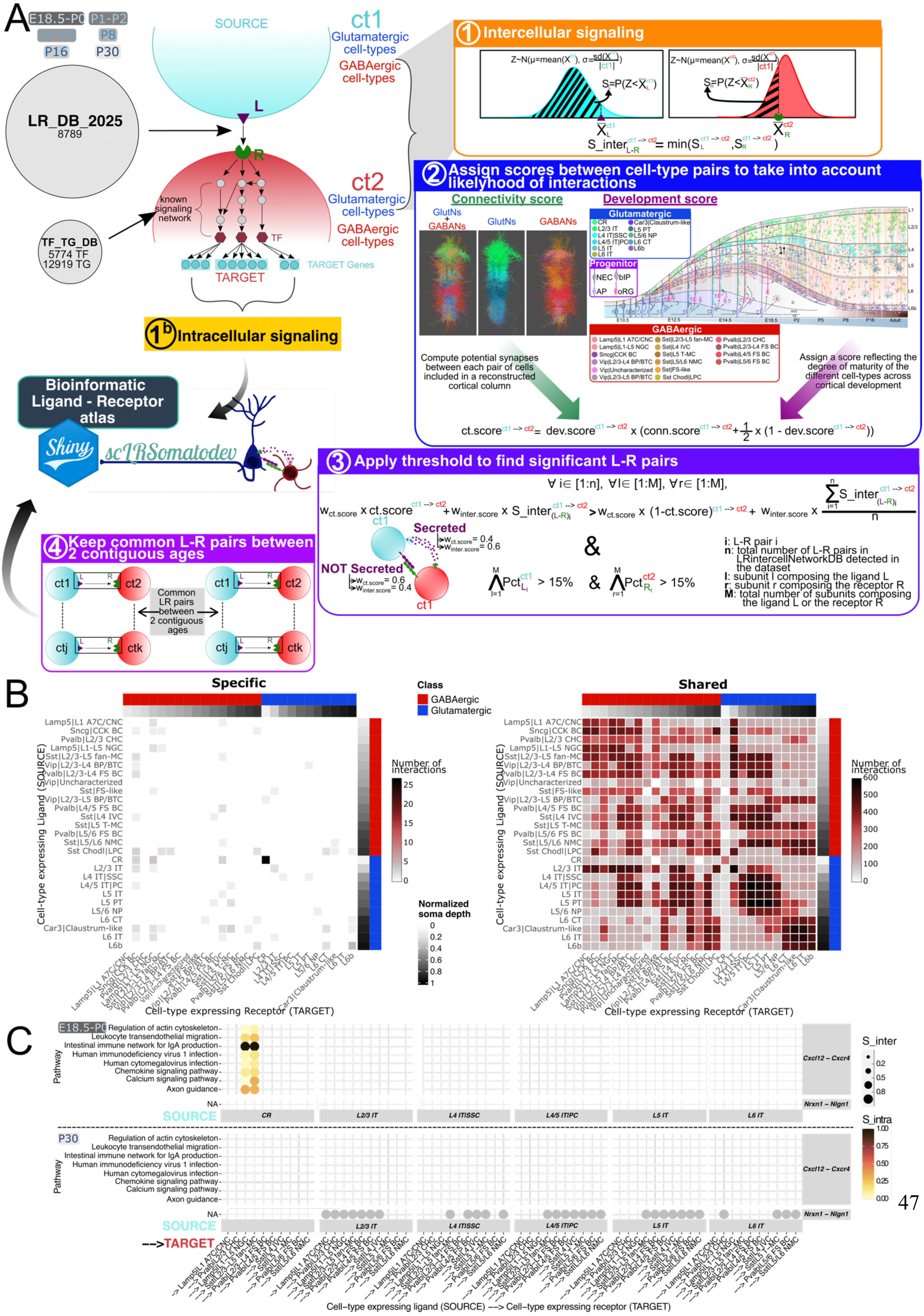
A
E18.5-P0
P1-P2
P8
P16
P30
LR_DB_2025
8789
TF_TG_DB
5774 TF
12919 TG
SOURCE
ct1
Glutamatergic cell-types
GABAergic cell-types
L
R
known signaling network
TF
ct2
Glutamatergic cell-types
GABAergic cell-types
TARGET
TARGET Genes
1b Intracellular signaling
Bioinformatic Ligand - Receptor atlas
Shiny
scLRSomatodev
1 Intercellular signaling
S_inter = min(S_L, S_R)
2 Assign scores between cell-type pairs to take into account likelyhood of interactions
Connectivity score
Development score
GlutNs + GABANs
GlutNs
GABANs
Glutamatergic
Progenitor
GABAergic
Compute potential synapses between each pair of cells included in a reconstructed cortical column
Assign a score reflecting the degree of maturity of the different cell-types across cortical development
ct.score = dev.score x (conn.score + 1/2 x (1 - dev.score))
3 Apply threshold to find significant L-R pairs
Secreted
NOT Secreted
i: L-R pair i
n: total number of L-R pairs in LRintercellNetworkDB detected in the dataset
l: subunit l composing the ligand L
r: subunit r composing the receptor R
M: total number of subunits composing the ligand L or the receptor R
4 Keep common L-R pairs between 2 contiguous ages
Common LR pairs between 2 contiguous ages
B
Specific
Shared
Class
GABAergic
Glutamatergic
Number of interactions
Normalized soma depth
Cell-type expressing Ligand (SOURCE)
Cell-type expressing Receptor (TARGET)
C
Pathway
Regulation of actin cytoskeleton
Leukocyte transendothelial migration
Intestinal immune network for IgA production
Human immunodeficiency virus 1 infection
Human cytomegalovirus infection
Chemokine signaling pathway
Calcium signaling pathway
Axon guidance
Cxcl12 – Cxcr4
Nrxn1 – Nlgn1
SOURCE
CR
L2/3 IT
L4 IT|SSC
L4/5 IT|PC
L5 IT
L6 IT
S_inter
S_intra
--->TARGET
Cell-type expressing ligand (SOURCE) ---> Cell-type expressing receptor (TARGET)

**Fig. S17. Establishment of a ligand-receptor atlas between glutamatergic and GABAergic cell-types.** (A) scSeqComm detailed pipeline. A Shiny app called scLRSomatoDev was constructed to allow the neuroscience community to view the ligand-receptor atlas and to follow gene expression throughout the somatosensory cortex development. (B) Specific and share interactions between all cell pairs that are contiguous between at least 2 stages. (C) Example dot plot for inter and intracellular communications.

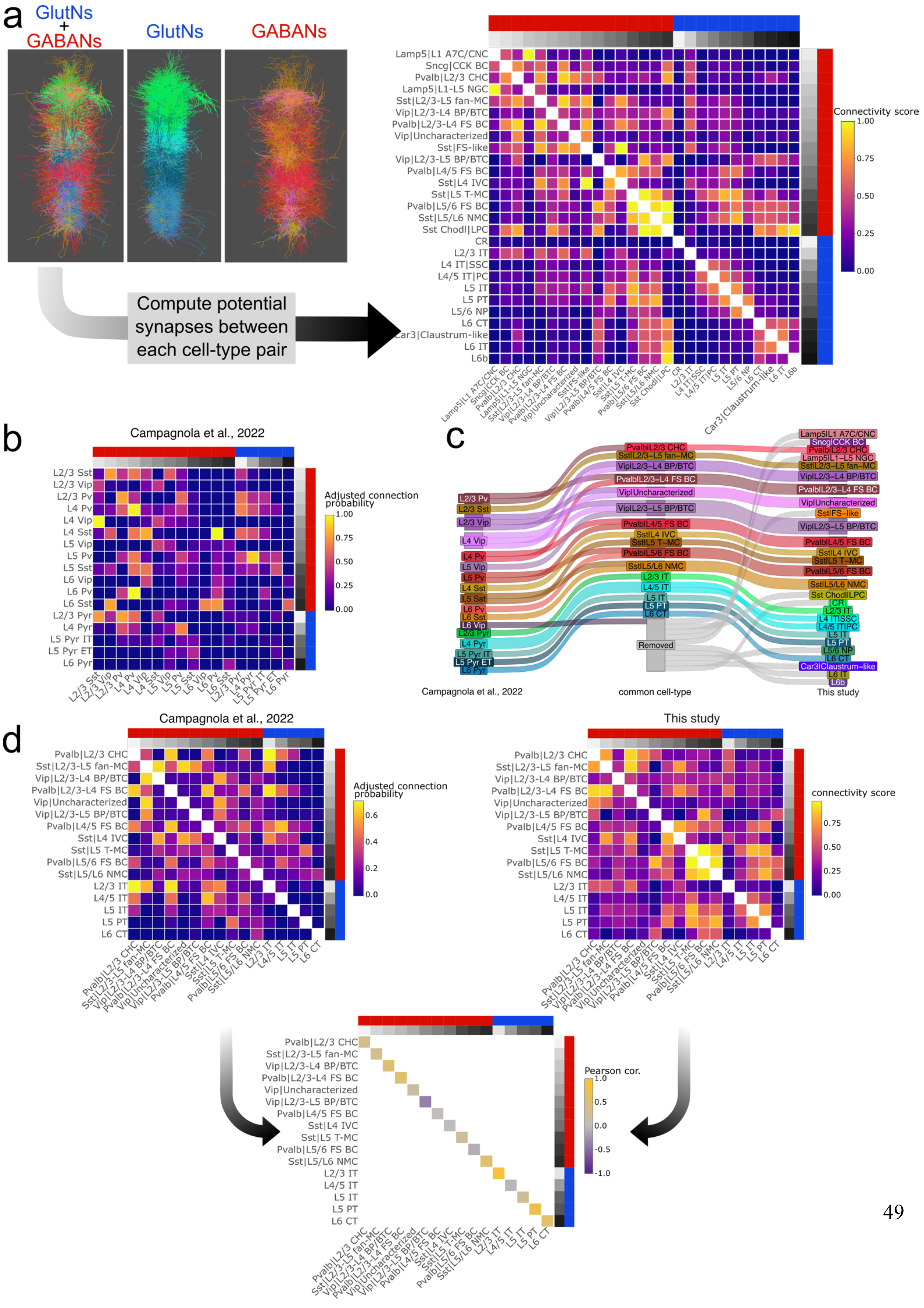
a
GlutNs + GABANs
GlutNs
GABANs
Compute potential synapses between each cell-type pair
Connectivity score
Lamp5|L1 A7C/CNC
Sncg|CCK BC
Pvalb|L2/3 CHC
Lamp5|L1-L5 NGC
Sst|L2/3-L5 fan-MC
Vip|L2/3-L4 BP/BTC
Pvalb|L2/3-L4 FS BC
Vip|Uncharacterized
Sst|FS-like
Vip|L2/3-L5 BP/BTC
Pvalb|L4/5 FS BC
Sst|L4 IVC
Sst|L5 T-MC
Pvalb|L5/6 FS BC
Sst|L5/L6 NMC
Sst Chodl|LPC
CR
L2/3 IT
L4 IT|SSC
L4/5 IT|PC
L5 IT
L5 PT
L5/6 NP
L6 CT
Car3|Claustrum-like
L6 IT
L6b
b
Campagnola et al., 2022
Adjusted connection probability
L2/3 Sst
L2/3 Vip
L2/3 Pv
L4 Pv
L4 Vip
L4 Sst
L5 Vip
L5 Pv
L5 Sst
L6 Vip
L6 Pv
L6 Sst
L2/3 Pyr
L4 Pyr
L5 Pyr IT
L5 Pyr ET
L6 Pyr
c
Removed
Campagnola et al., 2022
common cell-type
This study
d
Campagnola et al., 2022
This study
Adjusted connection probability
connectivity score
Pearson cor.

**Figure S18. Comparison between the connectivity score obtained from the reconstructed cortical column and the connection probability from Campagnola et al., 2022.** (a) Connectivity score between cell-type pairs obtained by computing potential synapses between each cell pairs from the reconstructed cortical column. (b) Adjusted connection probability between each cell-type pairs from Campagnola et al.,2022. (c) Sankey plot representing the common nomenclature used to compare cell-types from Campagnola et al., 2022 and this study. (d) Correlation between each common cell-type from Campagnola et al., 2022 and this study. As the adjusted connection probability between each cell-type pair was directed in Campagnola et al, 2022, the mean of the 2 directions was taken to make the comparison between Campagnola et al, 2022 and this study.

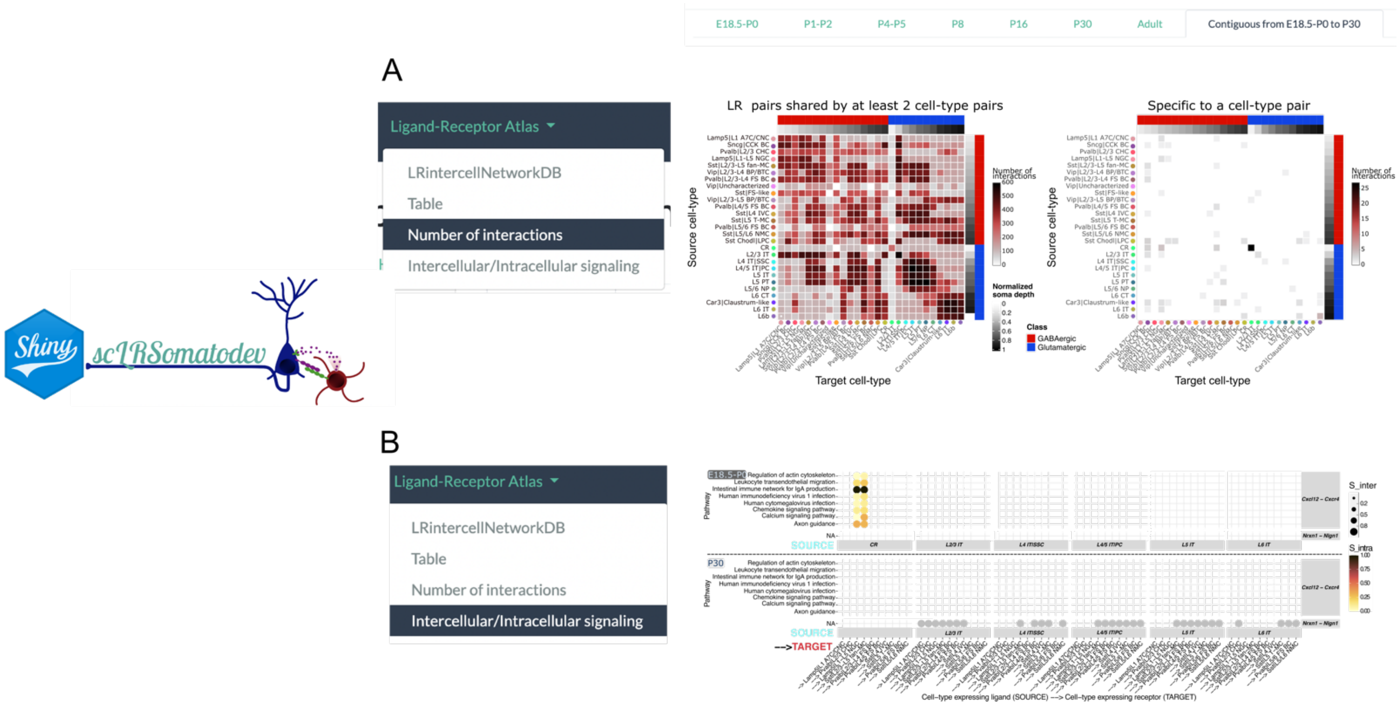


**Figure S19. LR prediction panel of the Shiny app scLRSomatodev. Top, function Number of interactions. Bottom, function Intercellular/Intracellular signaling.** (A) Heatmaps representing the number of interactions between all cell-types pairs that are specific to a cell-type pair (left) or shared by at least 2 cell-type pairs (right) and that are common between 2 contiguous ages. The blue color represents glutamatergic cell-types and the red color the GABAergic cell-types. The white to black color gradient represents the median of the normalized soma depth of each cell-type. 0 representing the pial surface and 1 the white matter. (B) Examples of intercellular (S_inter) and intracellular (S_intra) signaling for two ligand-receptor pairs (Cxcl12 - Cxcr4 and Nrxn1 - Nlgn1) between IT family, CR cell-type and Sst, Pvalb and Lamp5 familes at E18.5-P0 and P30. Pathways are shown on the left. LR pairs are represented on the right. Grey circles: LR pairs having no S_intra score values (i.e., LR pairs having no annotated receptor downstream transcription factors).

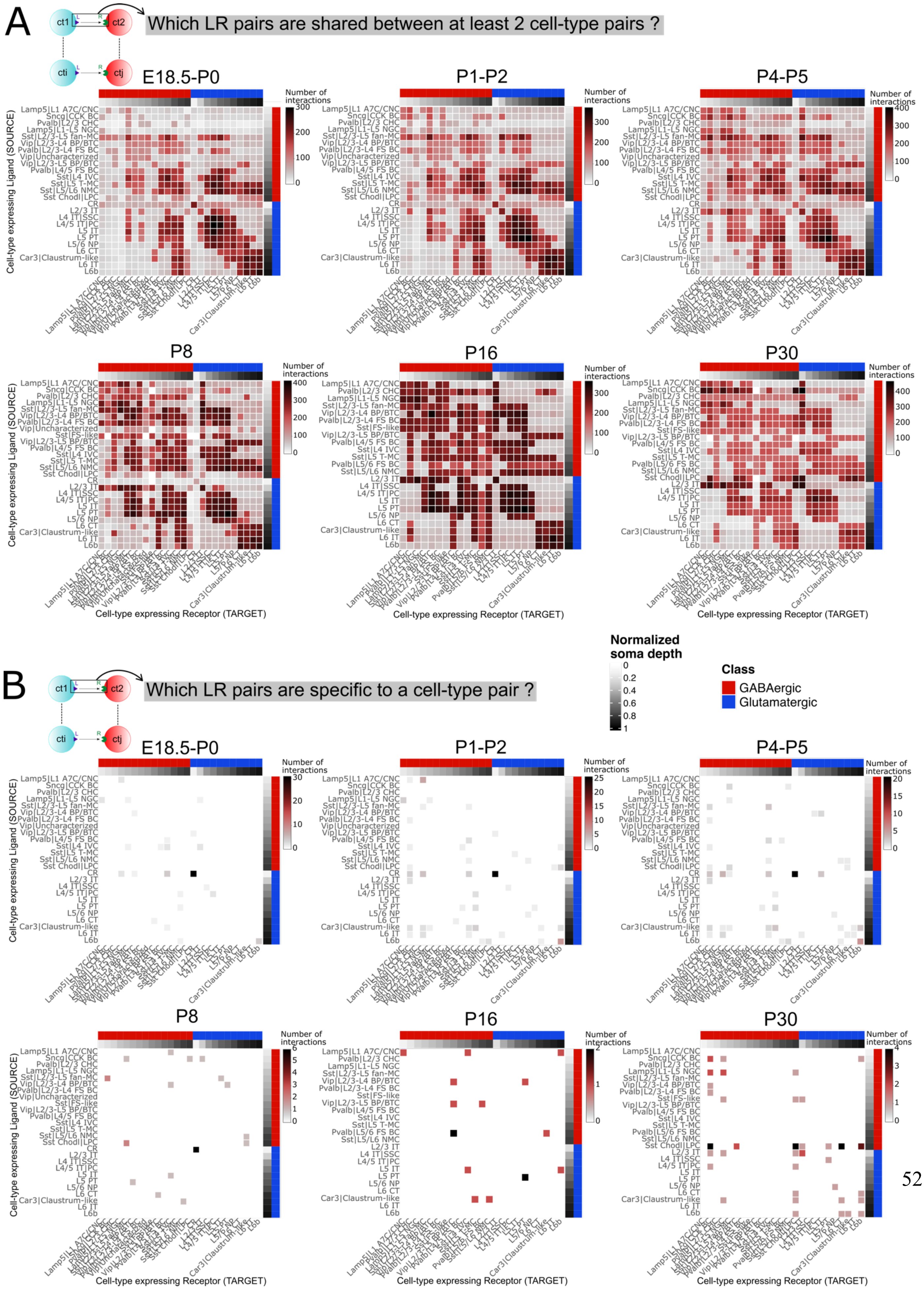
A
ct1
ct2
cti
ctj
Which LR pairs are shared between at least 2 cell-type pairs ?
E18.5-P0
P1-P2
P4-P5
P8
P16
P30
Number of interactions
Cell-type expressing Ligand (SOURCE)
Cell-type expressing Receptor (TARGET)
B
Which LR pairs are specific to a cell-type pair ?
Normalized soma depth
Class
GABAergic
Glutamatergic
Lamp5|L1 A7C/CNC
Sncg|CCK BC
Pvalb|L2/3 CHC
Lamp5|L1-L5 NGC
Sst|L2/3-L5 fan-MC
Vip|L2/3-L4 BP/BTC
Pvalb|L2/3-L4 FS BC
Vip|Uncharacterized
Vip|L2/3-L5 BP/BTC
Pvalb|L4/5 FS BC
Sst|L4 IVC
Sst|L5 T-MC
Sst|L5/L6 NMC
Sst Chodl|LPC
CR
L2/3 IT
L4 IT|SSC
L4/5 IT|PC
L5 IT
L5 PT
L5/6 NP
L6 CT
Car3|Claustrum-like
L6 IT
L6b
Sst|FS-like
Pvalb|L5/6 FS BC

**Figure S20. Number of shared and specific inferred LR interactions between cell-type pairs from E18.5-P0 to P30. (A)** Heatmap representing the number of interactions between all cell-type pairs that are at least shared by 2 cell-type pairs. **(B)** Heatmap representing the number of interactions between all cell-types pairs that are specific to a cell-type pair (cell-type expressing ligand). The blue color represents glutamatergic cell-types and the red color the GABAergic cell-types. The white to black color gradient represents the median of the normalized soma depth of each cell-type. 0 representing the pial surface and 1 the white matter.

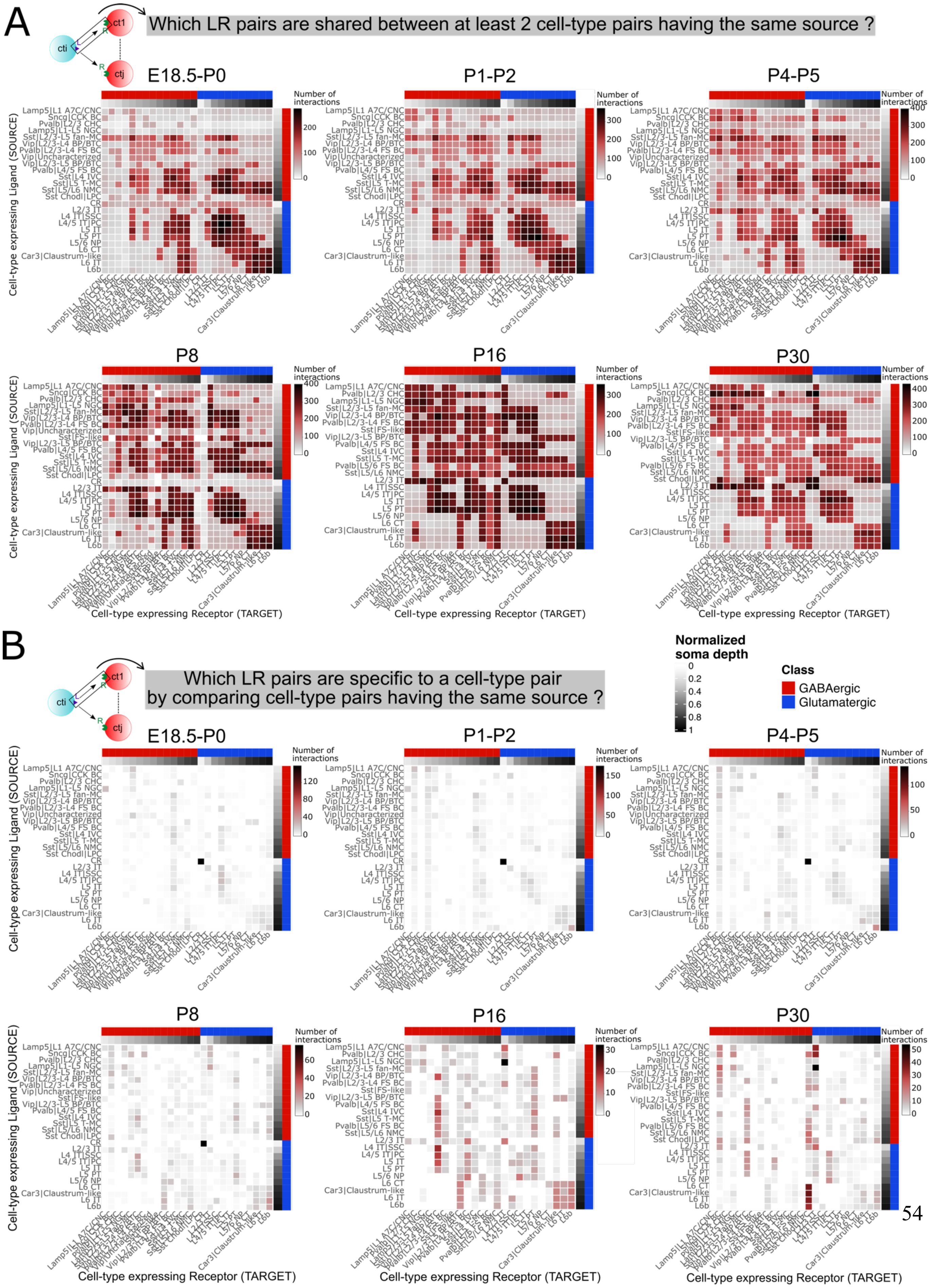

A
Which LR pairs are shared between at least 2 cell-type pairs having the same source ?
cti
ct1
ctj
E18.5-P0
P1-P2
P4-P5
P8
P16
P30
Number of interactions
Cell-type expressing Ligand (SOURCE)
Cell-type expressing Receptor (TARGET)
B
Which LR pairs are specific to a cell-type pair by comparing cell-type pairs having the same source ?
Normalized soma depth
Class
GABAergic
Glutamatergic

**Figure S21. Number of shared and specific inferred LR interactions between cell-type pairs having the same source from E18.5-P0 to P30.** (A) Heatmap representing the number of interactions between all cell-type pairs that are at least shared by 2 cell-type pairs having the same source. (B) Heatmap representing the number of interactions between all cell-type pairs that are specific to a cell-type pair by comparing cell- type pairs having the same source (cell-type expressing ligand). The blue color represents glutamatergic cell-types and the red color the GABAergic cell-types. The white to black color gradient represents the median of the normalized soma depth of each cell-type. 0 representing the pial surface and 1 the white matter.

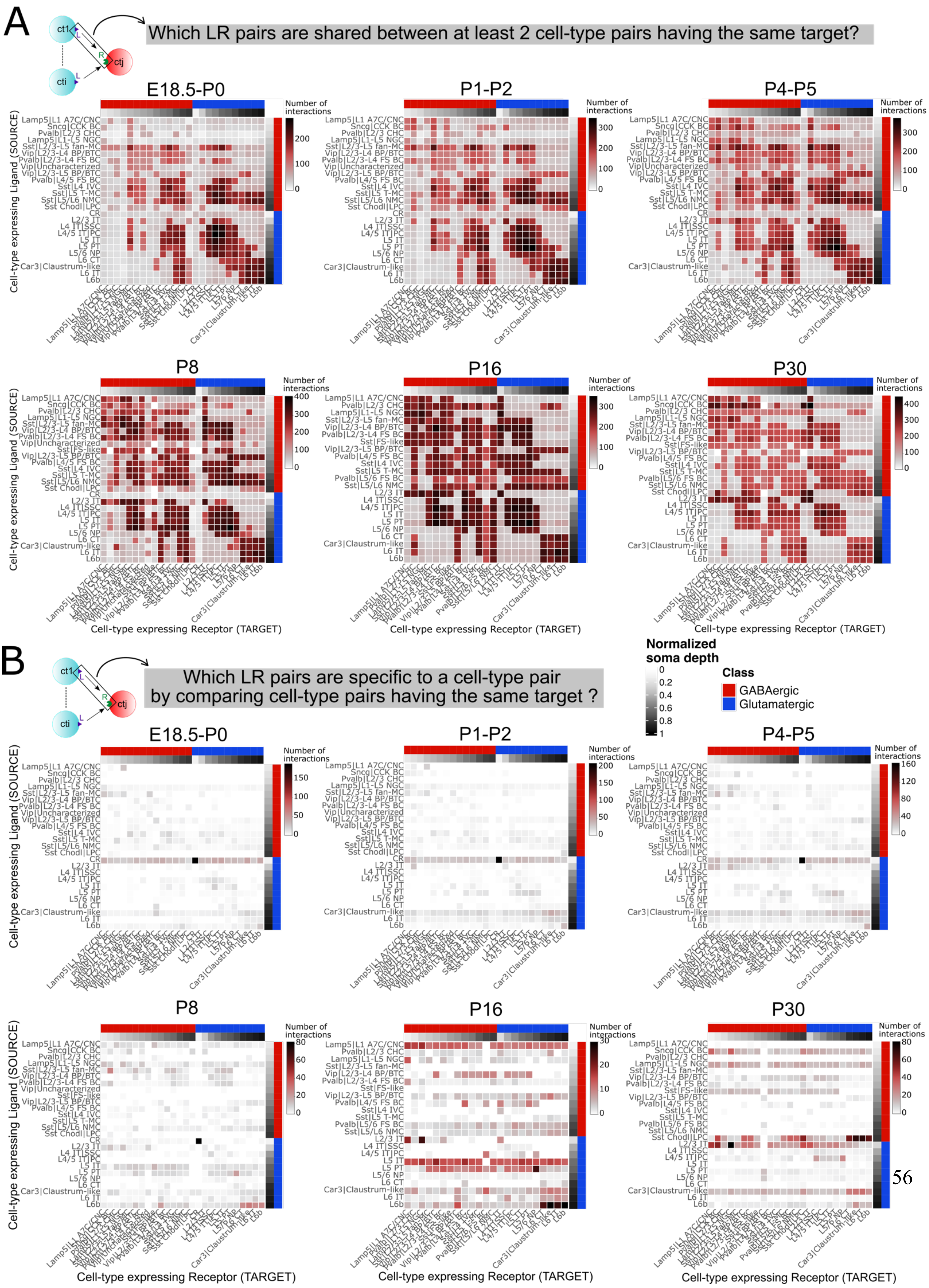
A
Which LR pairs are shared between at least 2 cell-type pairs having the same target?
ct1
ctj
cti
E18.5-P0
P1-P2
P4-P5
P8
P16
P30
Number of interactions
Cell-type expressing Ligand (SOURCE)
Cell-type expressing Receptor (TARGET)
B
Which LR pairs are specific to a cell-type pair by comparing cell-type pairs having the same target ?
Normalized soma depth
Class
GABAergic
Glutamatergic

**Figure S22. Number of shared and specific interactions between cell-type pairs having the same target from E18.5-P0 to P30.** (A) Heatmap representing the number of interactions between all cell-type pairs that are at least shared by 2 cell-type pairs having the same target. (B) Heatmap representing the number of interactions between all cell-type pairs that are specific to a cell-type pair by comparing cell-type pairs having the same target (cell-type expressing receptor).The blue color represents glutamatergic cell-types and the red color the GABAergic cell-types. The white to black color gradient represents the median of the normalized soma depth of each cell-type. 0 representing the pial surface and 1 the white matter.

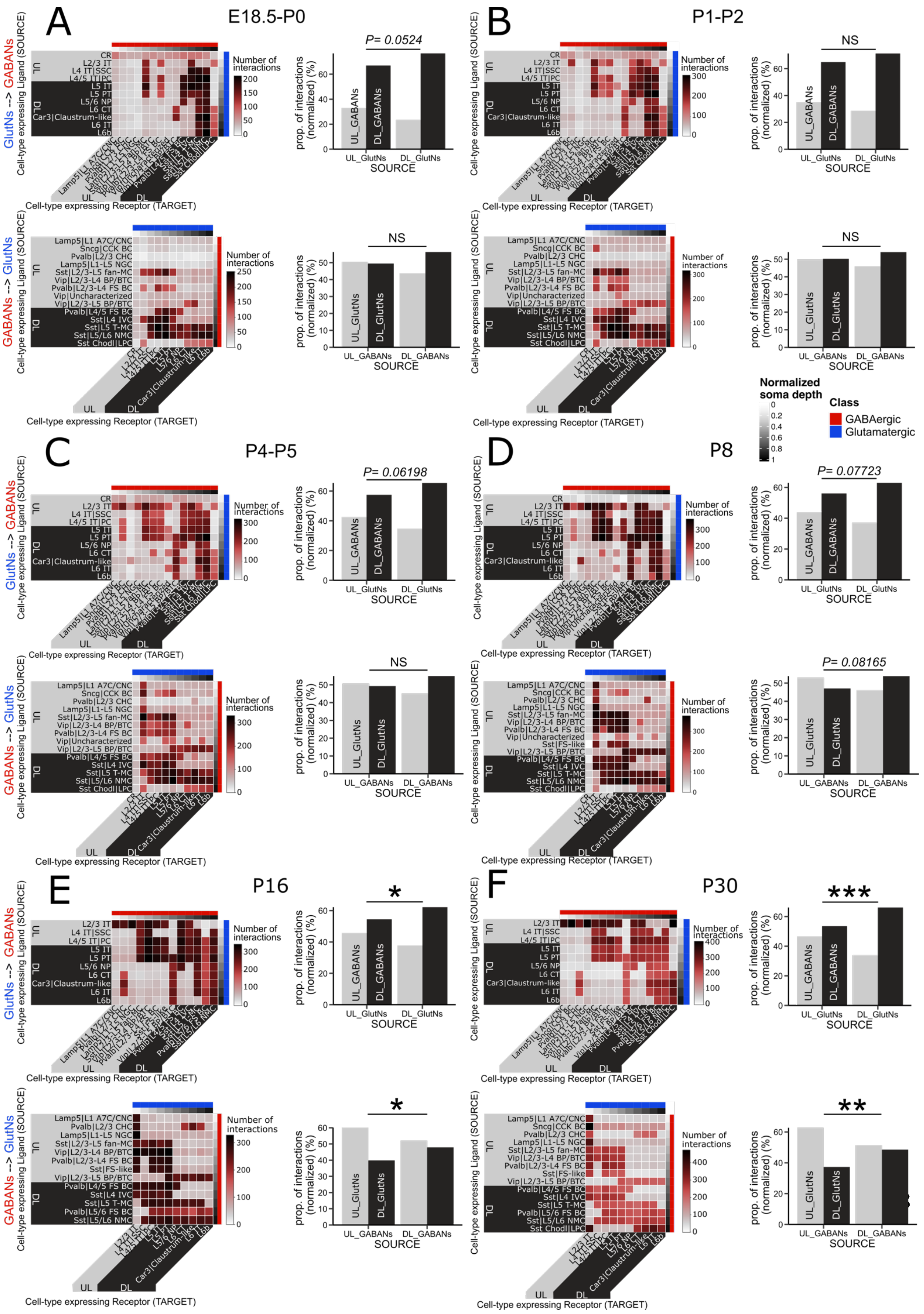
A
E18.5-P0
P= 0.0524
NS
B
P1-P2
NS
NS
C
P4-P5
P= 0.06198
NS
D
P8
P= 0.07723
P= 0.08165
E
P16
*
*
F
P30
***
**
GlutNs --> GABANs
GABANs --> GlutNs
Cell-type expressing Ligand (SOURCE)
Cell-type expressing Receptor (TARGET)
Number of interactions
prop. of interactions (normalized) (%)
SOURCE
UL_GlutNs
DL_GlutNs
UL_GABANs
DL_GABANs
UL
DL
Normalized soma depth
Class
GABAergic
Glutamatergic

**Figure S23. Number of interactions between GlutNs SOURCE (or TARGET) cell-types and GABANs TARGET (or SOURCE) cell-types.** Top: heatmap representing the number of interactions between GlutNs expressing ligand (SOURCE) and GABANs expressing receptor (TARGET) along with the total percentage of interactions (normalized) between the UL GlutNs and the UL and DL GABANs versus between the DL GlutNs and the UL and DL GABANs at E18.5-P0 **(A)**, P1-P2 **(B)**, P4-P5 **(C)**, P8 **D)**, P16 **(E)** and P30 **(F)**. Bottom: heatmap representing the number of interactions between GABANs expressing ligand (SOURCE) and GlutNs expressing receptors (TARGET) along with the total percentage of interactions (normalized) between the UL GABANs and the UL and DL GlutNs versus between the DL GABANs and the UL and DL GlutNs at E18.5-P0 **(A)**, P1-P2 **(B)**, P4-P5 **(C)**, P8 **(D)**, P16 **(E)** and P30 **(F)**. The number of interactions was normalized by dividing by the number of cell-type couples involved before achieving the $\chi^2$ test so that the comparison does not suffer from the unbalanced number of cell-types between UL and DL. *p<0.05, **p<0.01, ***p<0.001, $\chi^2$ test.

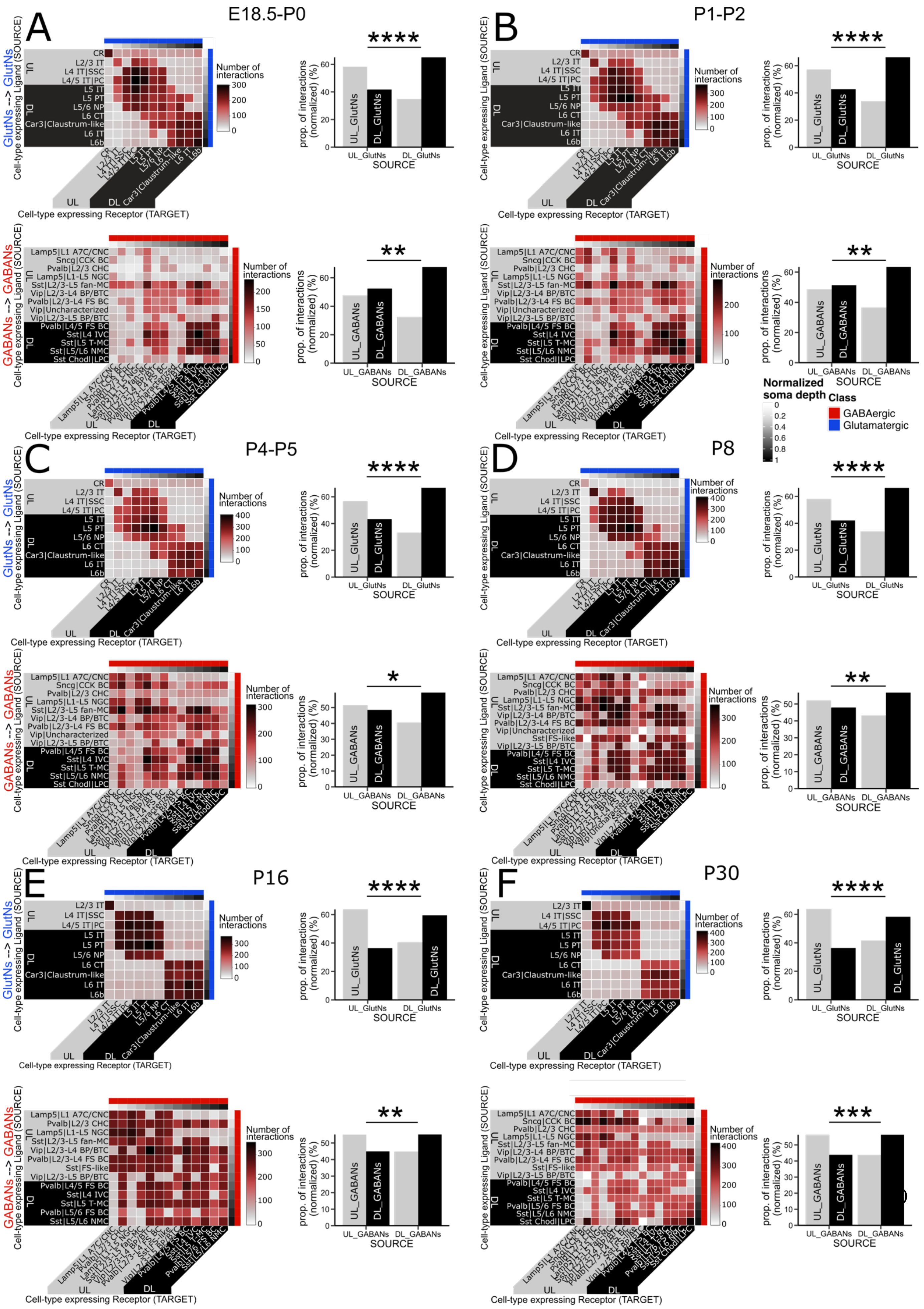

A
E18.5-P0
B
P1-P2
C
P4-P5
D
P8
E
P16
F
P30
GlutNs --> GlutNs
GABANs --> GABANs
Cell-type expressing Ligand (SOURCE)
Cell-type expressing Receptor (TARGET)
Number of interactions
UL
DL
CR
L2/3 IT
L4 IT|SSC
L4/5 IT|PC
L5 IT
L5 PT
L5/6 NP
L6 CT
Car3|Claustrum-like
L6 IT
L6b
Lamp5|L1 A7C/CNC
Sncg|CCK BC
Pvalb|L2/3 CHC
Lamp5|L1-L5 NGC
Sst|L2/3-L5 fan-MC
Vip|L2/3-L4 BP/BTC
Pvalb|L2/3-L4 FS BC
Vip|Uncharacterized
Vip|L2/3-L5 BP/BTC
Pvalb|L4/5 FS BC
Sst|L4 IVC
Sst|L5 T-MC
Sst|L5/L6 NMC
Sst Chodl|LPC
Sst|FS-like
Pvalb|L5/6 FS BC
****
***
**
*
prop. of interactions (normalized) (%)
UL_GlutNs
DL_GlutNs
UL_GABANs
DL_GABANs
SOURCE
Normalized soma depth
Class
GABAergic
Glutamatergic

**Figure S24. Number of interactions between cell-types of the same class.** Top: heatmap representing the number of interactions between GlutNs expressing ligand (SOURCE) and GlutNs expressing receptor (TARGET) along with the total percentage of interactions (normalized) between the UL GlutNs and the UL and DL GlutNs versus between the DL GlutNs and the UL and DL GlutNs at E18.5-P0 **(A)**, P1-P2 **(B)**, P4-P5 **(C)**, P8 **(D)**, P16 **(E)** and P30 **(F)**. Bottom: heatmap representing the number of interactions between GABANs expressing ligand (SOURCE) and GABANs expressing receptors (TARGET) along with the total percentage of interactions (normalized) between the UL GABANs and the UL and DL GABANs versus between the DL GABANs and the UL and DL GABANs at E18.5-P0 **(A)**, P1-P2 **(B)**, P4-P5 **(C)**, P8 **(D)**, P16 **(E)** and P30 **(F)**. The number of interactions was normalized by dividing by the number of cell-type couples involved before achieving the $\chi^2$ test so that the comparison does not suffer from the unbalanced number of cell-types between UL and DL. *p<0.05, **p<0.01, ***p<0.001, ****p<0.0001, $\chi^2$ test

**Table S1. The LR database LR_DB_2025.**

**Table S2. Categories of LRs belonging to 5 main neurodevelopmental processes**

**Table S3. Development scores**

**Table S4. Connectivity scores**

**Table S5. Categories of LRs belonging to neurodevelopmental diseases**

**Data S1. (separate file)**

Type or paste caption here.

10. Fischer, S. & Gillis, J. How many markers are needed to robustly determine a cell's type?

11. Gouwens, N. W. *et al.* Integrated Morphoelectric and Transcriptomic Classification of


`

Cortical GABAergic Cells. *Cell* **183**, 935-953.e19 (2020).

21. Cabello-Aguilar, S., Fau, C., Lacroix, M. & Colinge, J. SingleCellSignalR: inference of intercellular networks from single-cell transcriptomics.

28. Baruzzo, G., Cesaro, G. & Di Camillo, B. Identify, quantify and characterize cellular

communication from single-cell RNA sequencing data with *scSeqComm*. *Bioinformatics* **38**, 1920–1929 (2022).

`
